# Fluorogenic Covalent Probes for RNA

**DOI:** 10.1101/2025.08.29.673142

**Authors:** Jinwoo Shin, Moon Jung Kim, Eric T. Kool

## Abstract

Sequence-generalized fluorescent labels and stains for RNA can enable imaging, tracking and analysis of the biopolymer. However, current non-covalent RNA dyes are poorly selective for RNA over DNA, interact weakly with their target, and can show limited utility in cellular RNA staining due to poor selectivity and high background signals. Here we report a fluorogenic covalent labeling approach based on acylimidazole-mediated reaction of donor-acceptor fluorophores with 2′-hydroxyl (2′-OH) groups of RNA, providing a wavelength-tunable, sequence-independent strategy for selective labeling of the biopolymer. This reactive probe design enables labeling and imaging under mild aqueous conditions, providing up to 390-fold fluorescence enhancement and 970-fold selectivity for RNA over DNA, with four emission colors documented. The covalent fluorophore platform enables improved new tools for RNA-specific analysis and imaging in gels, in solution, and in living cells.

## INTRODUCTION

Ribonucleic acids (RNAs) play complex and central roles in cellular regulation, catalysis, and signaling^1-4^. Visualizing RNA during separation and analysis, as well as in its native cellular context, is an essential tool for analyzing its functions. The ability to image RNA populations can enable the monitoring of cellular transcription, track the mobility and dynamics of varied classes of transcripts, and analyze RNA-rich subcellular compartments and bioaggregates such as occur in nucleoli, P bodies, and stress granules^5-7^. However, despite decades of efforts, imaging the populations of RNAs *in vitro* and in cells and tissues remains a fundamental challenge in the chemistry of biomolecular visualization^8^.

Most recent efforts aimed at the development of RNA labeling and imaging strategies have focused primarily on specific transcripts rather than generalized imaging. Significant developments include aptamer-based systems, such as the Spinach and Mango aptamers, which employ engineered RNA motifs that bind dyes, resulting in a robust increase in emission^9-12^. While useful for specific engineered transcripts, the aptamer modules are not applicable to the native cellular population of RNAs. Molecular beacons, which are short deoxyribonucleic acid (DNA) hairpin oligonucleotides labeled with a fluorophore and quencher pair, constitute a second major strategy for RNA detection and imaging^13^. These modified DNAs fluoresce upon hybridization to target RNA, directed by sequence complementarity. Molecular beacons have found utility in applications such as real-time polymerase chain reaction (PCR), but are sequence-specific rather than general, and can show substantial background emission in intracellular application^8^. Other hybridization probes, including locked nucleic acid (LNA)^14^ and peptide nucleic acid (PNA)-based designs^15^, offer improved affinity and nuclease resistance but still rely on sequence-specific targeting and are often restricted to fixed-cell imaging^16, 17^. Enzyme-assisted strategies for RNA labeling, such as tRNA guanine transglycosylase (TGT)-mediated labeling^18^, DNAzyme labeling^19^, or CRISPR–Cas13-directed tagging^20^, provide covalent or modular labeling approaches for RNA but require RNA sequence engineering or are targeted to specific transcripts, and thus are not applicable to general RNA populations^21^.

Recently, bioorthogonal metabolic incorporation of nucleotides has enabled transcriptome-wide RNA labeling^22, 23^, and thus is not restricted to specific sequences. In this approach, modified nucleosides are taken up into cells, are phosphorylated by kinases and subsequently incorporated into new RNAs by extant RNA polymerase enzymes. Examples include 5-ethynyluridine (5EU)^24^ and N^6^-propargyladenosine^25^, which contain alkyne groups for further functionalization by click chemistry. Cells are then fixed to enable fluorescent tagging and washing away of excess dyes. Due to these required steps, the method is not generally applicable for imaging RNAs in live cells and is not useful for gel-based visualization and imaging of native RNAs.

There remains a pressing need for labeling strategies that enable direct, sequence-independent, yet highly selective labeling of RNA molecules that relies on intrinsic chemical features rather than sequence. Toward this end, small-molecule fluorescent dyes that associate non-covalently with RNA have come under development for general imaging of the biopolymer, leveraging their structural tunability^26^ (Figure 1a and Figure S1). These aromatic cation-based push-pull fluorophores associate with RNA non-covalently via electrostatic attraction and intercalation. They can be engineered to emit fluorescence over a broad range of wavelengths and are in some cases cell-permeable, making some examples suitable for real-time imaging as well as gel staining. Among the earliest dyes employed for RNA imaging in gels is the intercalator ethidium bromide (EtBr), which stains DNA preferentially but also yields RNA staining as well^27^. More recently, commercial fluorescent dyes such as the SYBR family (*e*.*g*., SYBR Green and SYBR Gold) have been developed and widely employed. These dyes bind to nucleic acids by electrostatics, intercalation or groove binding and offer improved sensitivity.

**Figure 1.**
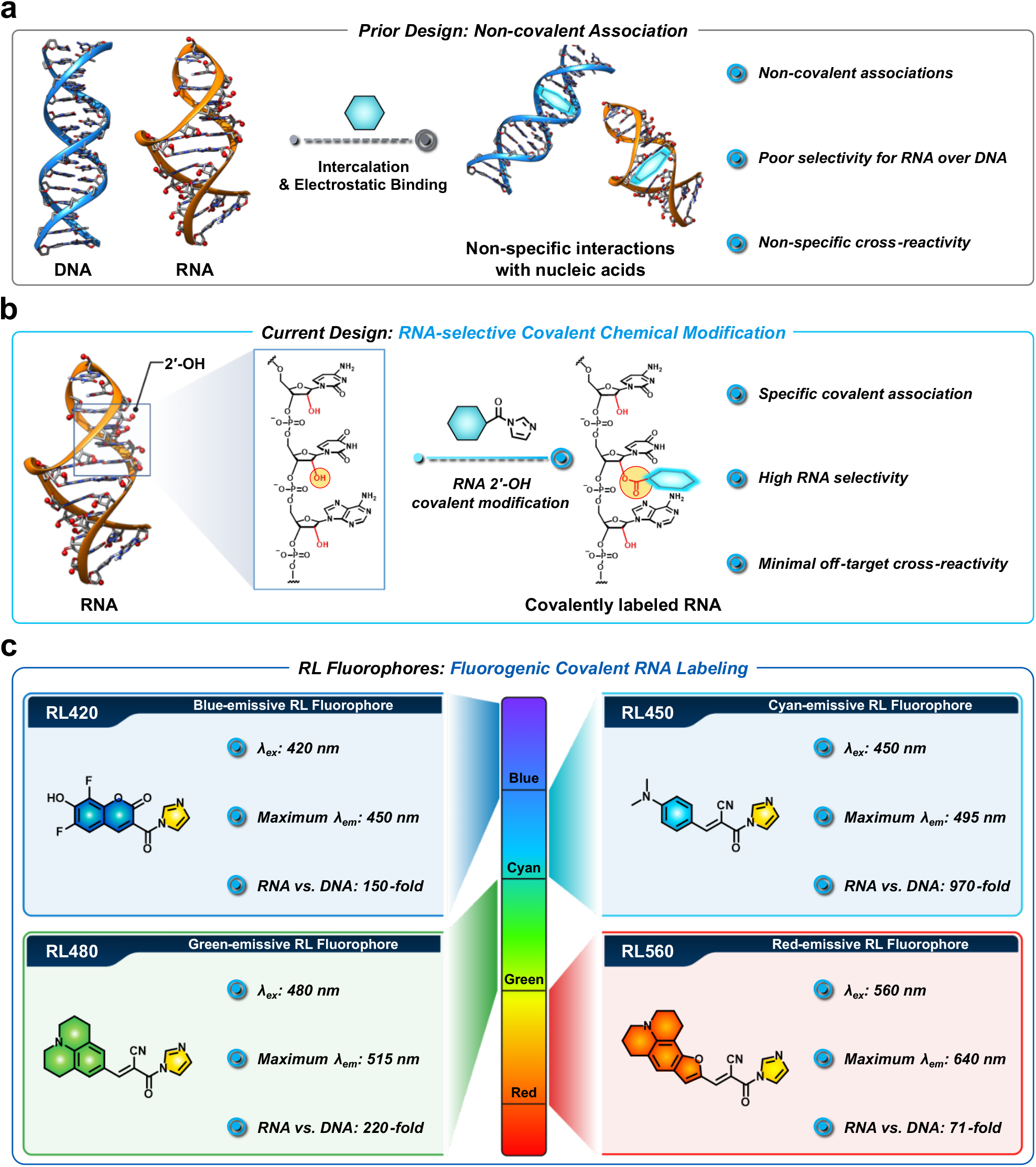
Development of RNA-selective fluorogenic covalent acylating probes (RiboLight, RL). (a) Traditional small-molecule RNA fluorophores primarily rely on non-covalent association, resulting in limited selectivity and susceptibility to background signals from DNA, mitochondria or proteins. (b) Current design strategy leverages the uniquely reactive 2′-OH group of RNA to achieve covalent chemical modification, enabling selective RNA labeling. (c) Structures and spectral properties of four wavelength-engineered **RL** fluorophores (**RL420, RL450, RL480** and **RL560**). Each exhibits distinct excitation and emission maxima with high fluorescence turn-on ratios for RNA over DNA (71-to 970-fold), enabling multicolor, wash-free, and RNA-selective labeling.

SYBR Green II stains both RNA and single-stranded DNA (ssDNA), while SYBR Gold exhibits a preference for double-stranded DNA (dsDNA), but can also stain ssDNA and RNA to a lesser extent^28^. Also employed is SYTO RNASelect, a commercial dye designed to selectively stain RNA in live cells. Recent data suggest that this dye has limited cell permeability and has reportedly low (1.6-fold) selectivity for RNA over DNA^29^. In addition to these, several other fluorophore designs have been tested, aiming for improvements in sensitivity, selectivity and imaging performance. QUID-2, a styryl-based bis-cationic dye, was reported to exhibit superior photostability and biocompatibility compared to SYTO RNASelect, enabling RNA imaging in live cells^29^. However, the authors incorporated a washing step prior to imaging. Push–pull cationic dyes incorporating methylpyridinium or methylquinolinium moieties have been reported to have over 100-fold fluorescence enhancement upon RNA binding and stain nucleolar and mitochondrial RNA^26^, although this latter may be difficult to distinguish given the similar structures of mitochondrial organelle stains (Figure S1). Naphthalimide-based Probe 1 exhibited a 32-fold fluorescence turn-on upon binding to ribosomal RNA and was reported to have rapid cell uptake with excellent photostability^30^. Indolizine-containing styrene cationic dyes provide possibly the highest signal enhancement (490-fold) among known RNA dyes and enable subcellular visualization through fluorescence lifetime imaging (FLIM)^31^. Finally, PYQU is a multifunctional probe capable of dual-channel detection of RNA and sulfur dioxide (SO_2_) and has been used to visualize endogenous RNA in both cultured cells and zebrafish via deep-red fluorescence although selectivity and light-up data were not reported^32^.

While some of these probes offer improvements in sensitivity and biological applicability, they all rely on non-covalent binding mechanisms of intercalation and electrostatic interaction, which increases off-target background fluorescence for negatively charged and hydrophobic species and environments, and results in poor discrimination between RNA and DNA^33, 34^. Notably, cationic aromatic fluorophores, some with structures closely resembling the reported RNA stains, are also widely used for mitochondrial labeling in cells (Figure S1), illustrating the problem of background and specificity that can result from these low-specificity interactions. Similarly, common commercial RNA stains show little or no selectivity for RNA over DNA. These limitations highlight the need for strategies for labeling that exploit unique physicochemical features of RNA to achieve high selectivity and stable signal generation.

Recent studies have illuminated a key chemical distinction that constitutes a defining feature of RNA as a biopolymer: the presence of the 2′-hydroxyl group (2′-OH) in the ribose sugar. This functional group, absent in DNA, constitutes an inherent chemoselective handle, providing a chemical basis for selective RNA targeting. The 2′-OH group in RNA exhibits a lower pK_a_ (*ca*. 12.5) than reference alcohols (ethanol pKa=15.9) and exhibits unusually high reactivity toward electrophilic agents^35, 36^. Examples of functional electrophiles that react with 2′-OH in high yields include acyl groups^37-39^, sulfonyl groups^40^, and aryl electrophiles^41^. These reactions can proceed efficiently under mild aqueous conditions, showing reactivity that in some cases rivals protein thiol modification in rate and specificity^42, 43^. Covalent reagents developed for 2′-OH modification have been applied to structural probing (*e*.*g*., SHAPE)^44, 45^, labeling^40, 41^, RNA protection and delivery^46, 47^, RNA purification^39, 48^, and interaction profiling^49, 50^, while preserving the underlying native structure of the RNA. Acyl adducts on RNA can be designed to be reversible via chemical triggers or catalysts, releasing unmodified RNA for further analysis^47, 48^. Reagents targeted to 2′-OH can exhibit tunable reactivity, solubility, and cellular compatibility, enabling emerging applications from transcriptome-wide structure mapping to live-cell RNA imaging and RNA-targeted chemical tool development^51^.

To address limitations of existing general RNA-labeling and staining strategies, here we develop a molecular approach that takes advantage of this 2′-OH reactivity (Figure 1b). We describe a previously unexplored class of fluorogenic acylating probes that make use of covalent reactions with RNA. These tunable reagents, termed RiboLight (**RL**) fluorophores, enable sequence-independent labeling of RNA with high selectivity with simultaneous robust signal enhancement (Figure 1c). Notably, **RL** fluorophores are not cationic and thus do not rely on nonspecific electrostatic attractions to interact with their target. Each **RL** fluorophore is designed to emit in a distinct spectral window, enabling the possibility of multicolor imaging and ratiometric analysis. By integrating fluorogenic activation with covalent bond formation in a single step, the **RL** platform overcomes key limitations of non-covalent fluorophores for RNA, such as low specificity, transient binding, modest signal enhancement, and the need for washing steps (Table S1). The probes are compatible with both gel-based and cellular RNA imaging and operate efficiently across diverse RNA sequences and structural contexts. Significantly, the absence of the 2′-OH group in DNA ensures minimal cross-reactivity of **RL** fluorophores, resulting in high levels of RNA:DNA selectivity. Finally, we also find that these fluorophores can be removed, if desired, from their target under mild conditions, restoring unmodified RNA for further analysis. Together, these features establish **RL** fluorophores as useful chemical tools for RNA imaging and analysis *in vitro* and in cells.

## RESULTS AND DISCUSSION

### Design of RNA-selective fluorogenic probes

The **RL** fluorophores were rationally designed to take advantage of two synergistic chemical principles: (1) selective covalent reactivity toward the 2′-OH group of RNA via an acylimidazole functionality and (2) fluorescence enhancement through modulation of twisted intramolecular charge transfer (TICT), achieved by conformational restriction as a result of covalent linkage to the biopolymer (Figure 1b, 1c and 2a). Donor–acceptor (“push–pull”) fluorescent chromophores with rotatable bonds typically undergo non-radiative de-excitation via TICT, resulting in quenched fluorescence in solution^52-54^. Noncovalent association of TICT dyes with biopolymers restricts bond rotations, resulting in increases in fluorescence quantum yields along with bathochromic shifts in absorption and emission^26^. These fluorophores can exhibit high signal-to-noise, potentially enabling wash-free imaging for cellular and gel-based analysis. This molecular strategy has been widely exploited in imaging proteins, DNA and subcellular organelles. However, previous applications to RNA have relied almost invariably on non-covalent interactions involving cationic fluorophores, which suffer from limited selectivity due to nonspecific electrostatic attractions to both RNA and DNA. To overcome this limitation, we designed **RL** fluorophores as noncationic molecular rotors that rely on 2′-OH reactivity to engender RNA selectivity rather than nonspecific cationic attraction. In principle, the electrophilic fluorophores remain non-fluorescent until they undergo rapid reaction with RNA, forming ester links that closely associate the chromophore with the polymer and suppress TICT quenching. This structural constraint induces a marked fluorescence turn-on, while unreacted dye molecules are rapidly hydrolyzed to yield inert, dark byproducts (imidazole and carboxylate), minimizing background and cytotoxicity. Through this combined chemical and photophysical strategy, the **RL** platform enables sequence-independent, covalent, and fluorogenic RNA labeling with minimal cross-reactivity toward DNA.

### RNA-selective reactivity of RL fluorophores

**RL** fluorophores were strategically designed to make dual use of carboxylic acid groups: as the electron acceptor in a push–pull structure, and, in their acylimidazole-activated form, as the electrophilic moiety for RNA-specific covalent labeling (Figure S2, see Supporting Information for synthetic details and characterization). We synthesized carboxylic acid precursors of four **RL** fluorophores (**CA420, CA450, CA480** and **CA560**, numerically designated by their absorption maxima) in one to three steps from commercial reagents (Figure S3-S10). These precursors were then activated via 1,1′-carbonyldiimidazole (CDI)-mediated coupling in anhydrous dimethyl sulfoxide (DMSO), directly generating 1 M stock solutions of the reactive dyes with one equivalent of imidazole. The reactions proceeded in high yields without the need for further purification (Figures S11–S22).

To evaluate RNA reactivity, the **RL** fluorophores (50 mM) were incubated with a short single-stranded RNA (ssRNA) oligonucleotide in nuclease-free water containing 5% DMSO at room temperature. Covalent adduct formation was analyzed by MALDI-TOF mass spectroscopy following purification (Figures 2b–g and Figures S23–S30). All four dyes showed robust reactivity within 1 h, forming one or more adducts per RNA strand. Although most applications would utilize low-occupancy labeling (see below), we confirmed that extended incubation yielded higher adduct numbers. **RL420** and **RL450** displayed the highest RNA reactivity, forming 5–6 adducts per strand with conversion rates approaching 90%. **RL480** showed intermediate reactivity, and the bulkier **RL560** exhibited lower conversion (*ca*. 15%) with a maximum of three adducts.

**Figure 2.**
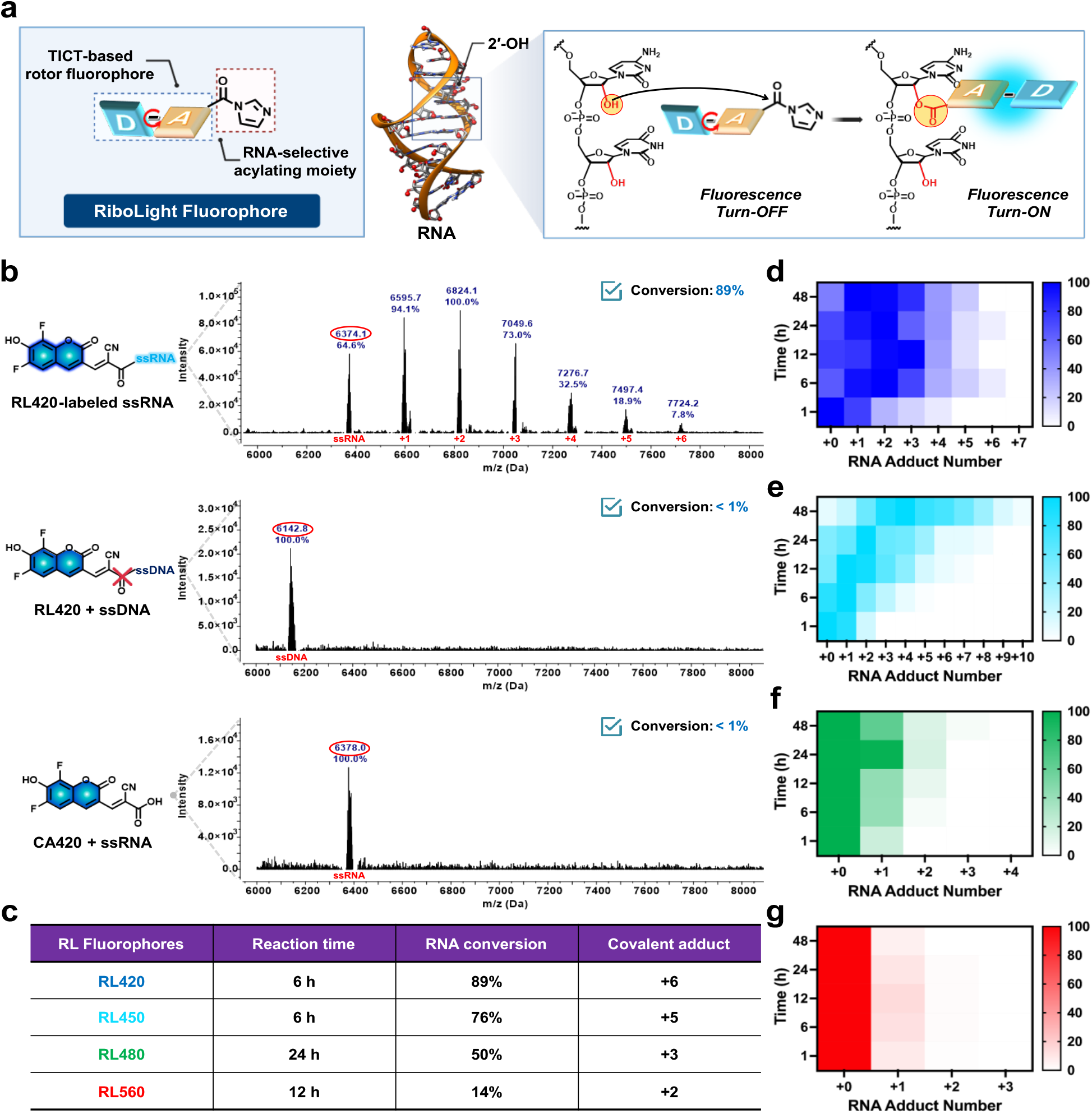
Covalent labeling of RNA via 2′-OH-selective acylation with RL fluorophores. (a) RL fluorophores were designed with a twisted intramolecular charge transfer (TICT)-based rotor fluorophore conjugated to an electrophilic acylimidazole, enabling covalent reaction with the RNA 2′-OH, a functional group absent in DNA. This nucleophile-guided acylation provides selective RNA labeling and induces fluorescence turn-on. (b) MALDI-TOF mass spectra of acylation reactions confirms multiple covalent adducts (+1 to +6) on ssRNA following treatment with RL420. No modification was observed for ssDNA under identical conditions, confirming RNA selectivity. The control compound CA420, which lacks an acylating moiety, showed no covalent modification of RNA, indicating that labeling requires specific acylation chemistry. (c) Summary of RL fluorophore acylating performance, including reaction times, overall conversion efficiencies and the number of covalent adducts formed, as determined by MALDI-TOF analysis. Conversion rate is defined as the proportion of RNA modified at one or more 2′-OH positions. (d–g) Heat maps showing the distribution of covalent adduct numbers for each RL fluorophore on ssRNA, as determined by MALDI–TOF.

To confirm RNA specificity and identify the reactive site, we performed parallel reactions with a DNA oligonucleotide of identical sequence. None of the dyes showed detectable reactivity with DNA, indicating that the 2′-OH group, which is present only in RNA, is essential for covalent bond formation. These results rule out significant contributions from reactions with nucleobase exocyclic amines or 5′ or 3′ terminal hydroxyls. Control experiments using the unactivated carboxylic acid precursors failed to produce any RNA adducts (Fig. 2b and Figures S23, S25, S27 and S29). Taken together, these data demonstrate that **RL** fluorophores undergo rapid, chemoselective 2′-OH acylation to form ester-linked adducts on RNA, enabling RNA-specific labeling with minimal cross-reactivity toward DNA. The marked selectivity over DNA provides a molecular basis for an RNA-specific fluorescence response.

**RL**-labeled RNAs can remain stable for extended periods. Although labeled RNA is sufficiently stable for most short-term applications such as gel or cellular imaging, follow-on use in certain molecular or cellular assays may require reversibility of the covalent dye adducts. Acyl adducts on RNA can impede reverse transcription, and when located within open reading frames (ORFs) of messenger RNA (mRNA), can interfere with ribosomal translation^55^. To address this issue, we first examined the stability of **RL** fluorophore modification by incubating **RL450**-labeled ssRNA in water at room temperature for up to 7 d with

MALDI-TOF analysis at defined timepoints (Figure S31). The adducts remained largely intact throughout, with only minor hydrolysis observed, indicating sufficient stability under aqueous conditions for most standard applications. We then asked whether the unmodified RNA could be recovered for further analytical and biological processes. Recent studies have shown that nucleophiles such as N,N-dimethylaminopyridine (DMAP) can de-acylate groups from RNA^47^. **RL450**-labeled ssRNA was co-incubated with 100 mM DMAP at room temperature, and time-dependent loss of adducts was monitored by MALDI-TOF mass spectroscopy (Figure S32). Nearly complete removal of the fluorophore was evident by 48 h, accompanied by restoration of the original unmodified RNA. These results demonstrate that **RL** fluorophores can be stably retained on RNA when desired, yet can be effectively removed under mild nucleophilic conditions, offering both durable labeling and reversibility for downstream applications.

### Photophysical properties and responses of RL fluorophores

Having established efficient and selective covalent labeling of RNA, we next evaluated the photophysical properties of the **RL** fluorophores both in free form and following reaction with RNA. Absorption and emission spectra were acquired for the unmodified carboxylic acid precursors, the activated acylimidazoles, and the covalently RNA-labeled forms (Figures 3, Table 1, Figures S33 and S34). Each form exhibited distinct absorption maxima, reflecting changes in electronic structure upon CDI activation and subsequent RNA conjugation (Figure S33).

**Table 1.** Photophysical properties of RL fluorophores.

| Fluorophore | Solvent<sup>a</sup> | $\lambda_{\text{Abs}}^b$ (nm) | $\lambda_{\text{Em}}^c$ (nm) | Stokes shift (nm) | $\Phi_{\text{Fl}}^{d,e}$ (%) | Fold-increase<sup>f</sup> |
| --- | --- | --- | --- | --- | --- | --- |
| RL420 | PBS | 441 | 450 | 9.0 | 1.1 | 81 |
|  | ssDNA | 403 | 444 | 41 | 4.3 | 150 |
|  | ssRNA | 395 | 452 | 57 | 30 | - |
| RL450 | PBS | 459 | 513 | 54 | 0.03 | 390 |
|  | ssDNA | 446 | 483 | 37 | 0.15 | 970 |
|  | ssRNA | 447 | 497 | 50 | 2.6 | - |
| RL480 | PBS | 483 | 513 | 30 | 0.03 | 320 |
|  | ssDNA | 481 | 508 | 27 | 0.18 | 220 |
|  | ssRNA | 479 | 514 | 35 | 8.1 | - |
| RL560 | PBS | 579 | 647 | 68 | 0.28 | 24 |
|  | ssDNA | 574 | 629 | 55 | 0.80 | 71 |
|  | ssRNA | 556 | 639 | 83 | 16 | - |

<sup>a</sup>PBS: phosphate buffer saline (1X, pH = 7.4). ssDNA and ssRNA were dissolved in PBS (1X, pH = 7.4). Measurements for ssRNA and ssDNA were performed using 20-nt oligomers with identical sequences. <sup>b</sup>Absorbance maximum wavelength <sup>c</sup>Fluorescence maximum wavelength. <sup>d</sup>Determined vs. fluorescein in 0.1 N NaOH for **RL420**, **RL450**, and **RL480**. <sup>e</sup>Determined vs. Cy5 in DMSO for **RL560**. <sup>f</sup>Fold-increase values were calculated by comparing the maximum fluorescence emission intensities of ssRNA-reacted samples upon excitation at the same wavelength.

**Figure 3.**
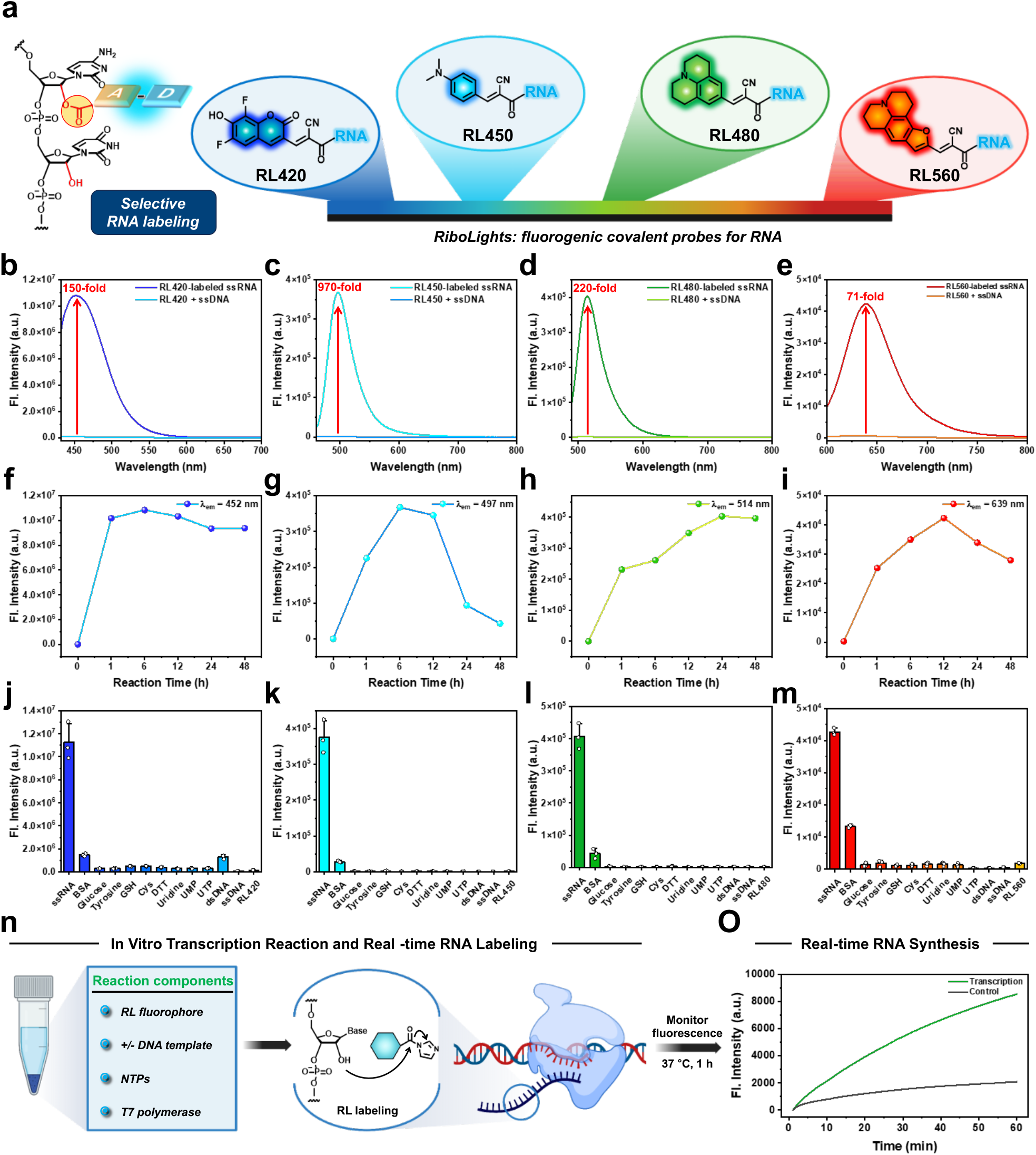
Multicolor fluorogenic RNA labeling via 2′-OH-selective acylation with RL fluorophores. (a) Schematic illustration of the **RL** platform comprising four wavelength-tuned fluorophores (**RL420, RL450, RL480** and **RL560**) that selectively react with the 2′-OH of RNA, enabling covalent fluorescence turn-on labeling. (b–e) Fluorescence emission spectra of **RL420** (b), **RL450** (c), **RL480** (d) and **RL560** (e) measured separately in the presence of ssRNA or ssDNA of identical sequence. (f–i) Time-dependent fluorescence enhancement of **RL420** (f), **RL450** (g), **RL480** (h) and **RL560** (i) following covalent acylation with ssRNA over 0–48 h, indicating fluorophore-specific kinetics and fluorescence activation profiles. (j–m) Fluorescence responses of **RL420** (j), **RL450** (k), **RL480** (l) and **RL560** (m) toward potential interferents, confirming selective response to RNA. Bar graphs represent mean fluorescence intensities ± *s*.*d*. from triplicate experiments. (n) Schematic illustration of real-time detection of post-transcriptional RNA synthesis using T7 transcription in the presence of **RL480**. (o) Real-time fluorescence monitoring of *in vitro* transcribed RNA using **RL480**, showing active transcription-dependent signal enhancement.

Notably, fluorescence emission spectra revealed robust turn-on enhancements upon reaction with RNA (Figures 3b–e). In contrast, measurements in the presence of ssDNA of identical sequence displayed minimal fluorescence, underscoring the high selectivity of 2′-OH covalent labeling. The magnitude of fluorescence enhancement varied from 71-fold (**RL560**) to 970-fold (**RL450**). To our knowledge, this level of selectivity in fluorescence enhancement for RNA over DNA is unprecedented, far exceeding levels reported for commercial RNA dyes or previously published dyes, both in RNA selectivity and in degree of fluorescence light-up.

Profiling the time-dependence of labeling confirmed progressive increases in emission as labeling proceeded (Figures 3f–i). Interestingly, **RL450** reached peak fluorescence intensities at 6 h, followed by moderate declines thereafter. MALDI-TOF analysis verified that the decrease was not due to loss of labeled adducts, but rather is likely attributable to dye-dye self-quenching effects^56^ arising from high-density labeling of up to seven 2′-OH groups out of twenty in the test RNA (Figure S26).

Examination of the emission spectra of the four **RL** fluorophores reveals well-separated emission spectra after RNA conjugation, enabling the potential for multicolor imaging or multiplexed detection (Figures 3b–e). Excitation wavelengths for each fluorophore fall within standard filter sets: **RL420** can be excited in the coumarin 343 range, **RL450** overlaps with cyan fluorescent protein (CFP)-type channels, **RL480** is compatible with fluorescein (FITC/Alexa Fluor 488) channels and **RL560** aligns with Cy3/TAMRA excitation profiles. While their brightness does not match that of those highly optimized commercial fluorophores, owing to the TICT-based design and strong dipolar structures, the **RL** fluorophores on RNA still showed moderate to good quantum yields, ranging from 2% to 30% (Table 1). Stokes shifts of the four dyes with RNA are 35-83 nm, larger than those of many common dye classes such as cyanine and BODIPY chromophores, thus providing a wide spectral window for varied excitation sources and filter sets.

To investigate the mechanism underlying fluorogenic activation, we measured the emission spectra of unreacted **RL** fluorophores in solvents of increasing viscosity (Figure S35). For **RL450, RL480**, and **RL560**, increasing viscosity led to bathochromic shifts and significant fluorescence enhancement, consistent with restricted bond rotations suppressing TICT de-excitation^57^. This supports the hypothesis that covalent conjugation to RNA restricts intramolecular rotation and activates fluorescence. One exception to this last phenomenon was **RL420**, which exhibited a redshift in emission without intensity enhancement, suggesting that its fluorescence is more strongly influenced by reduced polarity rather than by changes in viscosity^58, 59^.

The above experiments provided documentation of the large selectivity of the **RL** fluorophores in providing signals for RNA over DNA. However, in some applications such as cellular imaging, biological nucleophiles are abundant and constitute potential sources of background signals if the nucleophiles can react with the acylimidazole electrophile in **RL** fluorophores. To address this issue, we assessed their fluorescence responses in the presence of potentially competing nucleophiles to evaluate chemical selectivity. Each **RL** fluorophore was incubated with a range of analytes, including ssRNA, ssDNA, dsDNA, bovine serum albumin (BSA), and small-molecule nucleophiles such as tyrosine, cysteine, glucose, and uridine. All four dyes demonstrated strong fluorescence activation with RNA, minimal response with DNA, and negligible signal with small molecules (Figures 3j–m). BSA yielded modest but significant signals, likely arising from reaction with amino acid residues accompanied with partial restriction of chromophore rotations. The low background seen with small molecules, despite their highly nucleophilic groups, is presumably due not to absence of reaction with **RL** fluorophores, but rather to the inability of small adducts to sufficiently constrain bond rotations and suppress TICT-associated quenching. Taken together, these findings highlight robustness, selectivity, and fluorogenic behavior of **RL** fluorophores as covalent RNA labeling agents.

### Real-time reporting on RNA synthesis

We further explored the utility of **RL** fluorophores for real-time monitoring of RNA synthesis by performing a standard *in vitro* transcription assay supplemented with **RL480**. The reaction included T7 RNA polymerase, all four NTPs, and a linearized plasmid DNA template. A no-template control was included to compare fluorescence intensities between mixtures with active RNA synthesis those without (Figure 3n). Fluorescence was continuously measured using a real-time PCR instrument. Over the course of the reaction, we observed a time-dependent increase in fluorescence signal which correlates with the progressive accumulation of RNA transcripts (Figure 3o). These results demonstrate that **RL480** can report on newly synthesized RNA as it is produced, and the fluorescence enhancement enables realtime monitoring of transcriptional activity. Notably, the data suggest little off-target activity of **RL480** with the RNA polymerase, as the enzyme continued to function. Although labeling is covalent, as described above the adducts can be removed upon treatment with DMAP. In principle, just as RNA synthesis can be monitored in real-time, the removal of dye adducts might also be tracked by the diminishment of fluorescence.

### Application in PAGE gel imaging

To evaluate the compatibility of **RL** fluorophores with polyacrylamide gel electrophoresis (PAGE) and gel imaging, we conducted experiments using both a short ssRNA and an RNA ladder containing transcripts of varying lengths. Oligonucleotides were labeled by incubating with **RL** probes for 12 to 24 h, followed by purification via ethanol precipitation. The labeled samples were resolved by PAGE and imaged for fluorescence. Clear fluorescent bands were observed for RNA oligonucleotides, but not for DNA oligonucleotides, confirming the RNA selectivity of **RL** fluorophores (Figures 4a–b). The results were in marked contrast with SYBR Gold, which labeled both RNA and DNA nearly equally. **RL450** and **RL480** also successfully labeled the RNA ladder, demonstrating that RNA strands as long as 1000-nt can be visualized (Figure 4c). Because a 24 h incubation may be impractical in certain experiments, we also tested whether a 1 h incubation under the same conditions would yield comparable results. We found that labeling for as little as 1 h produced similar results, demonstrating that this approach can easily be integrated into experimental workflows (Figure S36). We also note that covalent labeling can modestly alter RNA mobility on the gel, particularly for short oligonucleotides (Figure 4a). This effect can be mitigated by using lower-percentage acrylamide gels and shortening electrophoresis time to preserve resolution.

**Figure 4.**
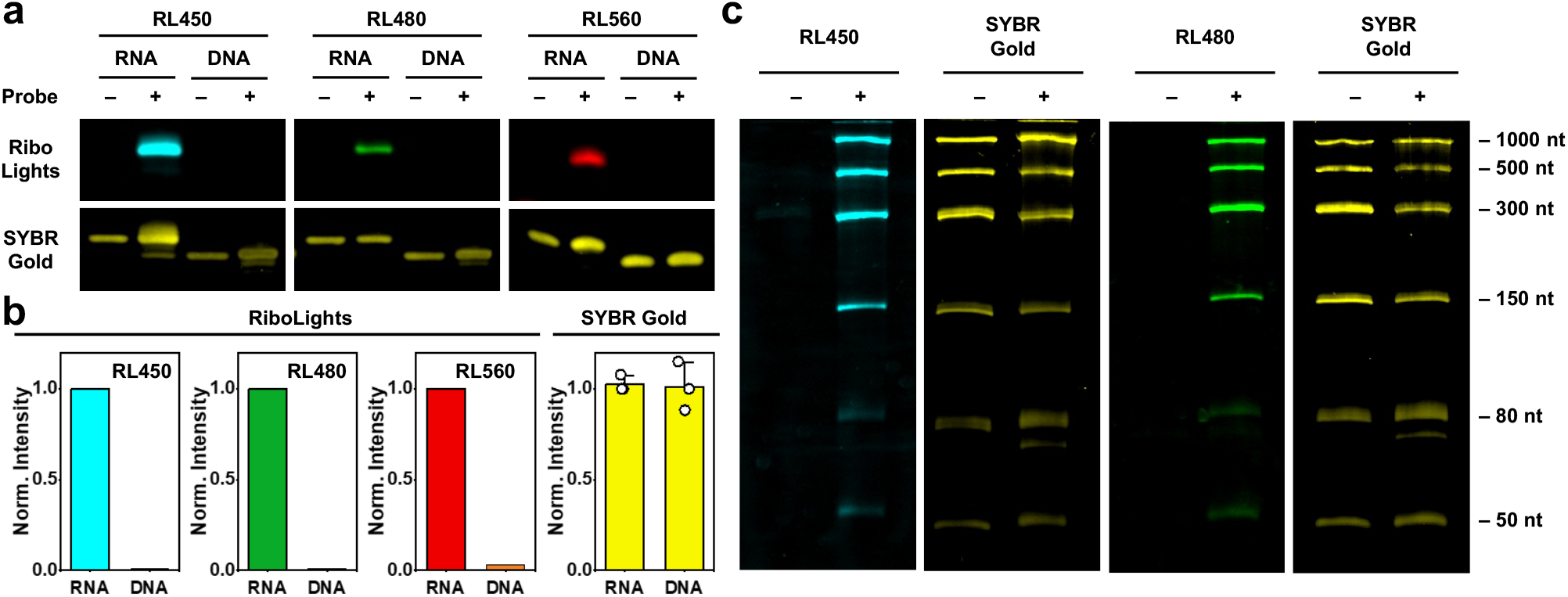
RNA-selective gel imaging using **RL** fluorophores. (a) PAGE analysis of ssRNA and ssDNA oligonucleotides (18-nt) labeled with **RL450, RL480** or **RL560** and visualized by in-gel fluorescence. “+” denotes sample incubated with **RL** fluorophores and “-” denotes untreated controls. Strong fluorescence bands were observed for RNA but not for DNA, confirming RNA selectivity. In contrast, SYBR Gold stained both RNA and DNA nearly equally. (b) Quantification of fluorescence intensity from the gels in a, showing high RNA-to-DNA selectivity for **RL** fluorophores. Bar graph of SYBR Gold represents mean fluorescence intensities ± *s*.*d*. from triplicate experiments. (c) Labeling of RNA ladders with **RL450** (12 h) or **RL480** (24 h) enabled visualization of RNA strands up to 1000-nt. SYBR Gold staining of the same gels is shown for comparison.

### RL fluorophores in cellular imaging

Encouraged by the performance of the probes in solution and in gel imaging, we next explored their utility in cellular imaging. To assess this, HeLa cells were incubated with **RL** fluorophores and visualized by confocal fluorescence microscopy. The julolidine-based green and red fluorophores **RL480** and **RL560** exhibited ready cell permeability and robust labeling in live cells (Figure 5a). Cells were well stained with 20 μM dye for 30 min. No washes were required for imaging, reflecting the high fluorescence enhancement upon covalent labeling, as well as the lack of fluorescence both in unreacted dye and the carboxylate byproducts of dye hydrolysis. In contrast, the blue and cyan dyes **RL420** and **RL450** showed limited permeability and required cell fixation and permeabilization prior to labeling for effective staining. Strong fluorescence signals were observed from the nucleolus, a compartment enriched in ribosomal RNA^30, 31^, indicating that the dyes retain their RNA selectivity and reactivity in the cellular environment (Figure 5a). In contrast, cells treated with the non-reactive carboxylic acid precursor **CA480** showed no detectable signals, confirming that the observed light-up fluorescence arises from covalent labeling of RNA (Figure 5b). To validate this, DNA and RNA extracted from **RL480**-labeled cells were analyzed by dot blot, revealing fluorescence from RNA exclusively (Figure S37). Notably, SYTO RNASelect failed to show comparable selectivity under similar conditions (Figure S38). Additional fluorescence was detected in the cytoplasm, with signal distributions similar to those reported for other RNA-selective dyes (Figure S39a)^60, 61^. Co-staining with MitoTracker and LysoTracker showed only moderate overlap, presumably due to the lack of cationic structure in **RL** probes. (Figures S39b–c).

**Figure 5.**
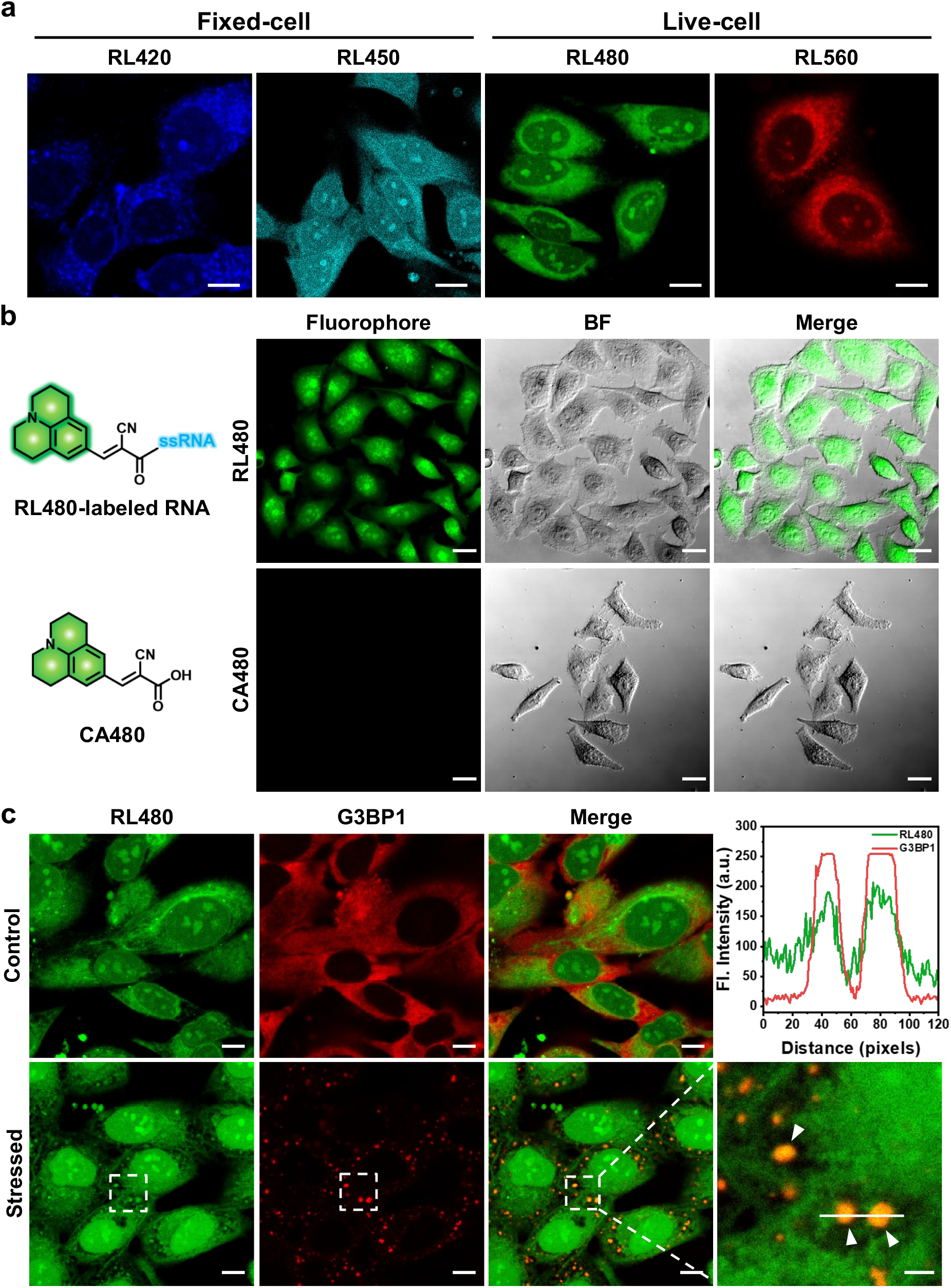
Cellular imaging of RNA using RL fluorophores. (a) Confocal fluorescence microscopy of HeLa cells labeled with **RL** fluorophores. Live-cell imaging with the green-emissive **RL480** and red-emissive **RL560** (20 μM, 30 min, no wash) showed strong intracellular fluorescence, whereas blue-emissive **RL420** and cyan-emissive **RL450** required fixation and permeabilization for effective labeling. Nucleolar signal enrichment was observed, consistent with RNA selectivity. (b) Live HeLa cells treated with **RL480** or the non-reactive carboxylic acid precursor **CA480**. The precursor lacking the acylating moiety showed no detectable fluorescence, confirming that the observed signal originates from covalent RNA labeling. (c) Co-localization analysis of **RL480** with stress granule marker G3BP1 in sodium arsenite–treated HeLa cells. Merged images and intensity profiles confirm **RL480** labeling of RNA within stress granules. Scale bars: 10 μm (a, c), 20 μm (b) and 5 μm (zoom in c).

To assess whether **RL** fluorophores can label dynamic RNA-rich structures, we induced stress granule formation in HeLa cells by treating them with sodium arsenite, followed by immunostaining with anti-G3BP1, a well-established stress granule marker. Our results show that **RL480** fluorescence colocalizes with G3BP1 signals, indicative of successful labeling of RNA within these compartments (Figure 5c). Finally, we find that **RL** fluorophores display no significant cytotoxicity at labeling concentrations (Figure S40). This low toxicity may be attributed to the inherently short lifetime of the acylimidazole warhead (t_1/2_ for hydrolysis of acylimidazoles can be as short as 30 min in aqueous solution)^62^, limiting prolonged or excessive intracellular labeling even at high concentrations.

## CONCLUSIONS

Our experiments establish a previously unexplored strategy for general RNA labeling, yielding the first ribopolymer-specific dyes. The design relies on covalent bond formation at 2′-OH to engender selectivity for ribonucleic acids, simultaneously coupled with robust increases in fluorescence emission due to restriction of bond rotations. Prior RNA dyes have relied on relatively nonspecific noncovalent association of cationic aromatic dyes with the polyanionic polymer; the current strategy results in far higher selectivity for RNA over DNA by taking advantage of the elevated reactivity of the 2′-OH group in RNA to acylating reagents in water. We find that the covalent RL dyes yield greater fluorescence enhancements with RNA – as large as 970-fold - than do prior RNA dyes; while some of those previous dyes also make use of TICT bond rotational restriction, we speculate that the uniformly closer association of the bonded dyes results in greater restriction of bond rotations, as all **RL** fluorophores are tightly constrained at a single bond length from the bulky biopolymer. Importantly, the **RL** probes do not respond to common strongly nucleophilic small interferents found in biological systems, such as the thiols of cysteine or glutathione, presumably because rotational bond restriction is inefficient with a non-bulky nucleophile. Our results demonstrate that **RL** fluorophores are broadly applicable across multiple RNA labeling contexts, including gel-based RNA detection, monitoring RNA synthesis in real-time, and wash-free cellular imaging. A notable finding is the low cytotoxicity of the **RL** platform, even at concentrations far exceeding typical working ranges and over extended incubation times. This was initially unexpected, given the possibility of covalent reaction of acylimidazole electrophiles with essential biological components. We attribute this low toxicity in part to the short half-life for hydrolysis of the acylimidazole warheads, giving the dyes limited time to react nonspecifically with off-targets. In contrast, the high reactivity of activated **RL** fluorophores with the targeted RNA allows rapid labeling at relatively low concentrations. The anionic carboxylate byproducts are permanently dark and exhibit apparently low toxicity.

Some limitations of **RL** fluorophores are worth noting and can be addressed with additional work. As noted above, covalent modification of RNA may interfere with downstream enzymatic processes such as reverse transcription and can slightly alter the gel mobility of short RNAs. Most imaging of RNA is performed without the need for further manipulation, but if such post-labeling analysis is desired, we find that it can be mitigated by leveraging the dyes’ reversibility in the presence of DMAP, allowing recovery of fully unmodified RNA. A second limitation of the current set of **RL** fluorophores is that, while **RL480** and **RL560** are readily cell permeable, the blue- and cyan-emitting dyes **RL420** and **RL450** showed limited cell permeability, restricting their use to fixed-cell applications. These challenges motivate structural optimization efforts for future variants of **RL** fluorophores. Given that there are many possible TICT chromophores based on the current generic design, this leaves ample room for further testing and improvement.

Future work will focus on expanding the spectral diversity of **RL** fluorophores, as well as on applications. Testing and development of a broader range of emission palettes seems possible, simply by converting carboxylic acid acceptor groups in push-pull chromophores to acylimidazole groups. Of special interest would be **RL** fluorophores emitting in deep-red and near-infrared wavelengths for tissue imaging applications. Also possible is the use of multiplexing in labeling, such as in pulse-chase experiments, which is potentially feasible with covalent dyes in contrast to noncovalent dyes that can rapidly exchange between RNAs.

## ASSOCIATED CONTENT

The Supporting Information is available free of charge at http://pubs.acs.org./doi/.

## AUTHOR INFORMATION

### Authors

Jinwoo Shin - Department of Chemistry, Stanford Cancer Institute, Stanford University, Stanford, CA, 94305, USA.

Moon Jung Kim - Department of Chemistry, Stanford Cancer Institute, Stanford University, Stanford, CA, 94305, USA.

### Author Contributions

J.S. synthesized and characterized the fluorophores. J.S. and M.J.K. conducted the gel and biological assays. J.S. and E.T.K. conceived and designed the project. J.S., M.J.K., and E.T.K. wrote the manuscript. E.T.K. provided project supervision.

^†^These authors contributed equally.

### Notes

The authors declare no competing financial interests.

## ACKNOWLEDGMENT

This work was supported by the U.S. National Institutes of Health (GM145357). Additional support for J.S. was provided by the Basic Science Research Program of the National Research Foundation of Korea (NRF) funded by the Ministry of Education (RS-2024-00412225).

## Table of Contents (TOC)

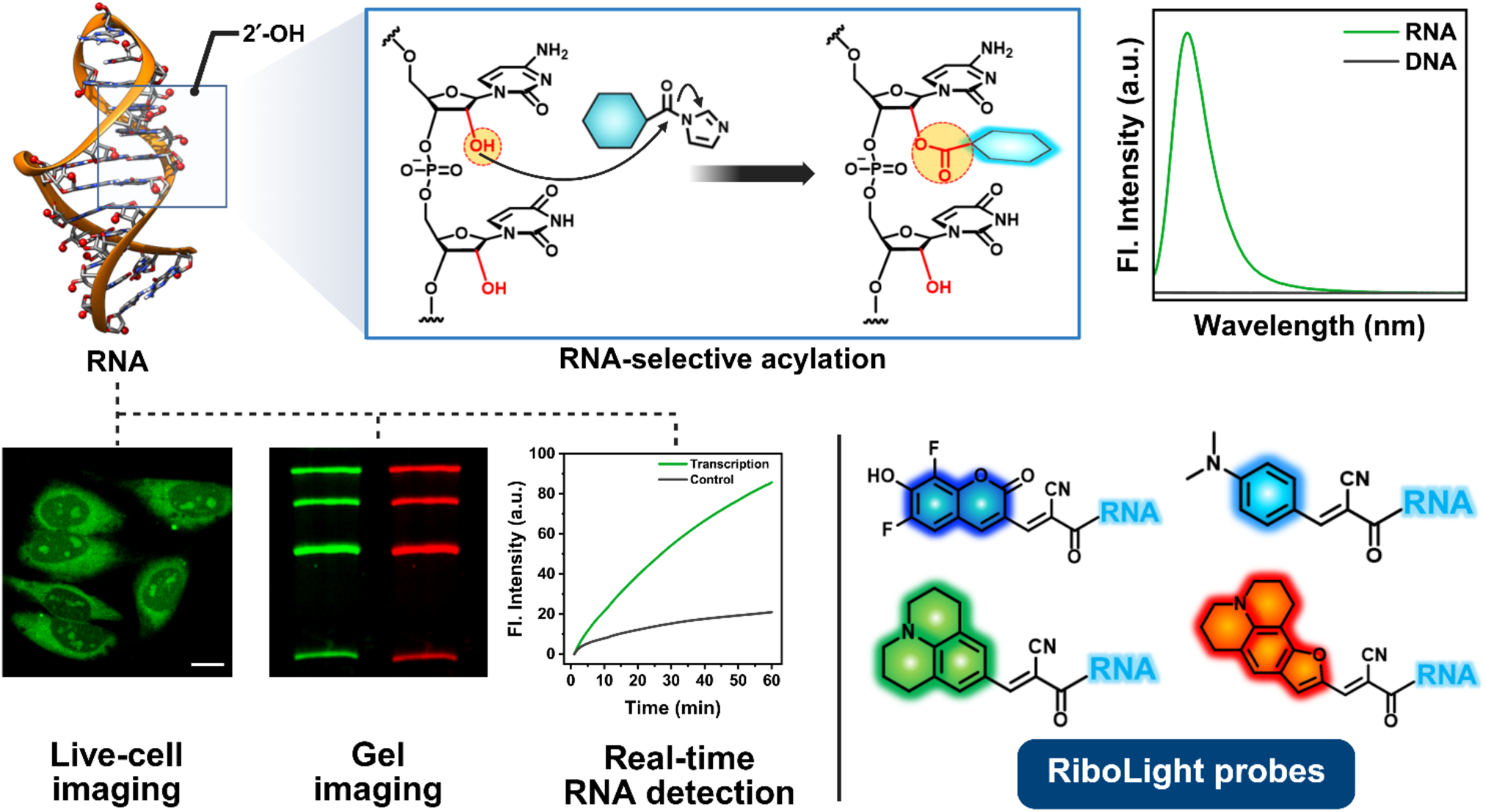

