## Supporting Information for "Fluorogenic Covalent Probes for RNA"

INSTRUMENTATION

All Nuclear magnetic resonance (NMR) spectra were recorded on a Bruker 400 MHz NMR instrument in Stanford University Department of Chemistry NMR facility. The spectra were analyzed using MestReNova (MNova, Mestrelab Research) software. All electrospray ionization mass spectrometry (ESI-MS) spectra were recorded at the Stanford University Mass Spectrometry Facility. The instrument used a Waters 2795 high-performance liquid chromatography (HPLC) system with dual wavelength ultraviolet (UV) detector, and ZQ single quadrupole mass spectrometry with electrospray ionization (ESI) source. All matrix-assisted laser desorption/ionization time-of-flight (MALDI–TOF) mass spectrometry spectra were recorded at the Stanford University Mass Spectrometry facility,. The spectral data was then recorded using Flex Control software (Bruker) and analyzed using MNova software. UV–visible (UV/Vis) spectra were recorded on a NanoDrop One Spectrophotometer (Thermo Scientific Invitrogen) and fluorescence emission spectra were recorded on a Fluorolog 3-221 spectrofluorometer (Horiba Jobin Yvon) with an external temperature controller. Luminescence readings were obtained on a Fluoroskan Ascent FL Microplate Reader (Thermo Fisher Scientific). Cellular images were recorded on an inverted Zeiss LSM 780 multiphoton laser scanning confocal microscope and Leica SP8 X White Light Laser Confocal Microscope in Cell Science Imaging Facility (CSIF) at Stanford. Fluorescent gel images were captured using the iBright FL1500 Imaging System (Thermo Fisher Scientific) and analyzed and quantified using ImageJ. Real-time detection of RNA synthesis from in vitro transcription reactions was performed using the Applied Biosystems StepOnePlus Real-Time PCR System. Molecular graphics and structural illustrations of single-stranded RNA (ssRNA) and double-stranded DNA (dsDNA) were prepared using UCSF Chimera (version 1.13.1). The ssRNA model was generated by manual modification of the 6XH2 PDB, while the dsDNA model was based on the 4YNQ PDB structure. Graphical abstract and figures were created in BioRender. Shin, J. (2026) https://BioRender.com/ihyeg9t.

METHODS

**Materials**

All chemical reagents and solvents used for synthesis were purchased from commercial suppliers (Merck, TCI and Santa Cruz) and used without further purification. Custom oligonucleotides were obtained from Integrated DNA Technologies (IDT) with high-performance liquid chromatography (HPLC) purification unless otherwise specified (see Supplementary Table 2 for sequences). All oligonucleotide cleanup reactions were performed using RNA Clean & Concentrator-5 Kit (Zymo Research Corporation). *In vitro* transcription was performed using HiScribe® T7 High Yield RNA Synthesis Kit (New England Biolabs). All polyacrylamide gels were prepared using the SequaGel® UreaGel System from National Diagnostics. HeLa human cervical cancer cell line (ATCC, CRM-CCL-2) was cultured in Dulbecco’s modified Eagle medium (DMEM) supplemented with 10% fetal bovine serum (FBS) at 37 °C in a humidified incubator containing 5% CO_2_. LDH-Glo Cytotoxicity Assay kit for cell viability and cytotoxicity measurements was purchased from Promega. Primary antibody, anti-G3BP1 (#61559) was purchased from Cell Signaling Technology and Goat anti-Rabbit IgG-Alexa Fluor™ 647 was purchased from Thermo Fisher Scientific.

**2′-OH Acylation of oligonucleotides**

All acylation reactions were performed in sterile 200 µL PCR tubes. To each tube, 9.5 µL of oligonucleotide stock solution (52.63 µM) was added and pre-cooled on ice for 30 min. Freshly prepared **RL fluorophores** (1 M in DMSO) were added (0.5 μL) to achieve a final concentration of 50 mM. The reaction mixtures were incubated at 37 °C for the desired reaction time. After incubation, samples were purified by either ethanol precipitation or spin column purification. The resulting pellet was either stored at -80 °C or redissolved in nuclease-free water or buffer for use in further experiments. Acylation efficiency was assessed by MALDI–TOF mass spectrometry and visualized by PAGE. Reaction conversion was defined as the percentage of oligonucleotides modified at one or more 2′-OH positions. Data were analysed using MestReNova (MNova, Mestrelab Research) software.

**Mass spectrometric analysis of RNA reactions**

Mass spectra of reacted RNAs were obtained with a Bruker Daltonik Microflex MALDI–TOF mass spectrometry equipped with an N_2_ laser. All spectra were recorded in linear negative mode and samples were plated on an MSP Anchorchip 96 target plate. 0.3 M trihydroxyacetophenone (THAP) in ethanol (matrix) and 0.1 M aqueous ammonium citrate (co-matrix) were mixed in a 2:1 ratio by volume to be used as a matrix mix. This mixture was always freshly prepared before analysis. After 2′-OH acylation of oligonucleotides, the resulting pellets were redissolved in 10 µL nuclease-free water. 1 μL of this solution was transferred to the target plate and dried under an Ar stream. 1 μL of the matrix mixture was then added directly on top of the dried sample and completely dried under Ar stream.

**Purification of oligonucleotides using ethanol precipitation and Spin column purification**

For ethanol precipitation, 90 μL of precipitation solution (composed of 10 μL of 1 M MgCl₂, 10 μL of 2 mg/mL glycogen, and 980 μL of nuclease-free water) was added to each reaction mixture in a 1.5 mL sterile microcentrifuge tube. Ice-cold absolute ethanol (3 volumes) was added, followed by vortexing for 30 s. Samples were stored at -80 °C overnight, followed by centrifugation at 14,800 rpm for 30 min at 4 °C. The supernatant was discarded, and the resulting pellet was washed with 70% ethanol. After air drying for 15 min, the pellet was either stored at -80 °C or redissolved in nuclease-free water or buffer for use in further experiments. For spin column purification, the Zymo RNA Clean & *Concentrator*-5 Kit was used according to the manufacturer’s instructions. Briefly, the reaction was diluted with 20 μL of RNA Binding Buffer and 30 μL of 96% ethanol. The mixture was transferred to a Zymo-Spin IC Column and centrifuged. The column was then washed with 400 μL of RNA Prep Buffer, followed by two washes with 700 μL and 400 μL of RNA Wash Buffer. The labeled RNA was eluted in 10 μL of nuclease-free water for use in subsequent experiments.

**Photophysical measurements**

For UV/Vis absorption measurements, reaction pellets were dissolved in 10 μL of nuclease-free water, and spectra were recorded using a NanoDrop One spectrophotometer (Thermo Scientific Invitrogen) with 1 μL of sample. Baseline correction was applied at 800 nm. Fluorescence emission spectra were acquired using a Fluorolog 3-221 spectrofluorometer (Horiba Jobin Yvon) with excitation wavelengths optimized for each **RL fluorophore**. For each measurement, 5 μL of sample was mixed with 495 μL of PBS in a narrow quartz cuvette. Fluorescence enhancement was quantified as the fold increase in emission intensity of RNA-labeled samples relative to unlabeled controls at their respective emission maxima.

**Fluorescence response assay with potential analytes**

To compare fluorescence responses with potential analytes, **RL fluorophores** were prepared as 100× stock solutions in DMSO. The stock concentration of each fluorophore was adjusted such that its absorbance matched the absorption intensity observed in modified RNA samples at the corresponding excitation wavelength, as determined by UV/Vis spectroscopy. For each assay, 5 μL of **RL fluorophore** stock solution analyte (100× stock in DMSO) and 5 μL of potential analyte (100× stock in nuclease-free water) were combined with 490 μL of PBS in a narrow quartz cuvette. Fluorescence spectra were recorded immediately using a Fluorolog 3-221 spectrofluorometer (Horiba Jobin Yvon) under the same excitation and emission settings as those used for RNA-labeled samples.

**Quantum yield determination**

To minimize the inner filter effect, fluorescence measurements were conducted under conditions in which the absorbance remained below 0.1 at all wavelengths equal to or longer than the excitation wavelength. Quantum yields (Ф_Fl_) of **RL 420**, **RL450**, and **RL480** were determined relative to fluorescein in 0.1 N NaOH (Ф_Fl_ = 0.92), while **RL560** was referenced to Cy5 in DMSO (Ф_Fl_ = 0.33).

**Solvent-dependent fluorescence spectroscopy with increasing viscosity**

Emission spectra of **RL fluorophores** (10 μM) were recorded at 37 °C in various solvents, including methanol (CH₃OH), ethylene glycol, a 1:1 (v/v) mixture of ethylene glycol and glycerol, and glycerol, using a Fluorolog 3-221 spectrofluorometer (Horiba Jobin Yvon). Stock solutions were prepared in DMSO, and all final solutions contained 1% (v/v) DMSO.

**Stability assay of RL-labeled ssRNA in nuclease-free water**

To assess the stability of RNA labeling, 10 μL reaction mixtures containing 18-nt ssRNA and **RL450** were prepared in sterile 200 μL PCR tubes following the 2′-OH acylation procedure. The reaction was incubated at 37 °C for 1 h to allow covalent labeling. **RL**-labeled RNA was then purified by ethanol precipitation and resuspended in 10 μL of nuclease-free water. Sample was stored at room temperature without light protection. At specific time points (starting, 1 d and 7 d), 1 μL aliquots was taken and analyzed directly by MALDI–TOF mass spectrometry as described above. The number of covalent adducts was determined from the observed mass shifts, and the relative abundance of unmodified and modified RNA species was calculated from peak intensities using MNova software.

**Reversibility of RL-labeled RNA via DMAP treatment**

To evaluate the reversibility of RNA labeling, 10 μL reaction mixtures were prepared in sterile 200 μL PCR tubes by combining the following components: 9.0 μL of ssRNA stock solution (50 μM, final 45 μM), 0.5 μL of **RL** **fluorophore** stock (1 M in DMSO, final 50 mM, 5% DMSO), and 0.5 μL of DMAP stock (2 M in nuclease-free water, final 100 mM). For the control group without DMAP, 0.5 μL of nuclease-free water was added in place of DMAP. Reactions were incubated at room temperature for up to 48 h. At specified time points (1, 6, 12, 24, and 48 h), 1 μL of each reaction mixture was directly analyzed by MALDI–TOF mass spectrometry as described above. The degree of deacylation was assessed by monitoring the disappearance of acylated RNA adduct peaks and the reappearance of unmodified RNA.

**Real-time detection of RNA synthesis**

*In vitro* transcription reactions were performed using the HiScribe T7 High Yield RNA Synthesis Kit (NEB) according to the manufacturer’s protocol, with minor modifications. Reactions (20 μL) were assembled on ice in sterile 200 μL PCR tubes in the following order: nuclease-free water, 2 μL of 10× reaction buffer, 2 μL of 100 mM nucleoside triphosphate (NTP) mix (10 mM final per NTP), 1 μg of linearized FLuc control plasmid DNA, 1 μL of 0.1 M dithiothreitol (DTT, 5 mM final), 10 μM **RL fluorophore** and 2 μL of T7 RNA polymerase mix. A no-template control (NTC) omitting the DNA template was prepared in parallel. Samples were transferred to a StepOnePlus Real-Time PCR System (Applied Biosystems) and incubated at 37 °C for 1 h. Fluorescence was monitored in real-time using the SYBR Green detection channel.

**PAGE analysis of labeled oligonucleotides**

To visualize labeled oligonucleotides, 10 pmol of RNA or DNA sample (labeled or unlabeled) was mixed with 5 μL of 2× RNA loading dye (NEB) and loaded onto a 12% polyacrylamide gel (PAGE). Electrophoresis was performed in 1× TBE buffer (pH 8.3, Sigma-Aldrich) at 15 W for 1 h. Fluorescence imaging was conducted prior to SYBR Gold staining, after which gels were re-imaged to visualize total oligonucleotide content. Band intensities were analysed using ImageJ (NIH) or FIJI.

**Dot blot analysis of DNA and RNA extracts from labeled cells**

HeLa human cervical cancer cells were cultured in Dulbecco’s Modified Eagle Medium (DMEM) supplemented with 10% fetal bovine serum (FBS) in a humidified incubator at 37 °C with 5% CO₂ until confluence in T25 flasks. Cells were then treated with 500 μM **RL480** for 24 h, with a vehicle-only treatment serving as a control. Following incubation, cells were harvested by scraping and the resulting pellets were washed once with PBS. DNA and RNA were extracted using Zymo DNA and RNA miniprep kits according to the manufacturer’s protocol. The extracted nucleic acids were normalized to 500 ng/μL, and 1 μL of each sample was spotted onto a pre-cut nitrocellulose membrane. Fluorescence imaging was performed, followed by methylene blue staining and subsequent imaging of the membrane.

**Cellular imaging**

HeLa human cervical cancer cells were cultured in Dulbecco’s Modified Eagle Medium (DMEM) supplemented with 10% fetal bovine serum (FBS) in a humidified incubator at 37 °C with 5% CO₂. For imaging experiments, cells were seeded onto glass-bottom culture dishes (FluoroDish, 35 mm diameter, 10 mm well, WPI) at a density of 30,000 cells per dish one day prior to staining. On the day of the experiment, live cells were stained with **RL fluorophores** (20 µM) and incubated at 37 °C for 30 min prior to imaging. To preserve stained samples, cells were fixed with 4% formaldehyde for 10 min at room temperature, washed three times with PBS, and stored at 4 °C. For dyes that are not cell-permeable, a permeabilization step was included by incubating fixed cells with 0.1% Triton X-100 for 10 min, followed by staining with **RL fluorophores** for 30 min at room temperature. Fluorescence images were acquired using a LSM 780 multiphoton laser scanning confocal microscope (Zeiss) or a SP8 X White Light Laser Confocal Microscope (Leica). Image analysis was performed using ImageJ.

**Comparison of RL vs CA fluorophore staining profiles**

HeLa cells were cultured and seeded onto glass-bottom culture dishes as described above. Live cells were stained with 10 μM of either **RL480** or **CA480** for 30 min at 37 °C. Following incubation, live cells were directly imaged without washing using a LSM 780 multiphoton laser scanning confocal microscope (Zeiss) maintained at 37 °C with 5% CO_2_. Image analysis was performed using ImageJ (NIH).

**Arsenite-induced stress granule imaging**

HeLa cells were seeded at a density of 30,000 cells per dish and cultured until approximately 80% confluency. Stress granules were induced by treating cells with 200 μM sodium arsenite for 30 min at 37 °C. Cells were then fixed with 4% formaldehyde for 10 min at room temperature and blocked for 2 h in PBS containing 3% bovine serum albumin (BSA) and 0.3% Triton X-100. Primary immunostaining was performed using anti-G3BP1 antibody (1:400 dilution in antibody buffer, 0.3% BSA and 0.03% Triton X-100 in PBS) with overnight incubation at 4 °C. After three PBS washes, cells were incubated with a secondary antibody (1:1000 dilution) for 1 h at room temperature. **RL fluorophores** (10 μM) were applied for 30 min at room temperature, followed by imaging.

**Cell viability assay**

Cell viability and cytotoxicity were assessed using the LDH-Glo Cytotoxicity Assay (Promega) following the manufacturer’s protocol. Briefly, HeLa cells were seeded at a density of 20,000 cells per well in white opaque 96-well plates and cultured to approximately 80% confluency. Cells were treated with **RL fluorophores** (**RL480** or **RL560**) at concentrations ranging from 0 to 1000 μM and incubated for 4 or 24 h at 37 °C. Wells containing medium only served as background controls. For 100% cytotoxicity controls, 2 μL of 10% Triton X-100 was added to designated wells and incubated for 10 min. After treatment, 2 μL of supernatant from each well was diluted in 48 μL of lactate dehydrogenase (LDH) Storage Buffer. A secondary dilution was performed by mixing 25 μL of the diluted sample with 75 μL of LDH Storage Buffer. Then, 50 μL of this solution was transferred to a new 96-well plate and combined with 50 μL of LDH Detection Reagent. Luminescence was measured after 1 h incubation at room temperature using a microplate reader.

Cytotoxicity was calculated using the following formula:

$$\% Cytotoxicity=100 \times\frac{Sample LDH Release -Negative Control}{Maximum LDH Release -Negative Control}$$

Cell viability was determined as:

$$\% Cell Viability=100 -\% Cytotoxicity$$

SYNTHESIS

**Synthesis of RL fluorophores**

Carboxylic acid precursors of the **RL** fluorophores (**CA420**, **CA450**, **CA480** and **CA560**) were synthesized following previously reported procedures (see Supplementary Information). For **RL** fluorophores, the corresponding carboxylic acid (1.0 equiv.) was reacted with 1,1′-carbonyldiimidazole (CDI, 1.0 equiv.) in dry dimethyl sulfoxide (DMSO) to a final concentration of 1 M. The reaction mixture was stirred at room temperature, and the reaction progress was monitored by ^1^H nuclear magnetic resonance (NMR) spectroscopy until the signals corresponding to the carboxylic acid precursor had significantly diminished or disappeared. The crude product was used directly as a 1 M stock solution without further purification. Final **RL** fluorophores were characterized by ^1^H NMR and ^13^C NMR spectroscopy and electrospray ionization mass spectrometry (ESI-MS).

**Synthesis of CA420 (Pacific Blue)**

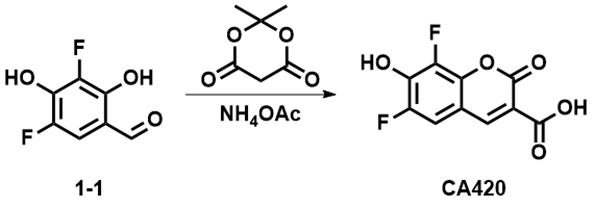

**CA420 (Pacific Blue)** was synthesized according to a previously reported protocol^1^. To a suspension of **compound 1-1** (250 mg, 1.4 mmol) in distilled water (5 mL) was added Meldrum’s acid (228 mg, 1.6 mmol), followed by ammonium acetate (33 mg, 0.4 mmol). The reaction mixture was stirred at room temperature for 5 h. After the addition of 2 M hydrochloric acid (3 mL), the mixture was cooled to 4 °C and stirred for 1 h. The resulting precipitate was collected by filtration, washed with cold water, and dried under high vacuum to yield **CA420 (Pacific Blue)** as a light-yellow solid (220 mg, 91%). ^1^H NMR (400 MHz, DMSO-*d*_6_): *δ* 8.68 (d, *J* = 1.2 Hz, 1H), 7.66 (dd, *J* = 10.5, 2.0 Hz, 1H).

**Synthesis of CA450**

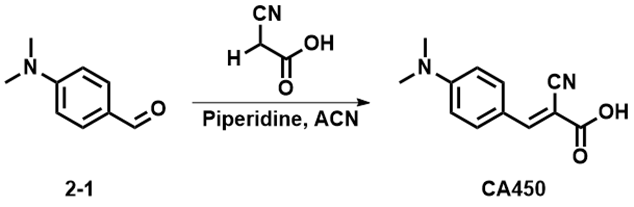

To a solution of **compound 2-1** (500 mg, 3.4 mmol) in acetonitrile, cyanoacetic acid (342 mg, 4.0 mmol) and piperidine (285 mg, 3.4 mmol) were added. The reaction mixture was heated to reflux for 2 h. After cooling to room temperature, the solvent was removed by rotary evaporation. The resulting residue was dissolved in water, acidified to pH 1 with hydrochloric acid, and extracted with dichloromethane. The combined organic layers were washed with brine, dried over anhydrous sodium sulfate, filtered, and concentrated under reduced pressure. The crude product was purified by silica gel column chromatography using ethyl acetate/hexane (1:2) as the eluent to yield **CA450** as a light-yellow solid (210 mg, 29%). ^1^H NMR (400 MHz, DMSO-*d*_6_): *δ* 8.07 (s, 1H), 7.94 (d, *J* = 8.7 Hz, 2H), 6.83 (d, *J* = 8.7 Hz, 2H), 3.08 (s, 6H).

**Synthesis of CA480**

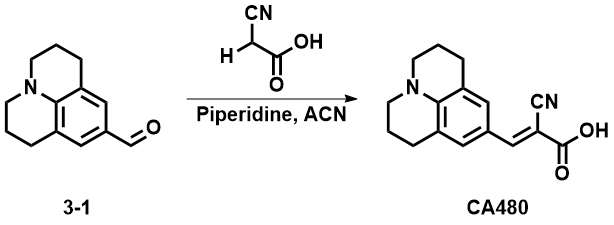

To a solution of **compound 3-1** (1.0 g, 5.0 mmol) in acetonitrile, cyanoacetic acid (854 mg, 10.0 mmol) and piperidine (427 mg, 5.0 mmol) were added. The reaction mixture was heated to reflux for 2 h. After cooling to room temperature, the solvent was removed by rotary evaporation. The resulting residue was dissolved in water, acidified to pH 1 with hydrochloric acid, and extracted with dichloromethane. The combined organic layers were washed with brine, dried over anhydrous sodium sulfate, filtered, and concentrated under reduced pressure. The crude product was purified by silica gel column chromatography using ethyl acetate/hexane (1:2) as the eluent to yield **CA480** as a yellow solid (821 mg, 61%). ^1^H NMR (400 MHz, DMSO-*d*_6_): *δ* 7.87 (s, 1H), 7.50 (s, 2H), 3.34 (s, 4H), 2.67 (t, *J* = 6.2 Hz, 4H), 1.87 (p, *J* = 6.3 Hz, 4H).

**Synthesis of 4-2**

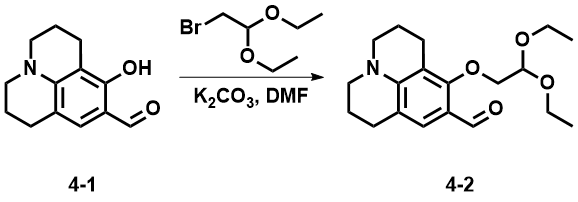

To a stirred solution of 4-(dimethylamino)-2-hydroxybenzaldehyde (3.6 g, 18.4 mmol) and potassium carbonate (7.0 g, 36.8 mmol) in anhydrous dimethylformamide (10 mL), **compound 4-1** (2.0 g, 9.2 mmol) was added. The reaction mixture was heated to 100 °C and stirred overnight. After cooling to room temperature, the mixture was concentrated under reduced pressure. The resulting residue was extracted with ethyl acetate, and the combined organic layers were dried over anhydrous sodium sulfate, filtered, and concentrated. The crude product was purified by silica gel column chromatography using ethyl acetate/hexane (1:5) as the eluent to yield **compound 4-2** as a solid (1.7 g, 55%). ^1^H NMR (400 MHz, DMSO-*d*_6_): *δ* 10.03 (s, 1H), 10.03 (s, 1H), 7.31 (s, 1H), 7.31 (s, 1H), 4.85 (t, *J* = 5.1 Hz, 1H), 4.85 (t, *J* = 5.1 Hz, 1H), 3.91 (d, *J* = 5.2 Hz, 2H), 3.75 (dq, *J* = 9.3, 7.1 Hz, 2H), 3.63 (dq, *J* = 9.3, 7.0 Hz, 2H), 3.30 – 3.23 (m, 4H), 2.77 (t, *J* = 6.3 Hz, 2H), 2.69 (t, *J* = 6.2 Hz, 2H), 1.97 – 1.85 (m, 4H), 1.24 (t, *J* = 7.1 Hz, 6H).

**Synthesis of 4-3**

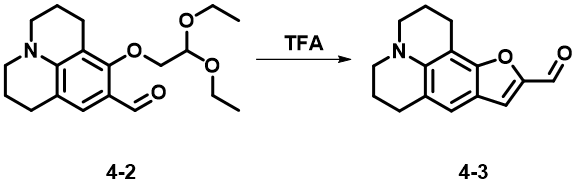

To a mixture of tetrahydrofuran (3 mL) and water (3 mL), **compound 4-1** (236 mg, 0.7 mmol) was added, followed by trifluoroacetic acid (1.1 mL). The reaction mixture was stirred at room temperature for 5 h, then poured into saturated aqueous sodium bicarbonate. The aqueous layer was extracted with ethyl acetate, and the combined organic extracts were dried over anhydrous sodium sulfate. After removal of the solvent under reduced pressure, the crude residue was purified by silica gel column chromatography using ethyl acetate/hexane (1:9) as the eluent to yield **compound 4-2** as a yellow oil (150 mg, 17%). ^1^H NMR (400 MHz, CDCl_3_): *δ* 9.57 (s, 1H), 7.34 (s, 1H), 7.09 (s, 1H), 3.26 (q, *J* = 5.8 Hz, 4H), 2.98 (t, *J* = 6.5 Hz, 2H), 2.84 (t, *J* = 6.4 Hz, 2H), 2.00 (ddt, *J* = 15.7, 11.5, 6.2 Hz, 4H).

**Synthesis of CA560**

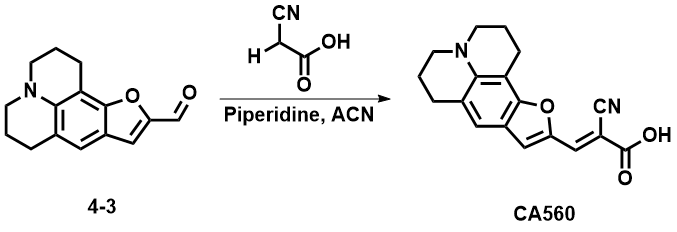

To a solution of **compound 4-3** (93 mg, 0.4 mmol) in acetonitrile, cyanoacetic acid (66 mg, 0.8 mmol) and piperidine (33 mg, 0.4 mmol) were added. The reaction mixture was heated to reflux for 3 h. After cooling to room temperature, the solvent was removed by rotary evaporation. The resulting residue was dissolved in water, acidified to pH 1 with HCl, and filtered. The solid was washed with water and dried under high vacuum to yield **CA560** as a purple solid (105 mg, 88%). ^1^H NMR (400 MHz, DMSO-*d*_6_): *δ* 7.93 (s, 1H), 7.57 (s, 1H), 7.13 (s, 1H), 3.29 (dt, *J* = 11.1, 5.6 Hz, 4H), 2.81 (dt, *J* = 31.4, 6.3 Hz, 4H), 1.98 – 1.83 (m, 4H); ^13^C NMR (100 MHz, DMSO-*d*_6_): *δ* 155.08 (s), 147.54 (s), 144.16 (s), 135.96 (s), 121.02 (s), 119.56 (s), 118.82 (s), 118.56 (s), 116.73 (s), 102.09 (s), 49.93 (s), 49.61 (s), 28.10 (s), 21.71 (s), 21.49 (s), 20.58 (d, *J* = 7.8 Hz); MS (ESI): C_18_H_16_N_2_O_3_ [M + H]^+^, *m*/*z* calcd 308.1161, found 390.1287.

**Synthesis of RL fluorophores**

For **RL fluorophores**, the corresponding carboxylic acid (1.0 equiv.) was reacted with 1,1′-carbonyldiimidazole (CDI, 1.0 equiv.) in dry DMSO to afford a final concentration of 1 M. The resulting solution was stirred at room temperature and the reaction progress was monitored by NMR spectroscopy until the signals corresponding to the carboxylic acid precursor had significantly diminished or disappeared. The crude product was used directly as a 1 M stock solution without further purification. Final **RL fluorophores** were characterized by ^1^H NMR spectroscopy, ^13^C NMR spectroscopy, and electrospray ionization mass spectrometry (ESI-MS).

**Characterization of RL420**

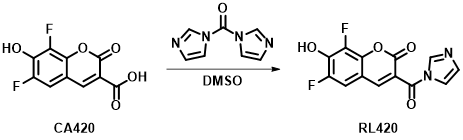

^1^H NMR (400 MHz, DMSO-*d*_6_): *δ* 8.42 (d, *J* = 1.9 Hz, 1H), 8.24 (t, *J* = 1.1 Hz, 1H), 8.22 (d, *J* = 1.8 Hz, 1H), 7.61 (t, *J* = 1.4 Hz, 1H), 7.01 (dd, *J* = 1.6, 0.9 Hz, 1H); ^13^C NMR (100 MHz, DMSO-*d*_6_): *δ* 165.37 (s), 163.61 (s), 158.84 (s), 158.62 (s), 149.36 (s), 148.78 (s), 138.82 (s), 129.73 (s), 118.25 (s), 110.10 (s), 110.04 (s), 109.90 (s), 109.84 (s); MS (ESI): C_13_H_6_F_2_N_2_O_4_ [M + H]^+^, *m*/*z* calcd 292.0296, found 293.1360.

**Characterization of RL450**

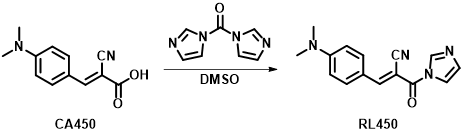

^1^H NMR (400 MHz, DMSO-*d*_6_): 8.42 – 8.38 (m, 1H), 8.17 (s, 1H), 8.07 (d, *J* = 9.3 Hz, 2H), 7.76 (t, *J* = 1.4 Hz, 1H), 7.14 (dd, *J* = 1.8, 0.9 Hz, 1H), 6.91 (d, *J* = 9.4 Hz, 2H), 3.15 (s, 6H); ^13^C NMR (100 MHz, DMSO-*d*_6_): *δ* 161.76 (s), 157.14 (s), 154.52 (s), 137.73 (s), 134.88 (s), 129.95 (s), 121.37 (s), 118.59 (s), 118.36 (s), 118.23 (s), 92.11 (s), 40.43 (s); MS (ESI): C_15_H_14_N_4_O [M + H]^+^, *m*/*z* calcd 266.1168, found 267.0578.

**Characterization of RL480**

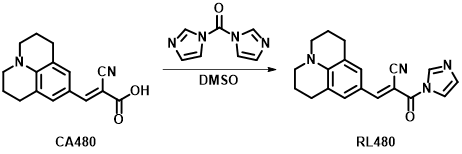

^1^H NMR (400 MHz, DMSO-*d*_6_): *δ* 8.35 (s, 1H), 7.95 (s, 1H), 7.72 (d, *J* = 1.6 Hz, 1H), 7.66 (s, 1H), 7.48 (s, 1H), 7.11 (d, *J* = 1.6 Hz, 1H), 3.41 (t, *J* = 5.9 Hz, 4H), 2.71 – 2.67 (m, 4H), 1.92 – 1.85 (m, 4H); ^13^C NMR (100 MHz, DMSO-*d*_6_): *δ* 161.94 (s), 156.20 (s), 149.20 (s), 137.53 (s), 129.75 (s), 121.05 (s), 120.47 (s), 118.90 (s), 118.33 (s), 117.76 (s), 88.86 (s), 49.80 (s), 26.93 (s), 20.34 (s); MS (ESI): C_19_H_18_N_4_O [M + H]^+^, *m*/*z* calcd 318.1481, found 319.3224.

**Characterization of RL560**

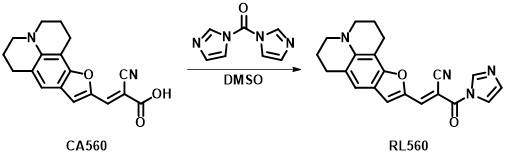

^1^H NMR (400 MHz, DMSO-*d*_6_): *δ* 8.41 (s, 1H), 8.03 (s, 1H), 7.81 (s, 2H), 7.23 (s, 1H), 7.13 (d, *J* = 1.4 Hz, 1H), 2.87 (t, *J* = 6.3 Hz, 2H), 2.78 (t, *J* = 6.2 Hz, 2H), 1.98 – 1.86 (m, 4H); ^13^C NMR (100 MHz, DMSO-*d*_6_): *δ* 161.32 (s), 156.83 (s), 147.06 (s), 146.37 (s), 139.10 (s), 137.64 (s), 129.86 (s), 122.53 (s), 120.46 (s), 118.34 (s), 117.53 (s), 117.20 (s), 100.61 (s), 90.45 (s), 49.74 (s), 49.41 (s), 27.69 (s), 20.94 (s), 19.93 (s), 19.70 (s); MS (ESI): C_21_H_18_N_4_O_2_ [M + H]^+^, *m*/*z* calcd 358.1430, found 359.2881.

Supplementary Figures

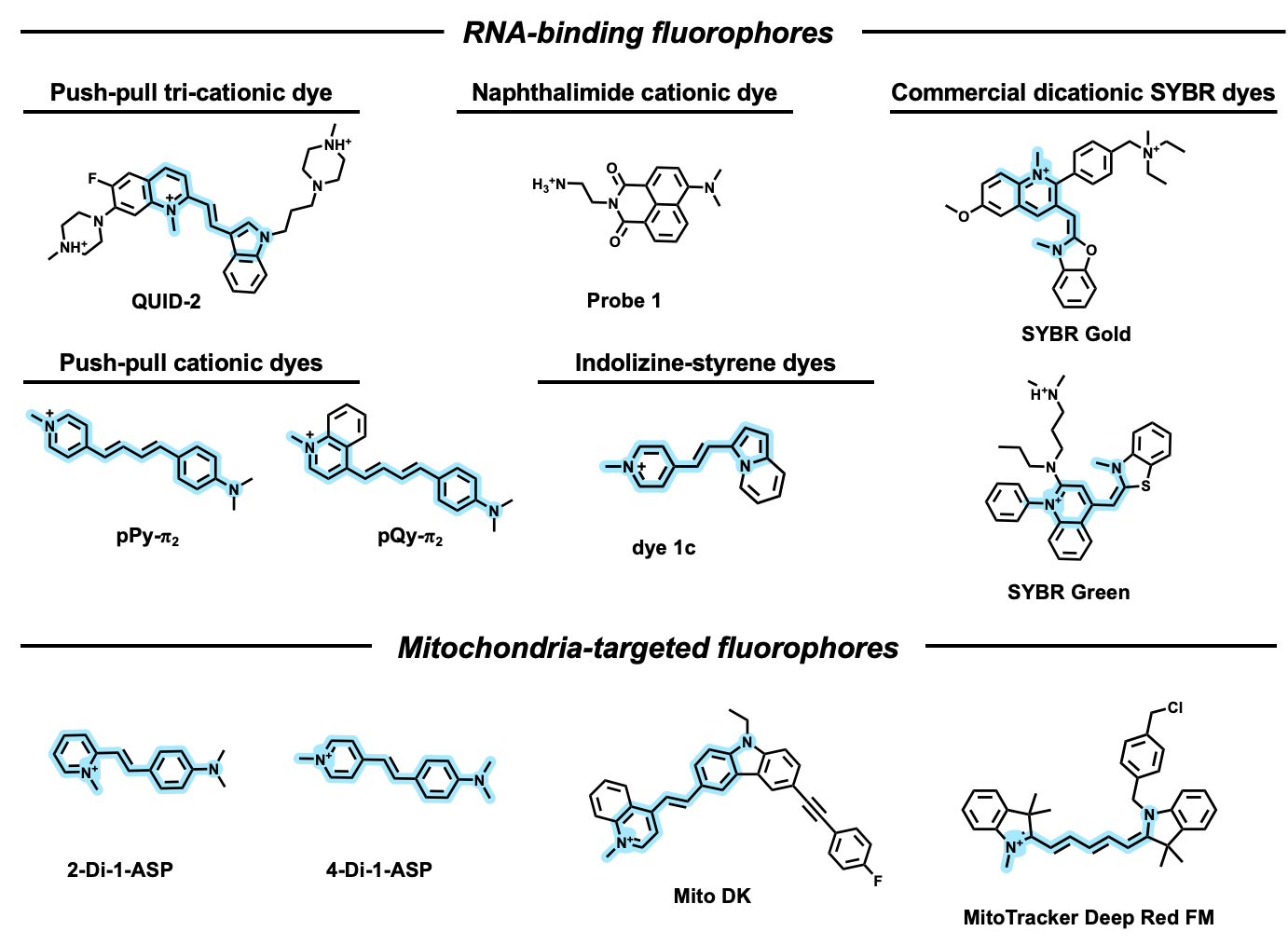

Figure S1. Structures of previously reported non-covalent RNA-binding fluorophores and structurally related mitochondria dyes. (top) Representative structures of published and/or commercial dyes that bind RNA through non-covalent interactions. Commonly employed push-pull pyridinium structure is highlighted. While these dyes exhibit fluorescence enhancement upon nucleic acid binding, they show limited selectivity over DNA. (bottom) Structurally related cationic mitochondrial dyes also possess extended π-systems and delocalized amine push-pull positive charges, enabling organelle targeting and fluorescence enhancement in cell imaging. The close structural similarity between some mitochondrial dyes and non-covalent RNA dyes highlights potential risks of using such cationic aromatic motifs to associate with RNA.

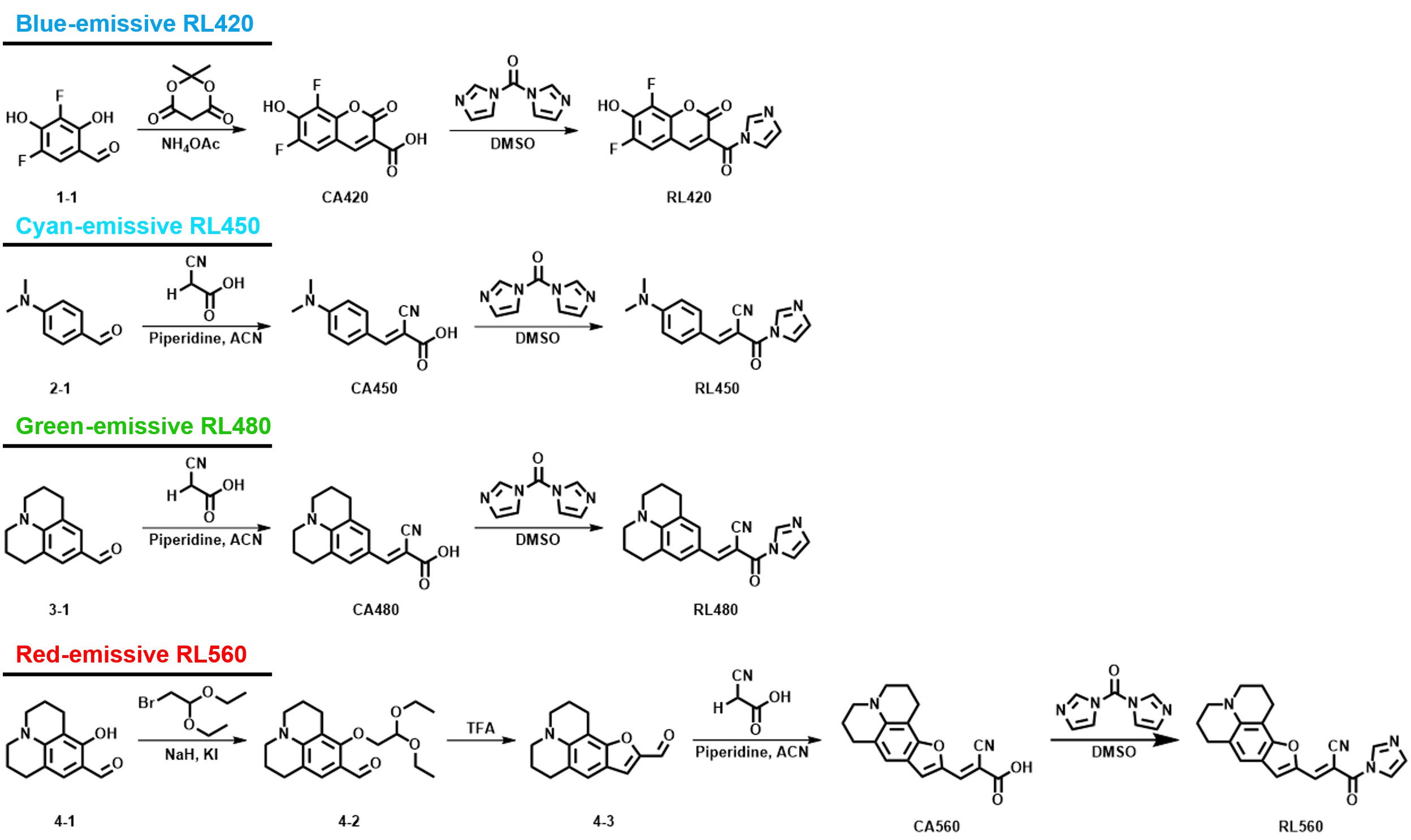

Figure S2. Synthetic scheme for the preparation of RL fluorophores. Synthesis of wavelength-engineered RNA covalently selective RL platform (RL420, RL450, RL480 and RL560) via CDI-mediated activation of corresponding carboxylic acid precursors (CA420, CA450, CA480 and CA560).

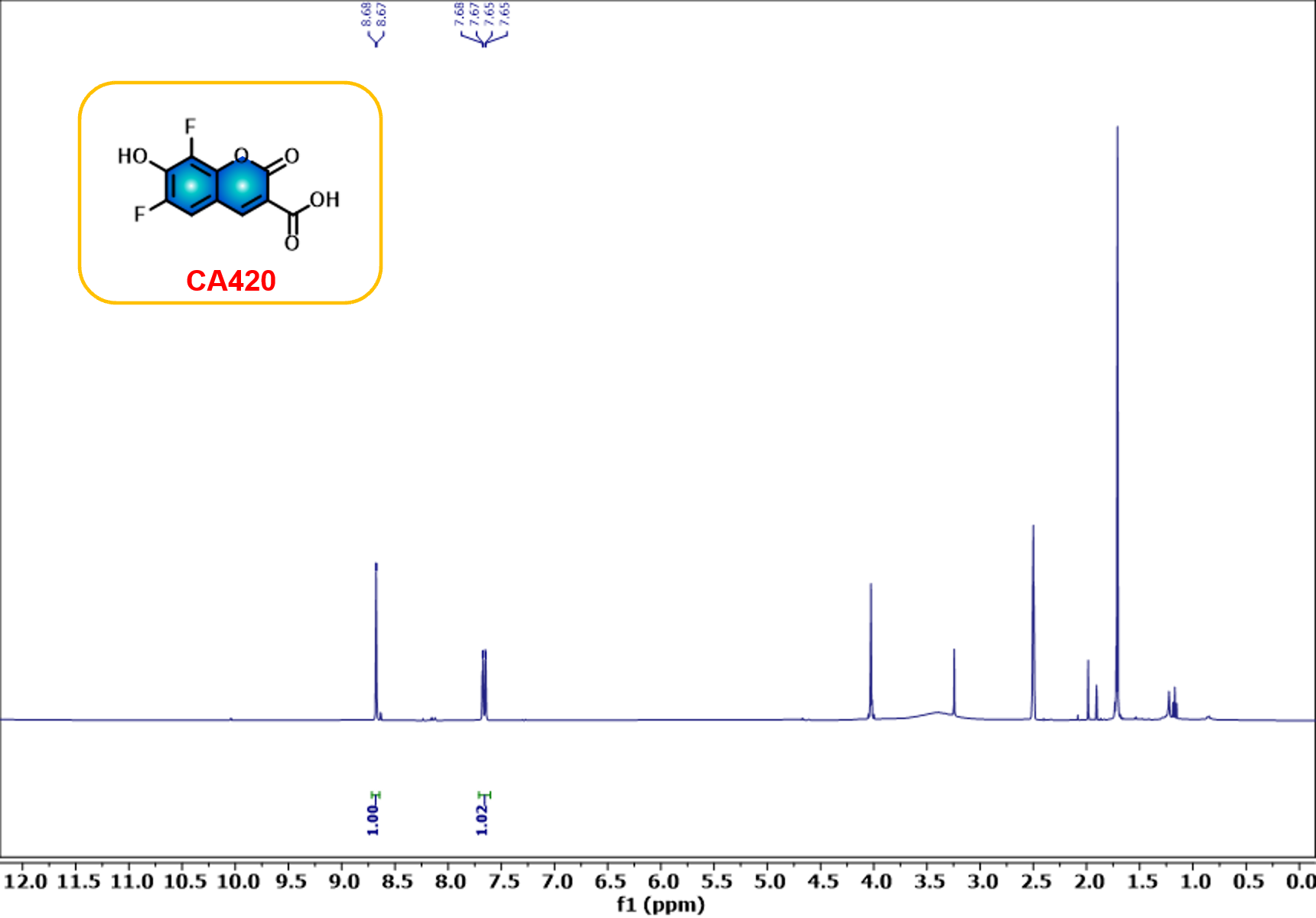

Figure S3. ^1^H NMR spectrum (400 MHz) of CA420 in DMSO-*d*_6_.

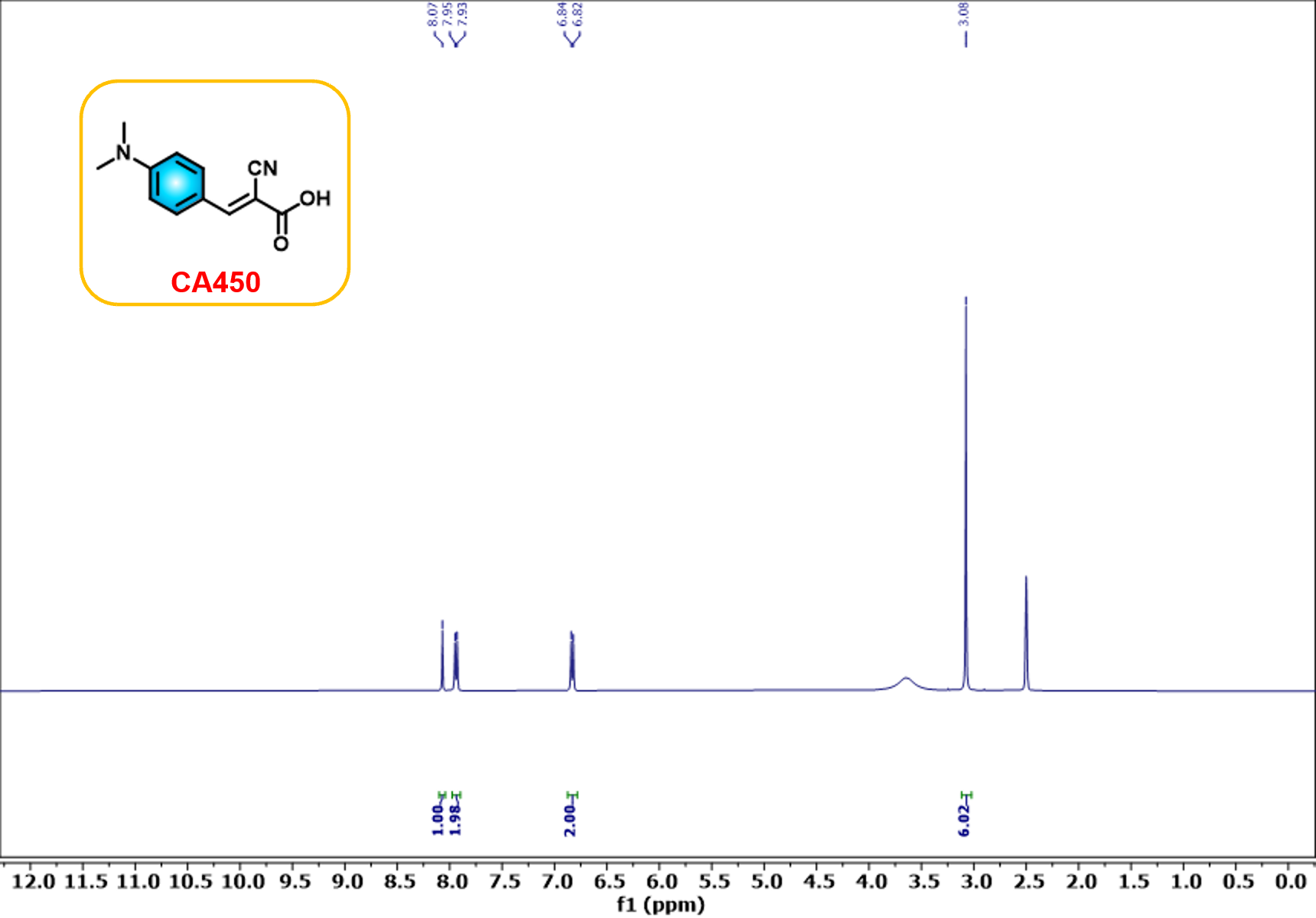

Figure S4. ^1^H NMR spectrum (400 MHz) of CA450 in DMSO-*d*_6_.

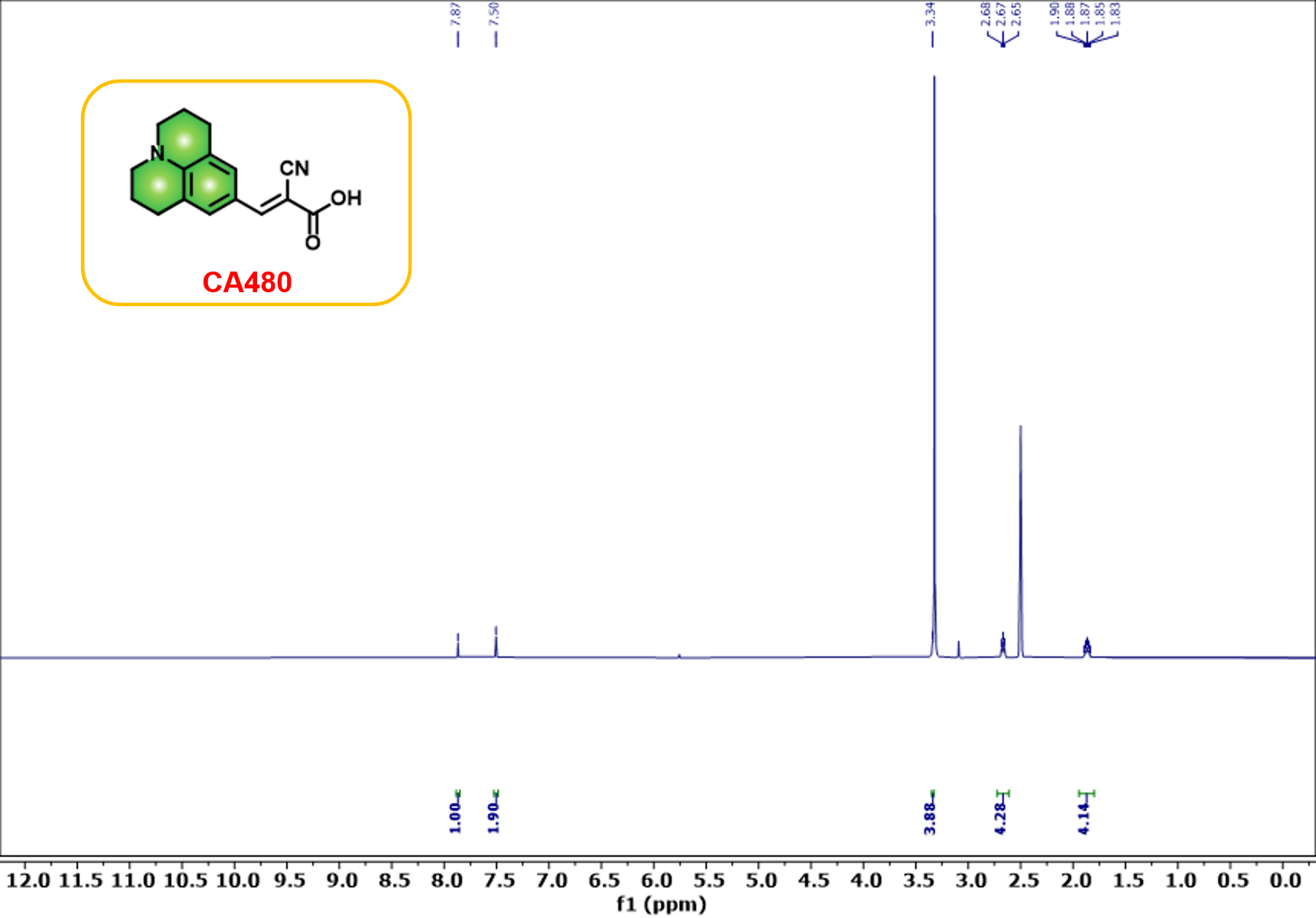

Figure S5. ^1^H NMR spectrum (400 MHz) of CA480 in DMSO-*d*_6_.

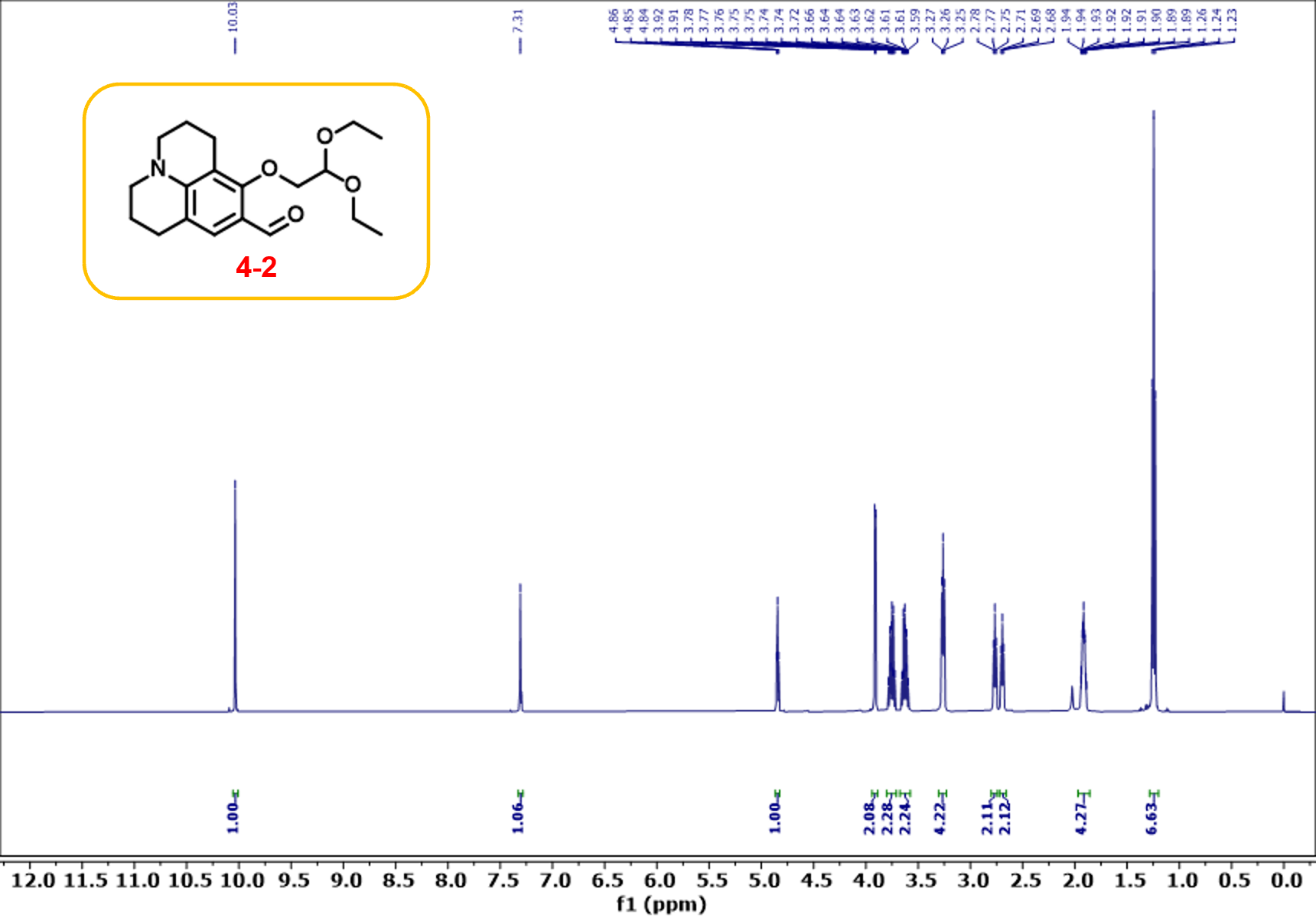

Figure S6. ^1^H NMR spectrum (400 MHz) of Compound 4-2 in DMSO-*d*_6_.

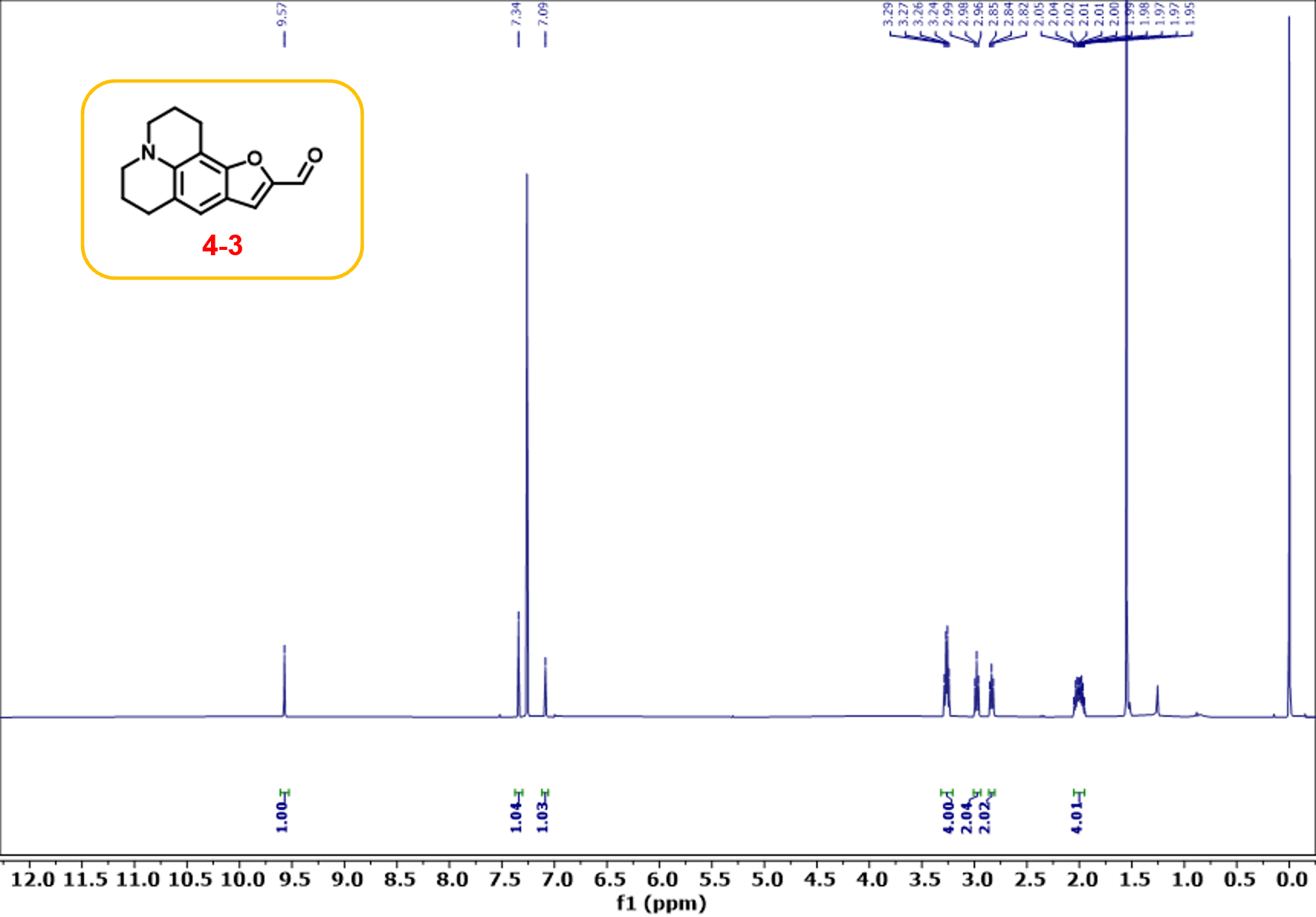

Figure S7. ^1^H NMR spectrum (400 MHz) of Compound 4-3 in CDCl_3_.

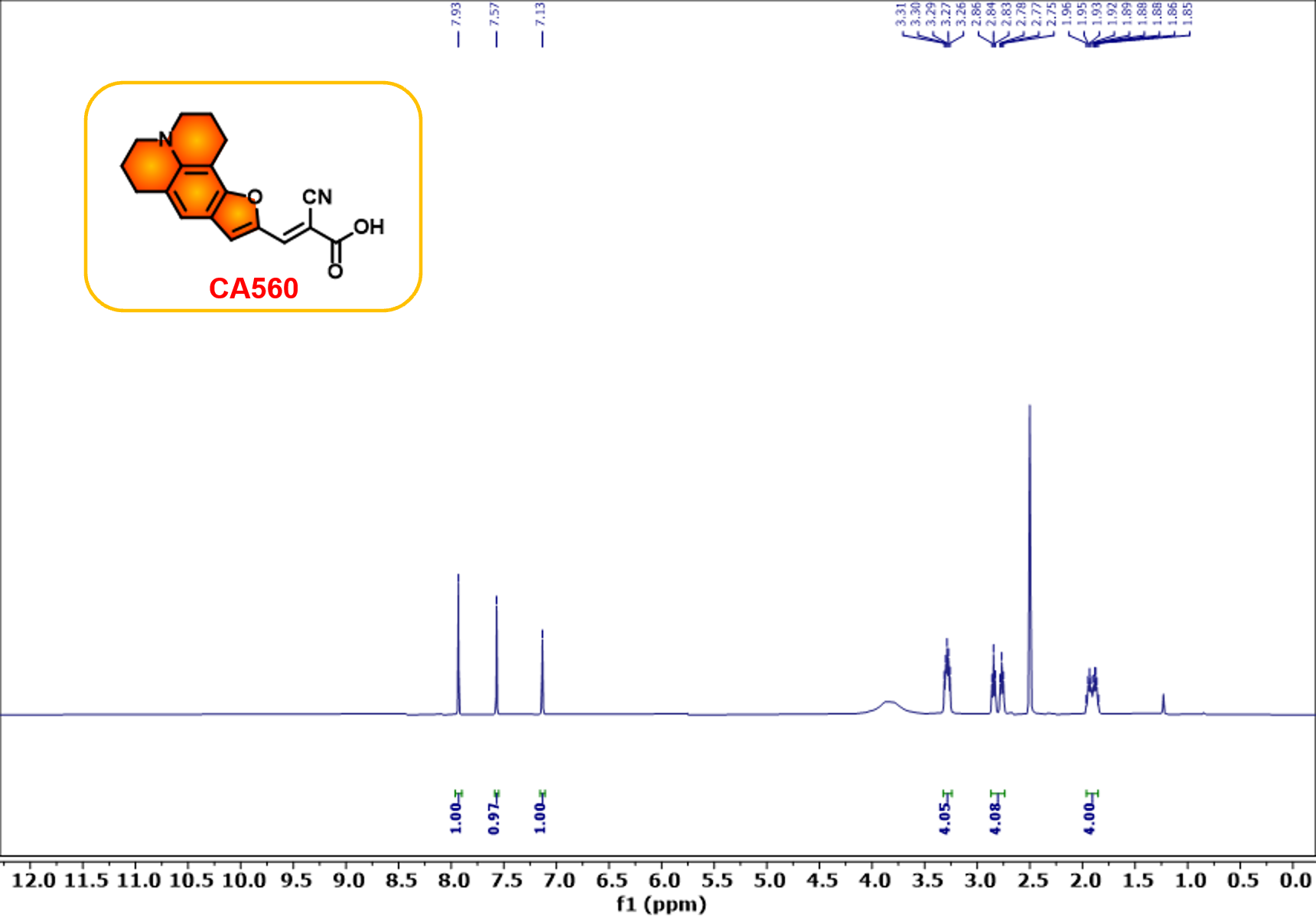

Figure S8. ^1^H NMR spectrum (400 MHz) of CA560 in DMSO-*d*_6_.

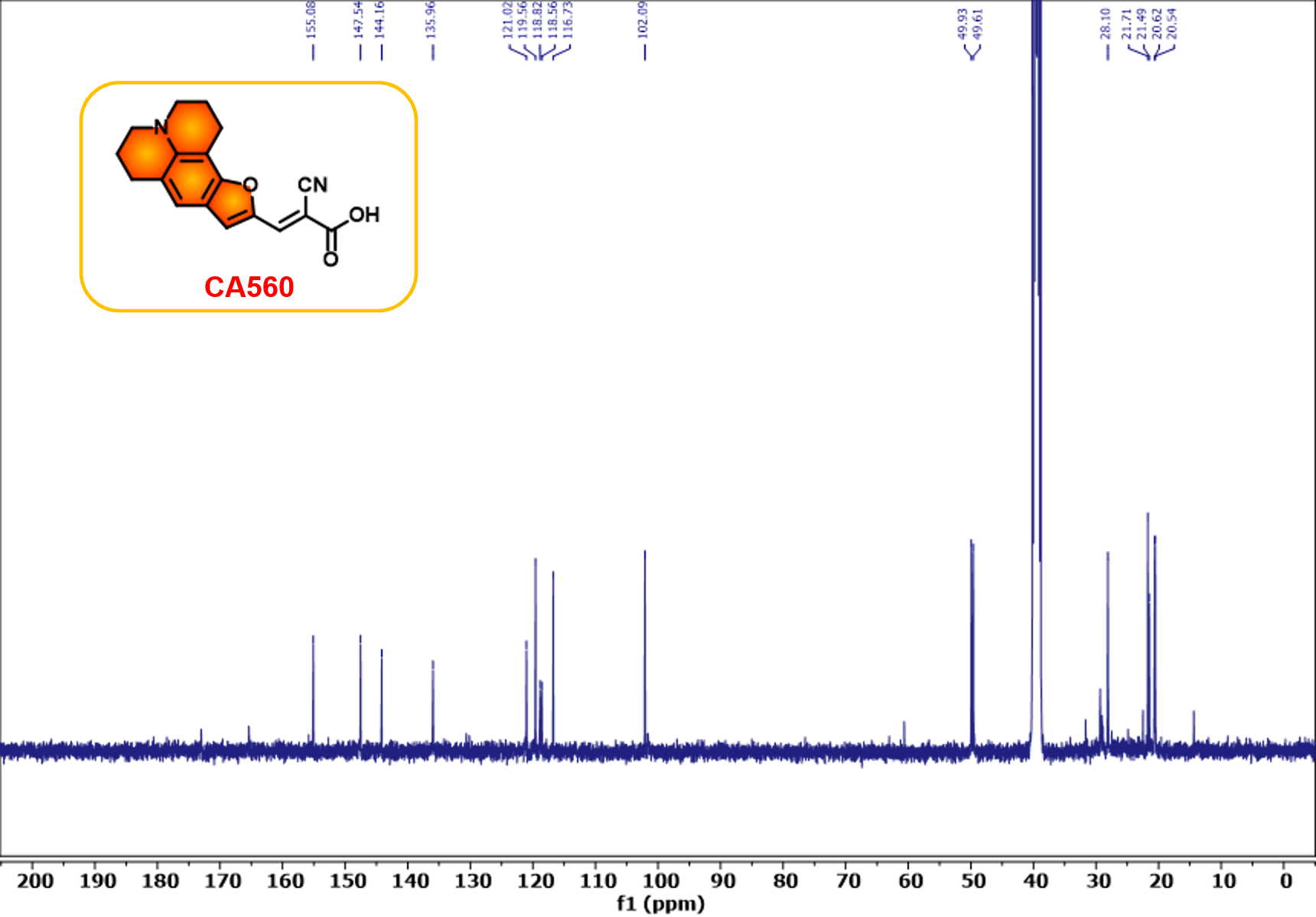

Figure S9. ^13^C NMR spectrum (100 MHz) of CA560 in DMSO-*d*_6_.

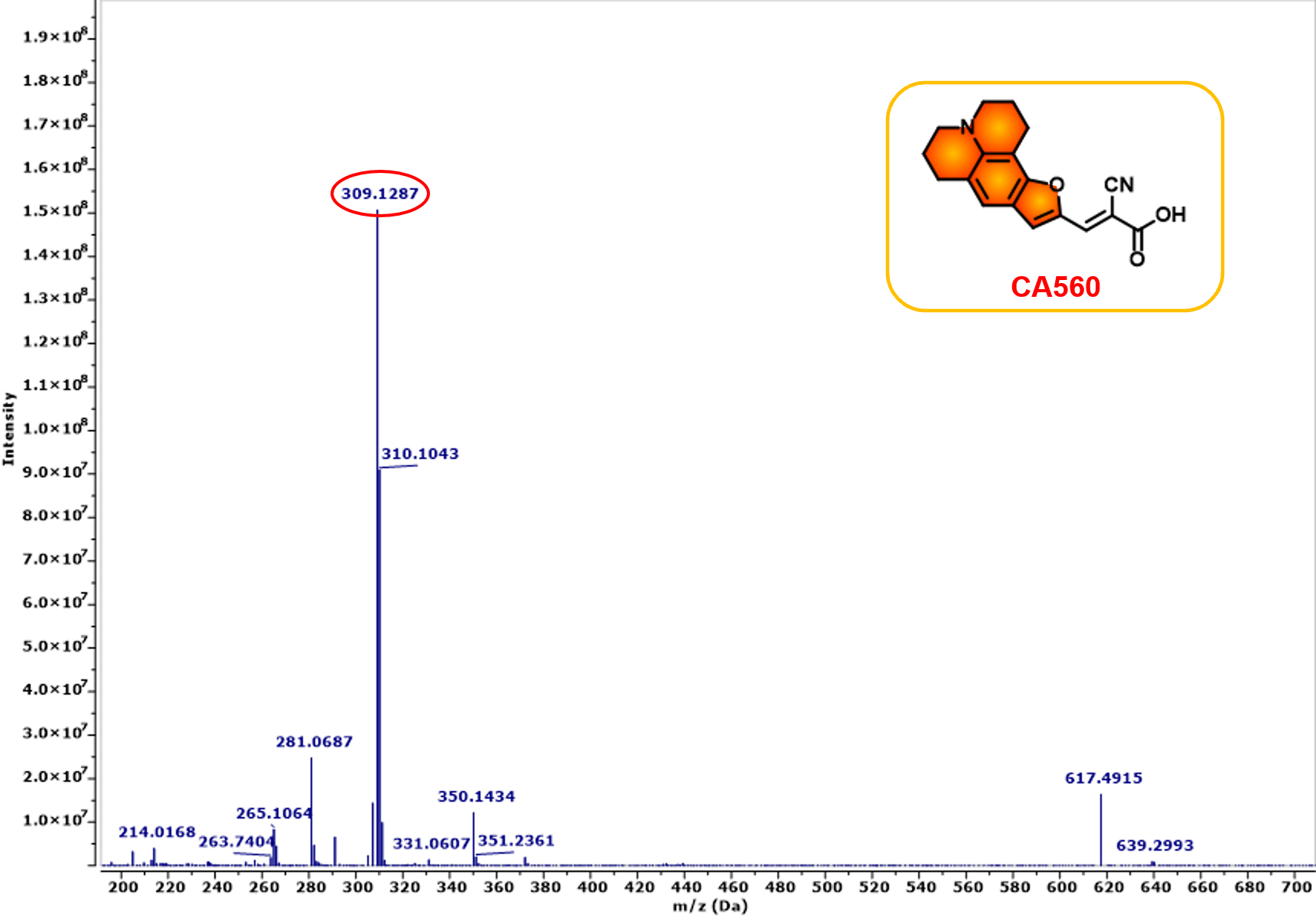

Figure S10. ESI-MS spectrum of CA560.

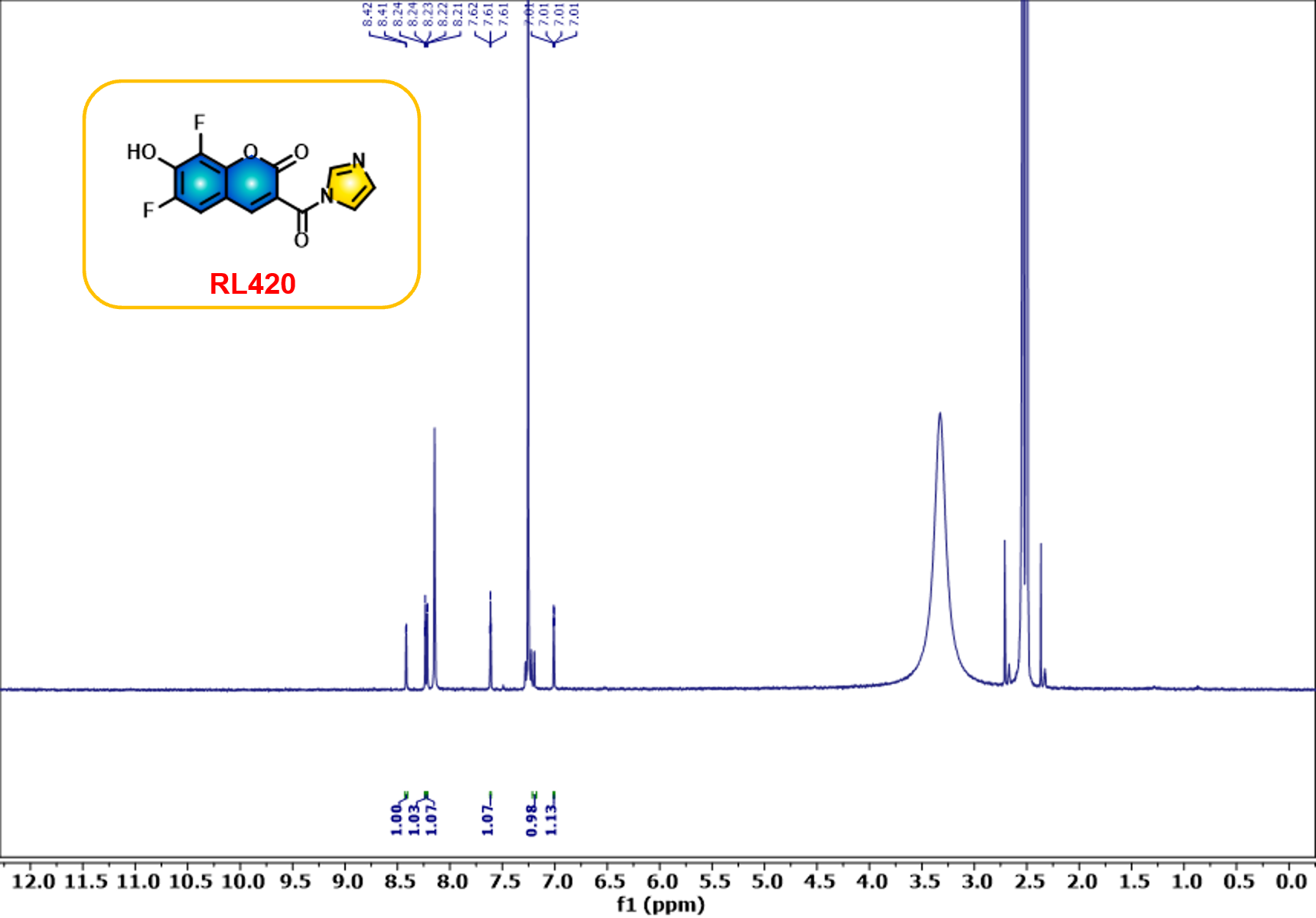

Figure S11. ^1^H NMR spectrum (400 MHz) of RL420 in DMSO-*d*_6_.

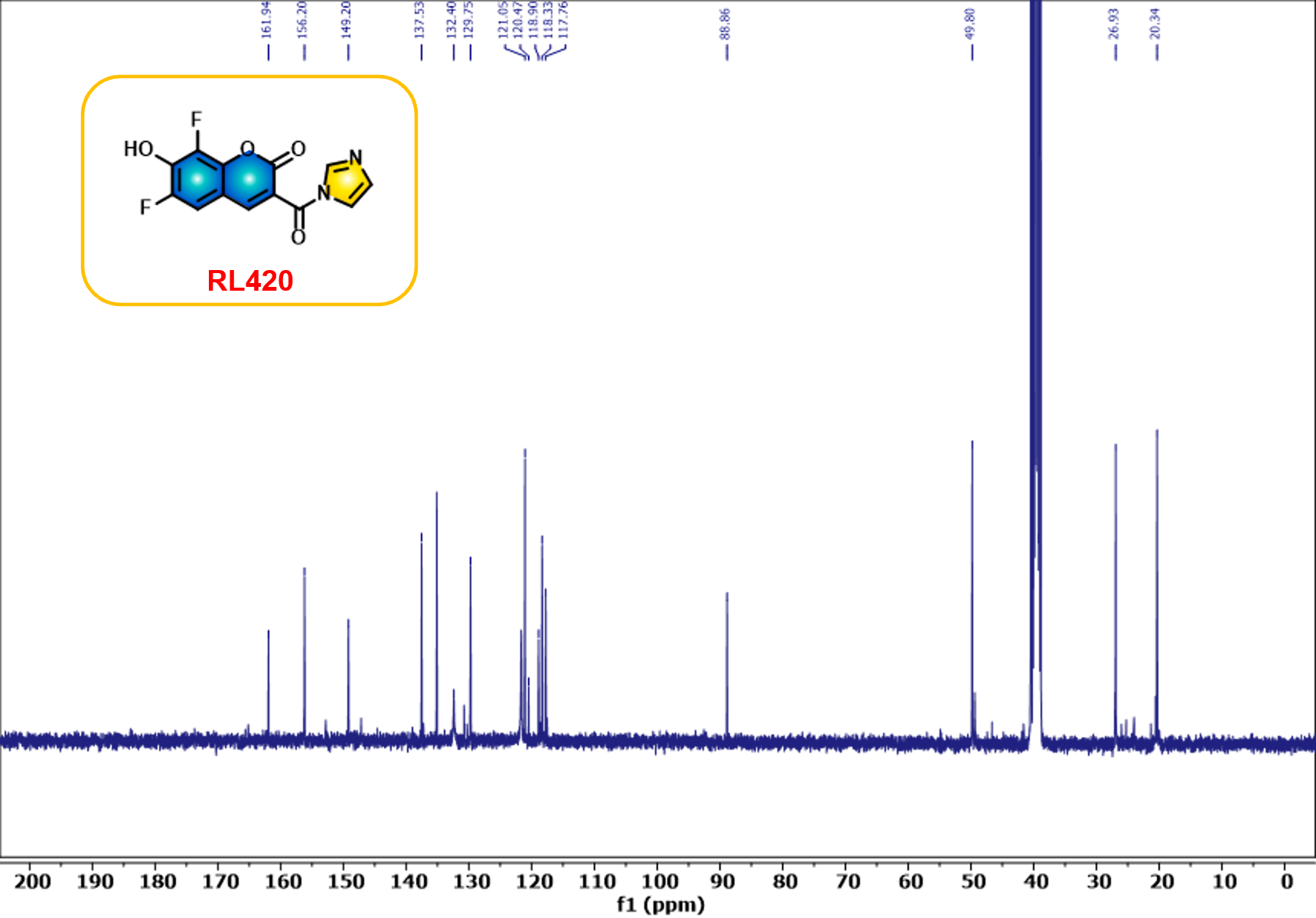

Figure S12. ^13^C NMR spectrum (100 MHz) of RL420 in DMSO-*d*_6_.

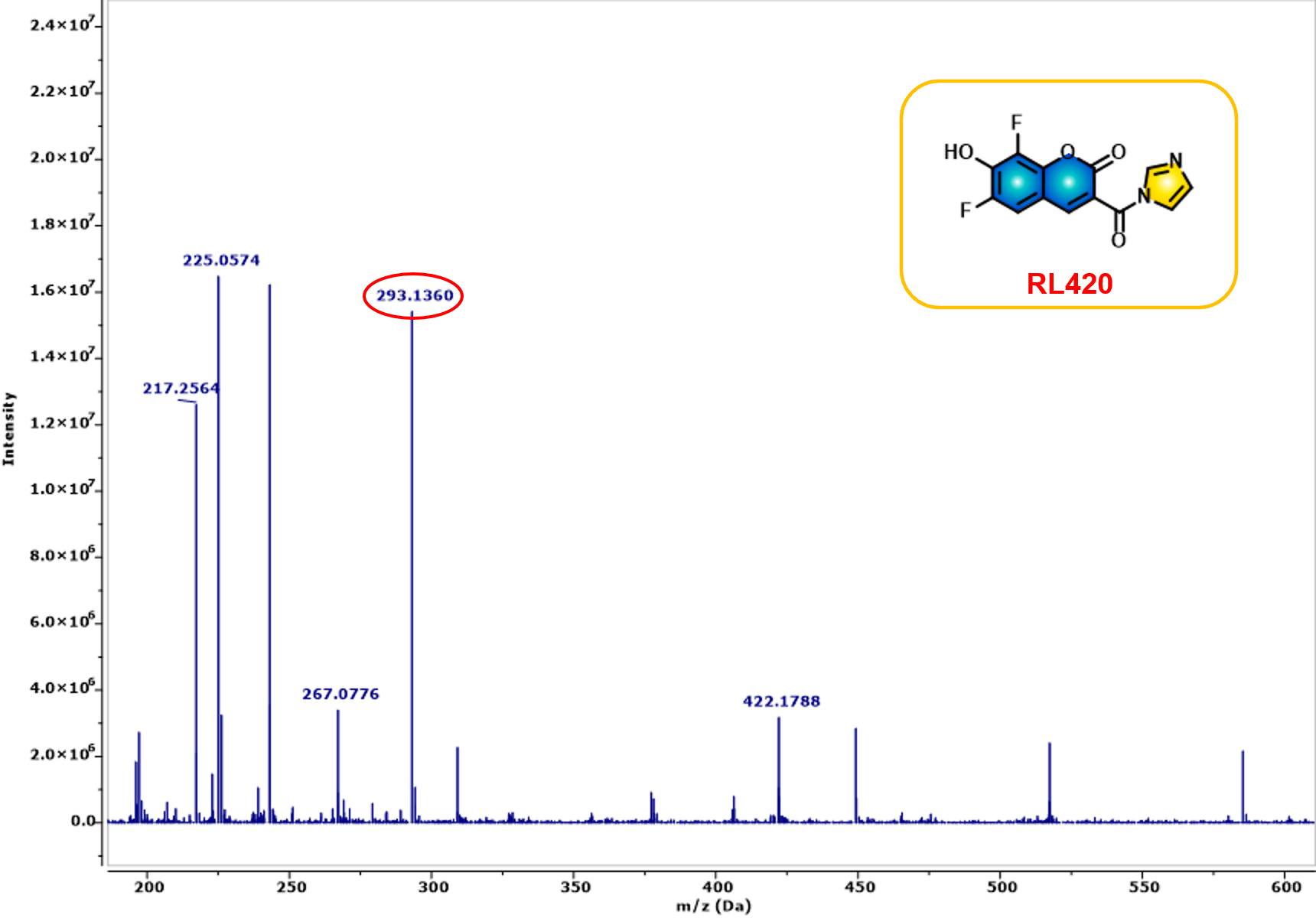

Figure S13. ESI-MS spectrum of RL420.

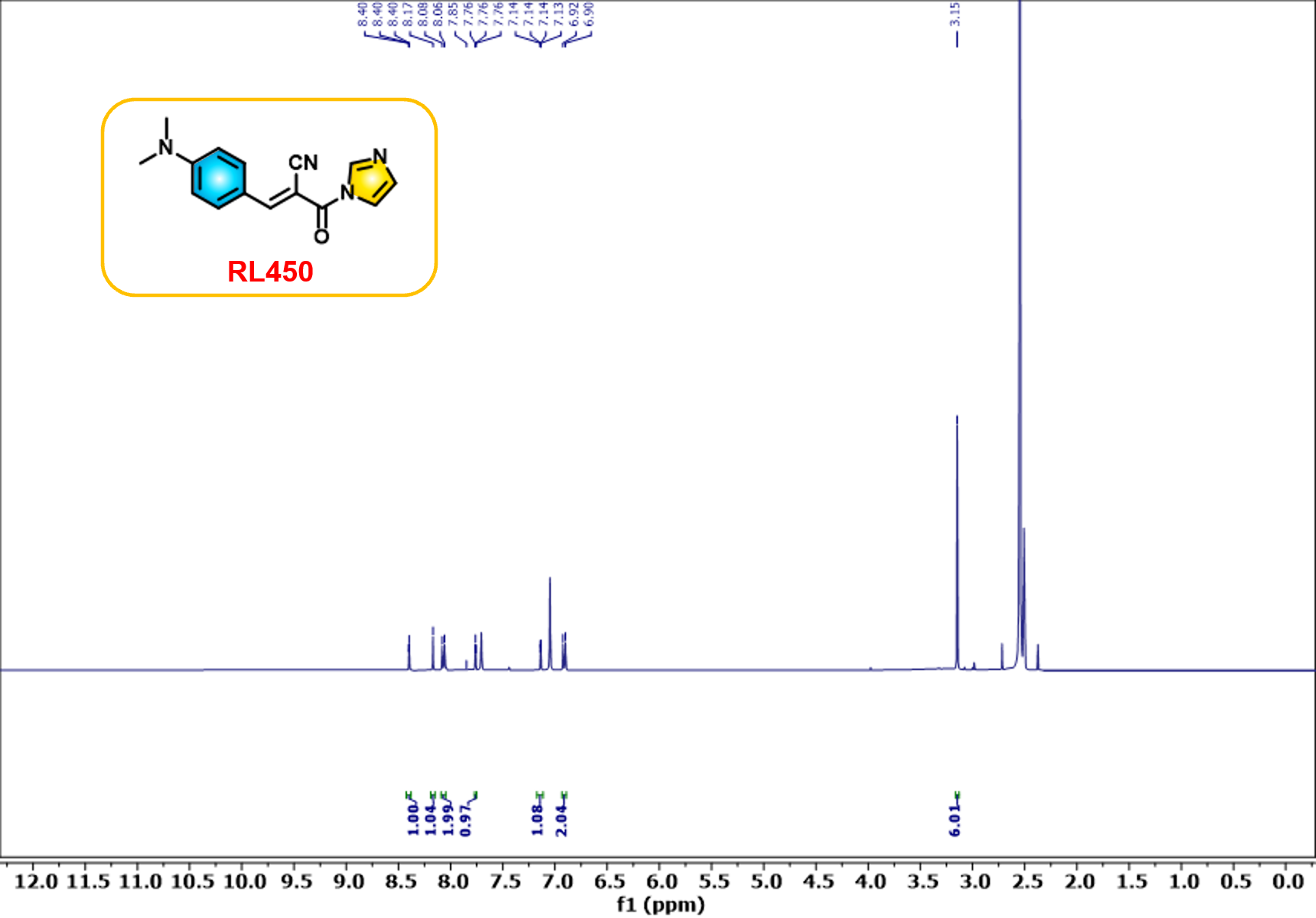

Figure S14. ^1^H NMR spectrum (400 MHz) of RL450 in DMSO-*d*_6_.

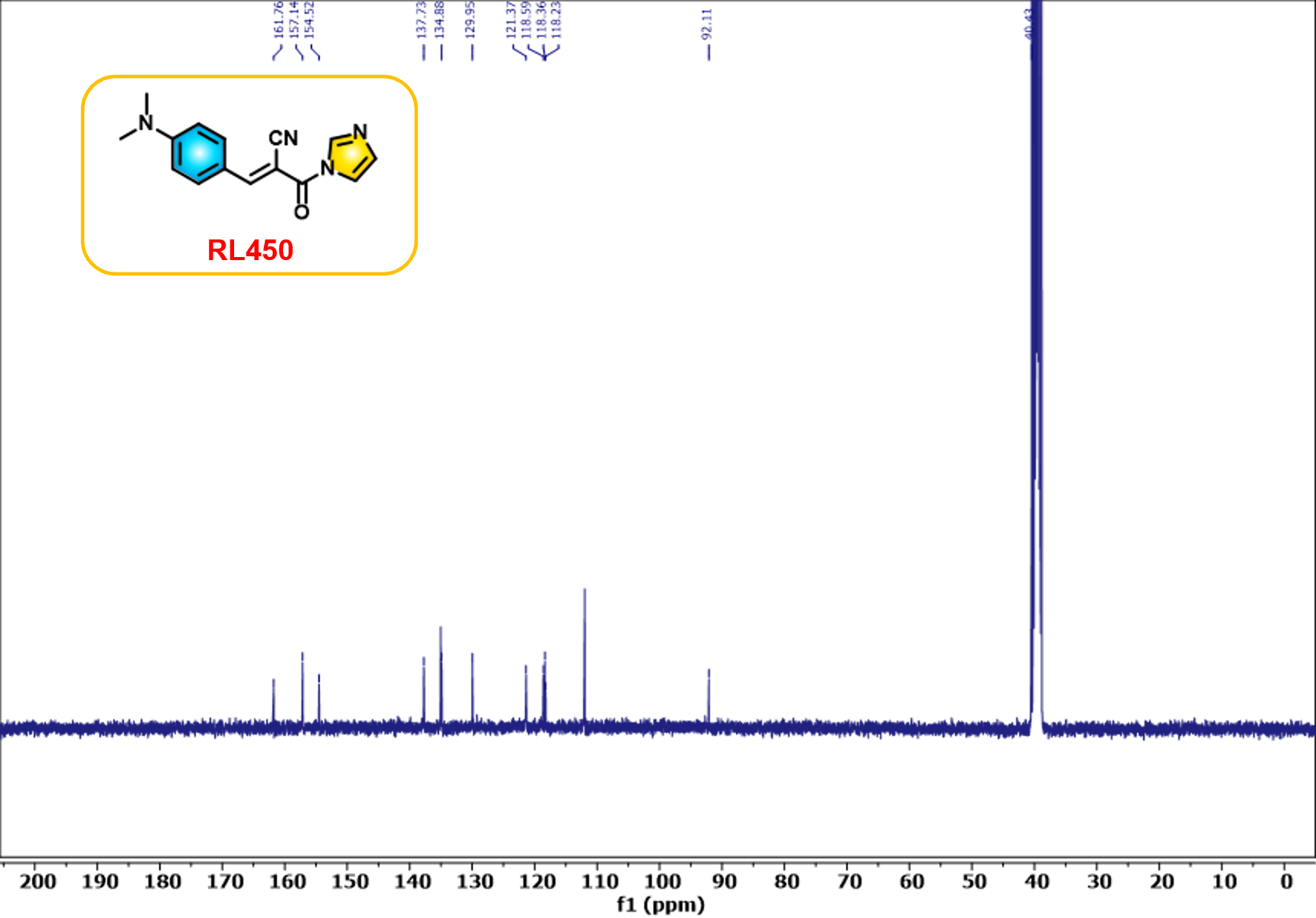

Figure S15. ^13^C NMR spectrum (400 MHz) of RL450 in DMSO-*d*_6_.

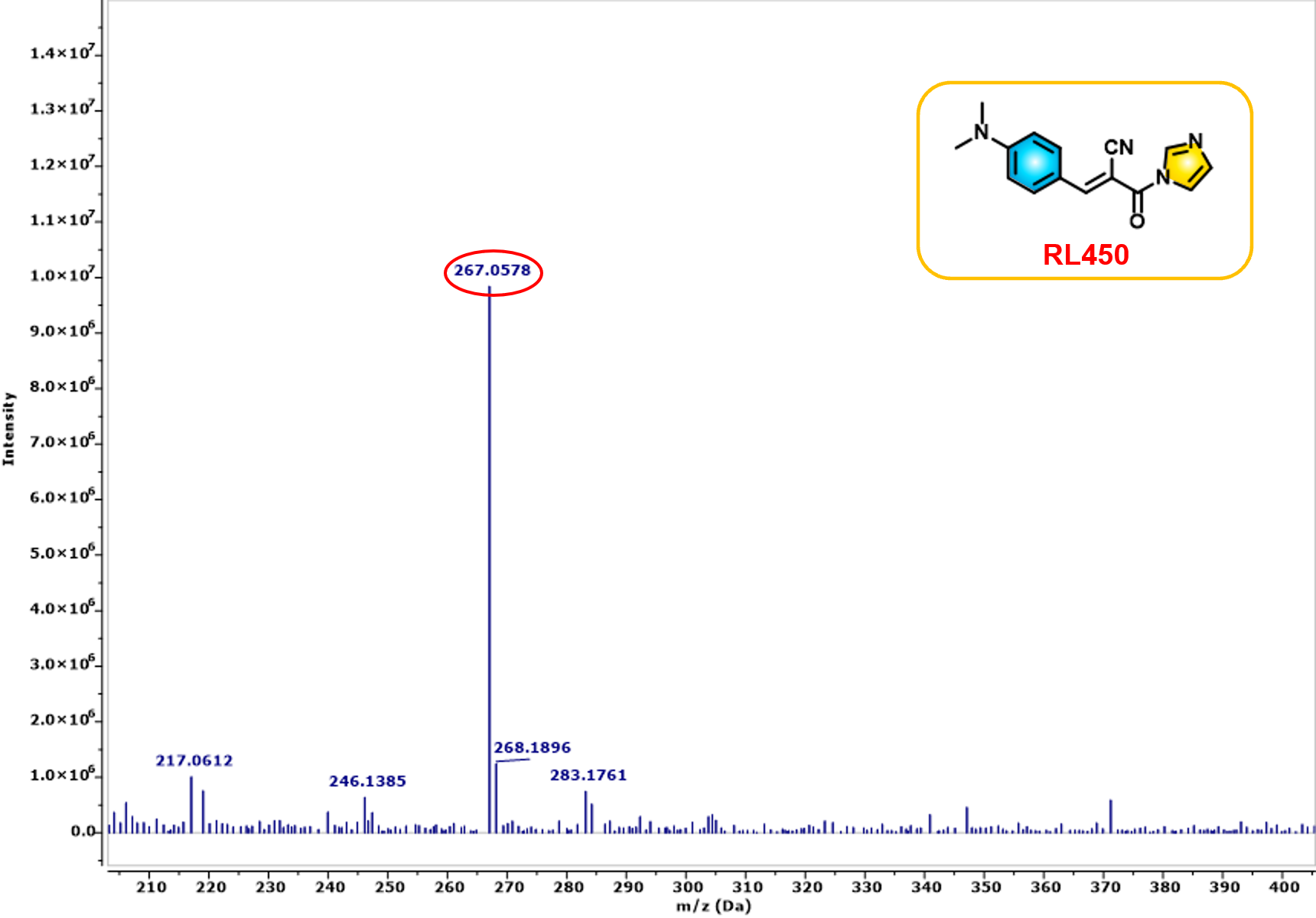

Figure S16. ESI-MS spectrum of RL450.

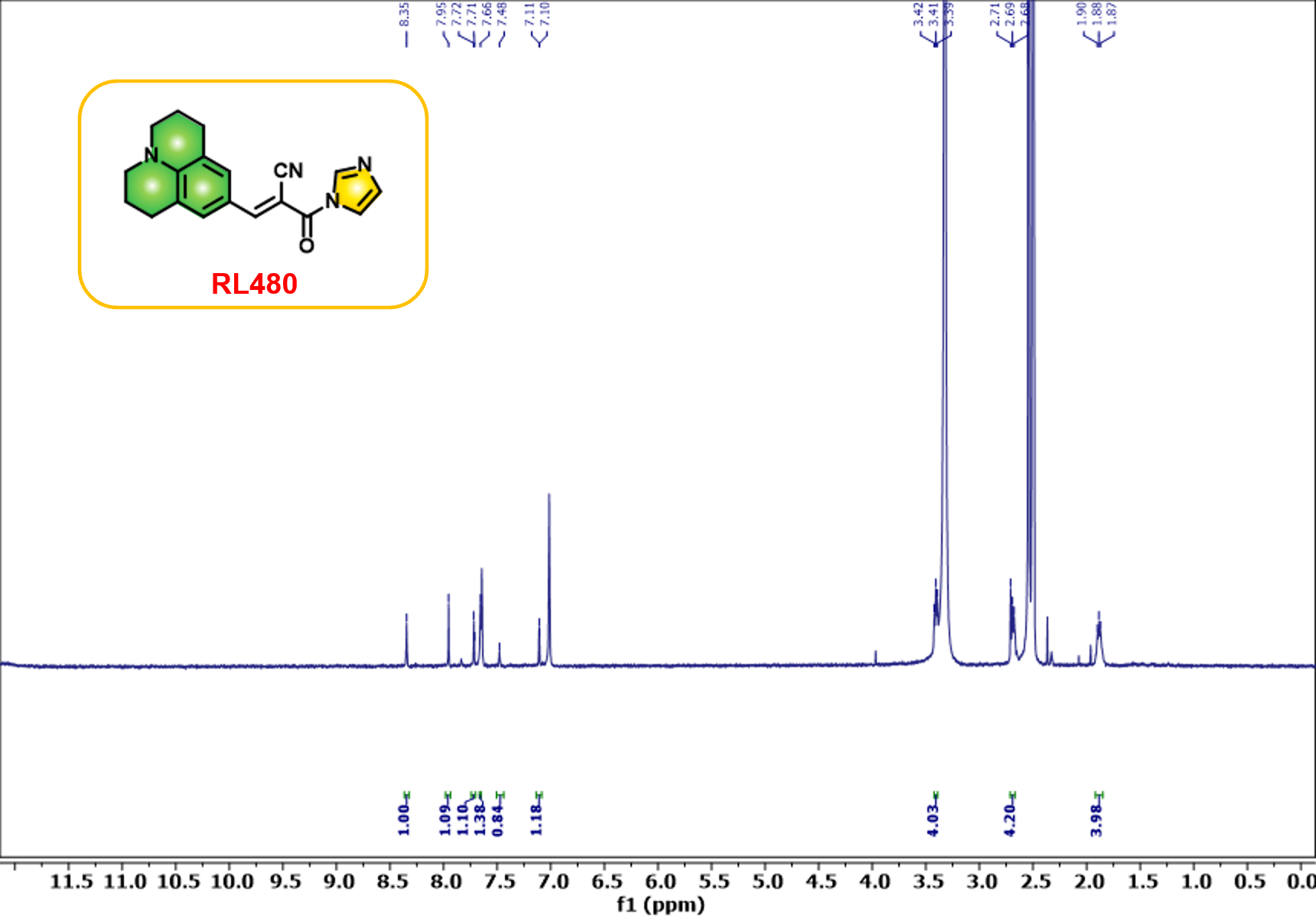

Figure S17. ^1^H NMR spectrum (400 MHz) of RL480 in DMSO-*d*_6_.

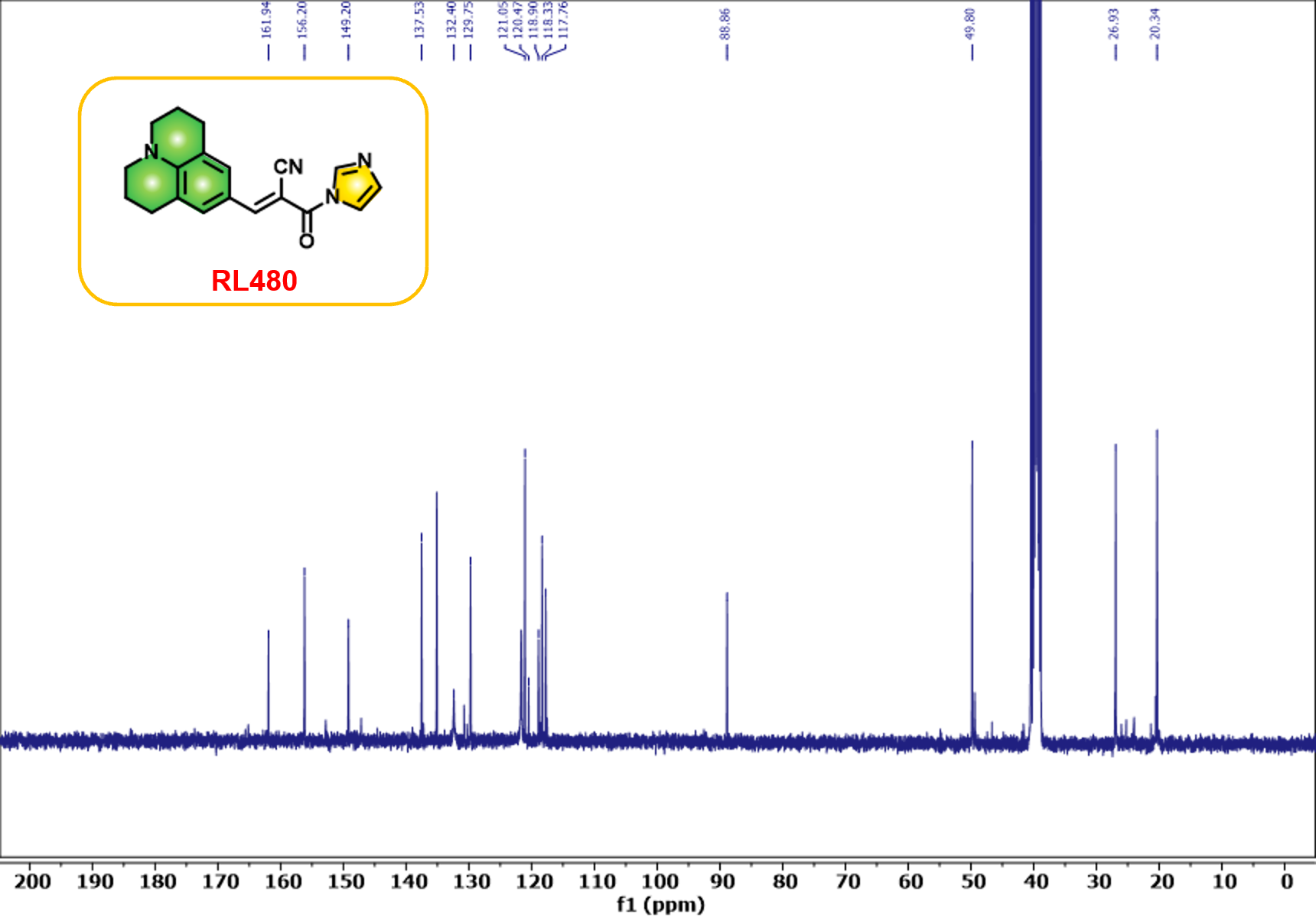

Figure S18. ^13^C NMR spectrum (100 MHz) of RL480 in DMSO-*d*_6_.

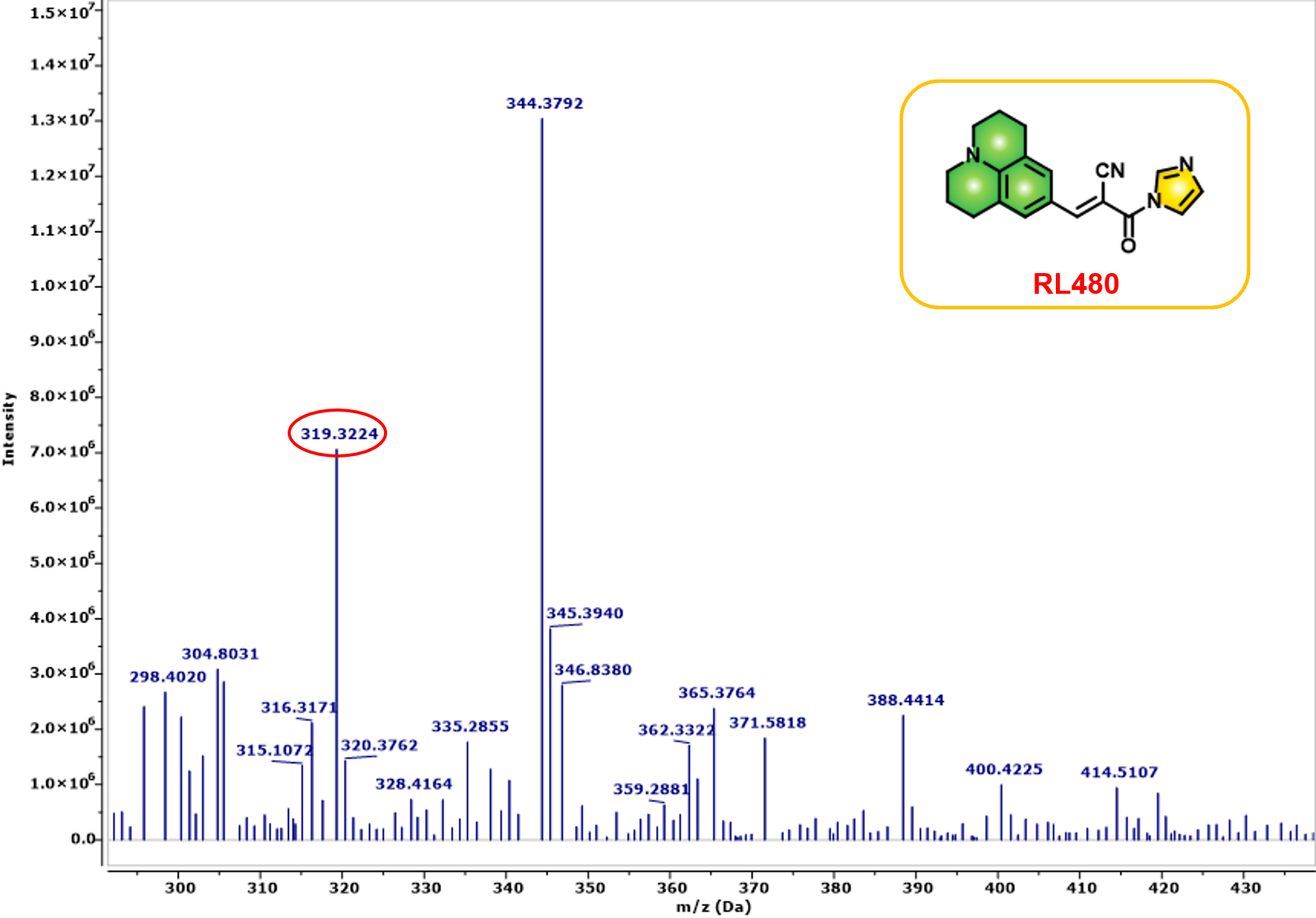

Figure S19. ESI-MS spectrum of RL480.

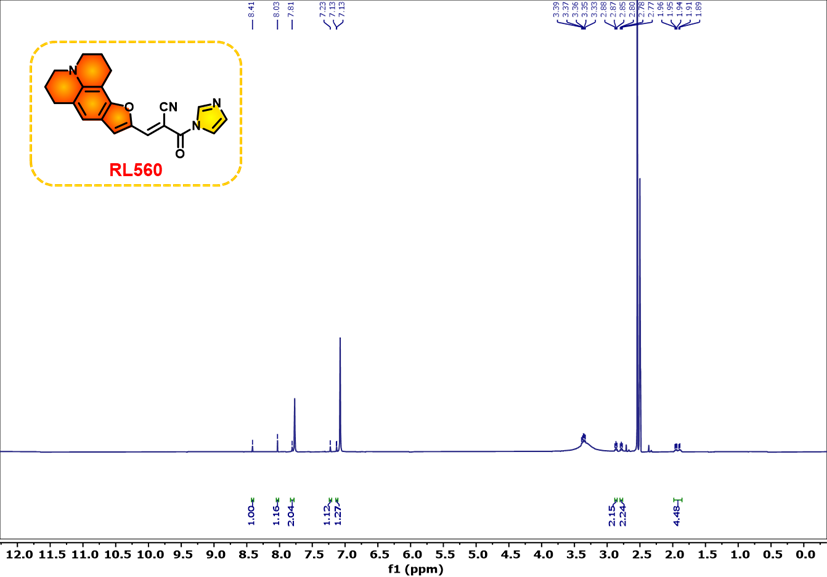

Figure S20. ^1^H NMR spectrum (400 MHz) of RL560 in DMSO-*d*_6_.

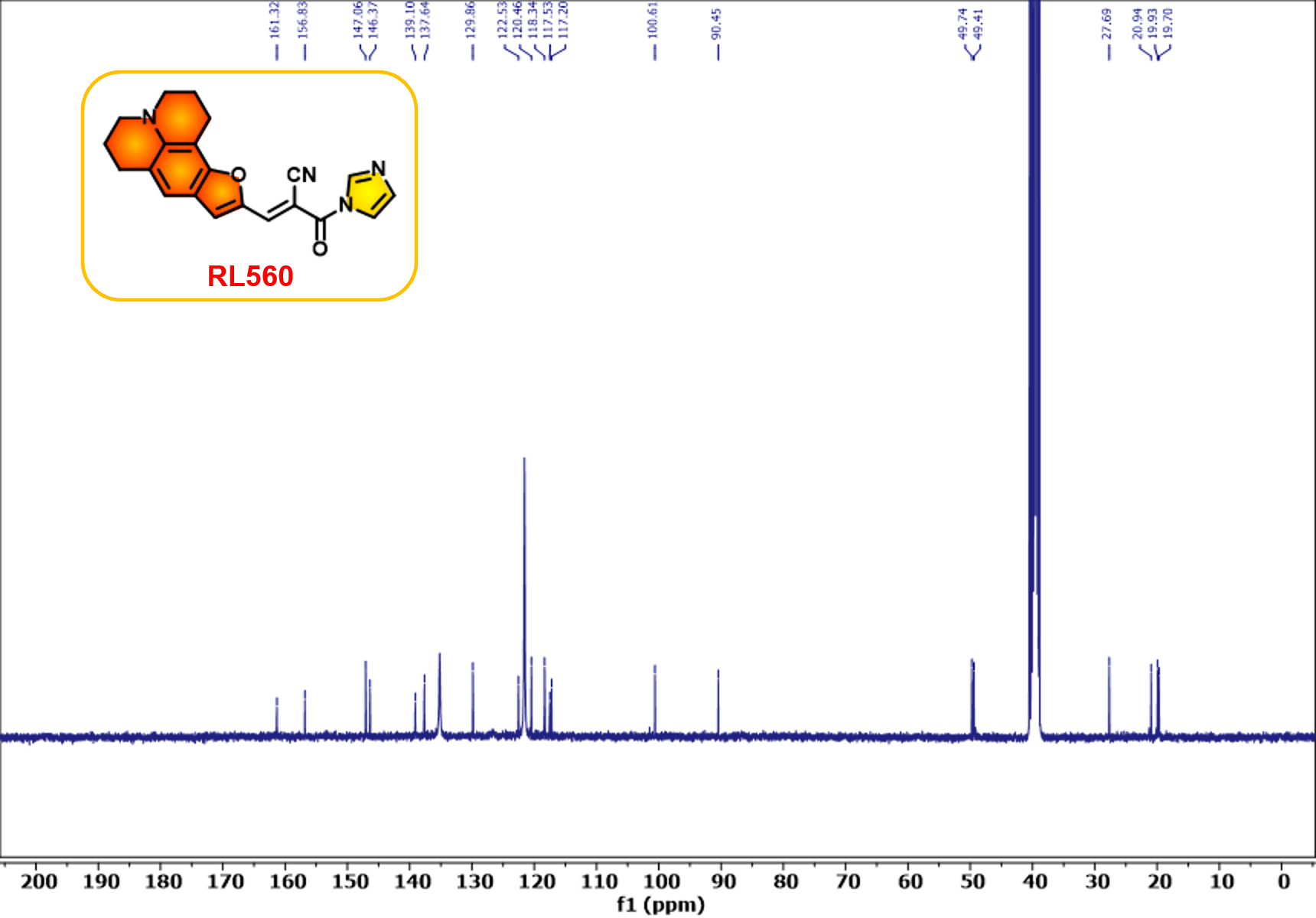

Figure S21. ^13^C NMR spectrum (100 MHz) of RL560 in DMSO-*d*_6_.

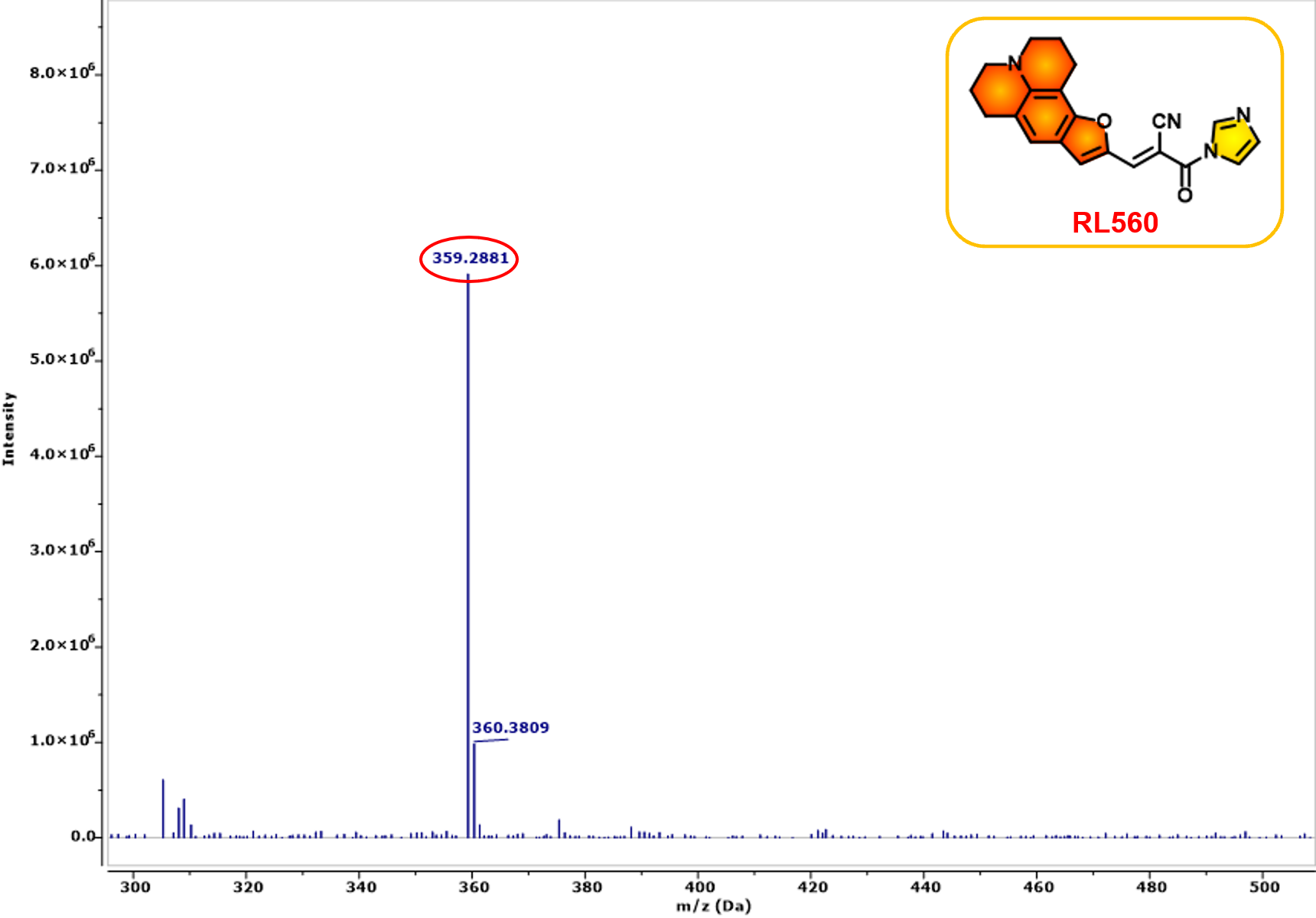

Figure S22. ESI-MS spectrum of RL560.

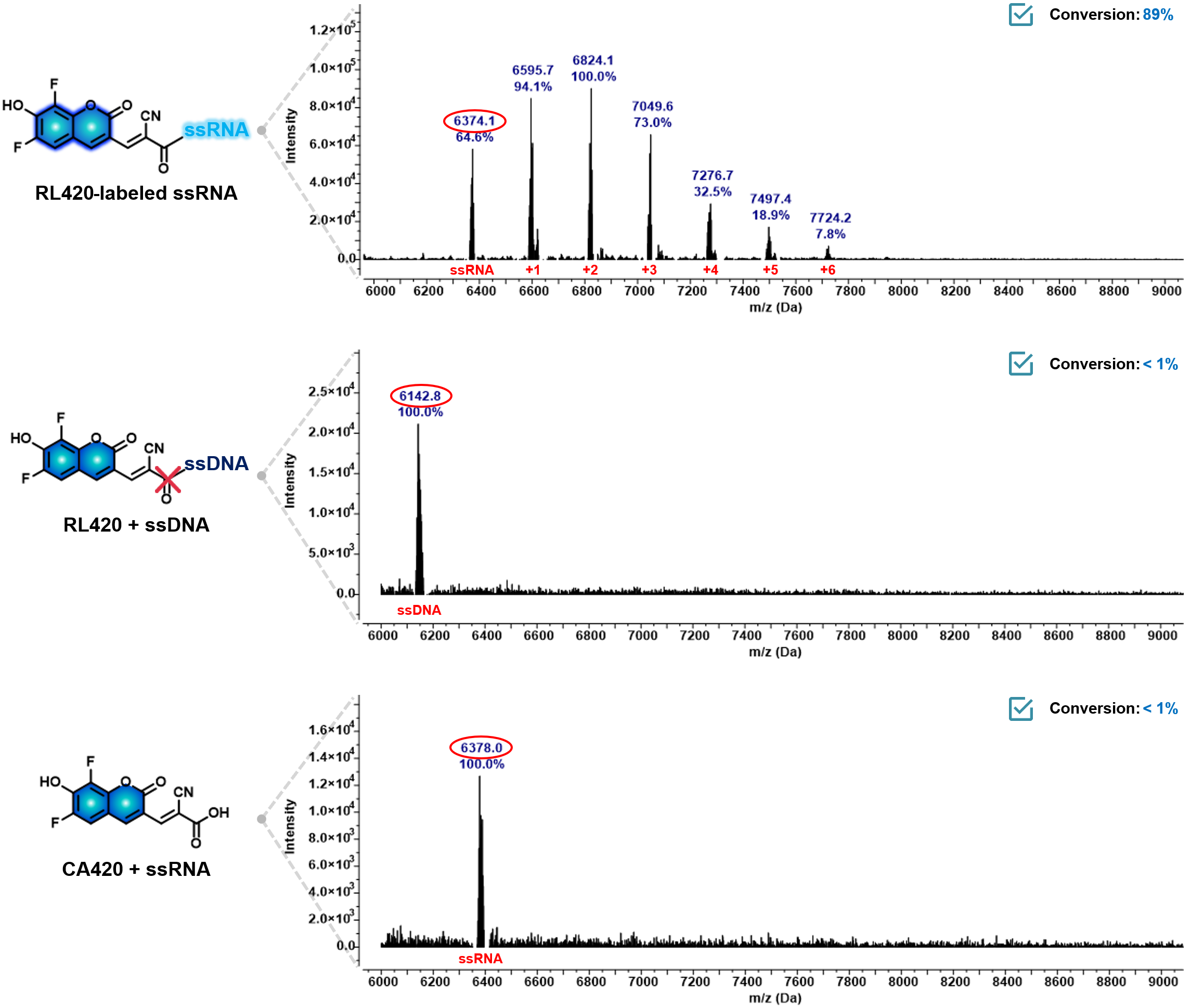

Figure S23. Selective reactivity of RL420 toward ssRNA over ssDNA and structural analog CA420. MALDI–TOF analysis of reactions between RL420 and various nucleic acid substrates. RL420 exhibited efficient covalent labeling with ssRNA (top), as evidenced by mass shifts corresponding to mono- and multi-labeled RNA species (up to +6). In contrast, no labeling was observed with ssDNA (middle), and a structural analog CA420 failed to label ssRNA (bottom). Mass peaks corresponding to unmodified and labeled oligonucleotides are annotated, and relative signal intensities (%) are shown

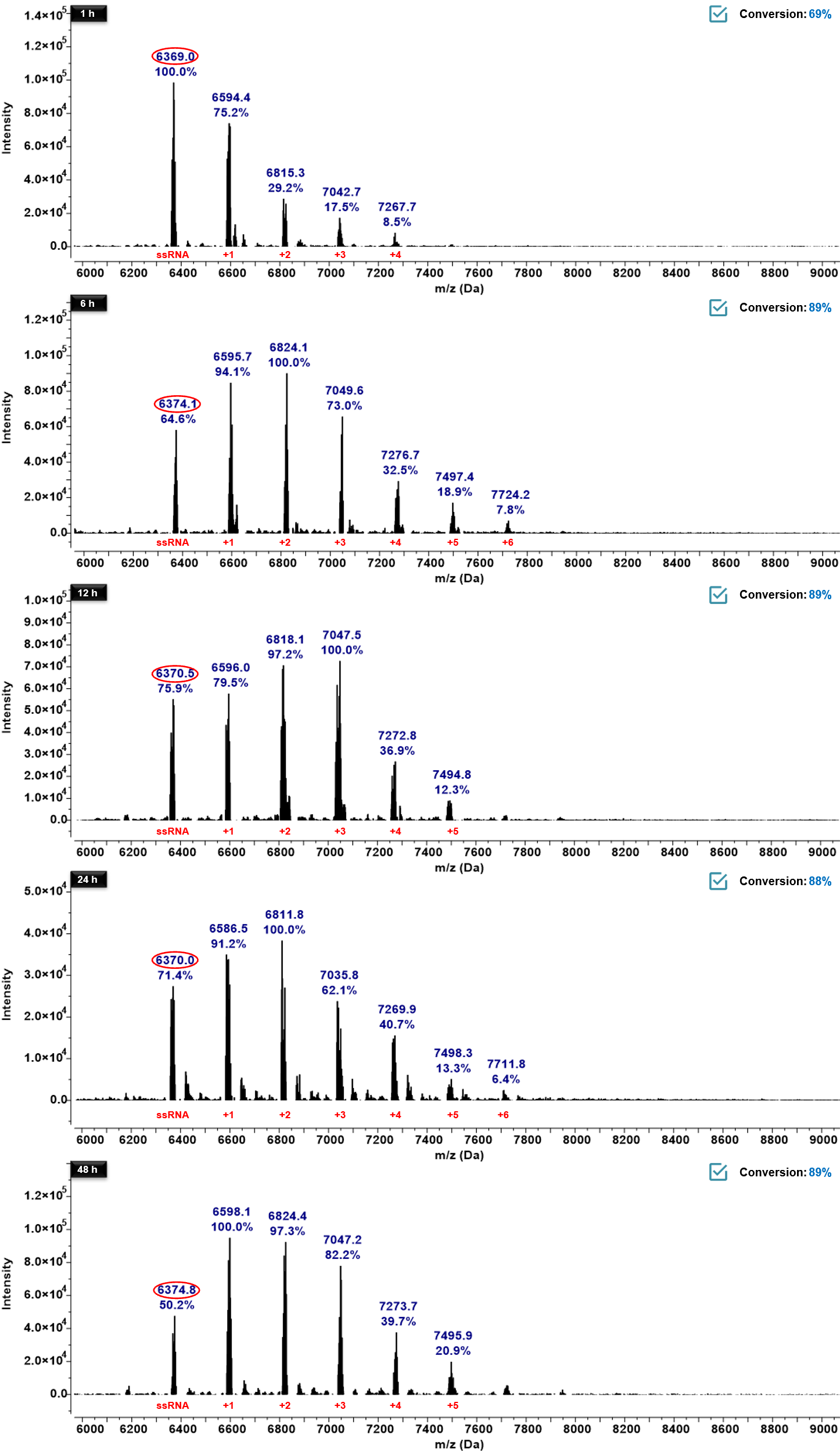

Figure S24. Time-dependent reaction conversions of RL420 with ssRNA monitored by MALDI–TOF. The reaction mixtures of RL420 and ssRNA were incubated at 37 °C and analyzed at 1 h, 6 h, 12 h, 24 h, and 48 h. Mass peaks corresponding to unreacted ssRNA and RL420-labeled products were identified, and their relative signal intensities (%) are indicated. The data demonstrates a time-dependent increase in RNA labeling efficiency by RL420.

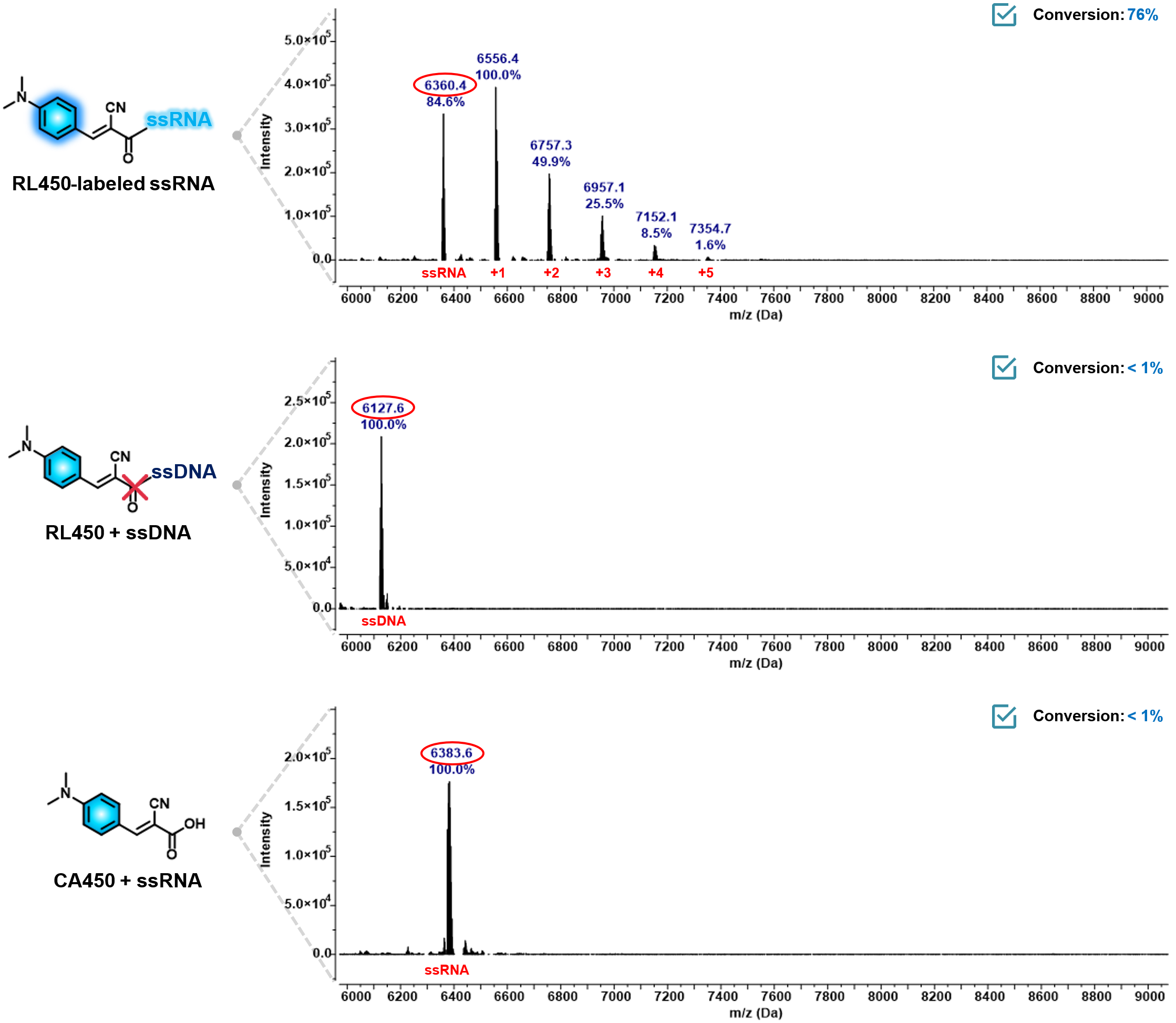

Figure S25. Selective reactivity of RL450 toward ssRNA over ssDNA and structural analog CA450. MALDI–TOF analysis of reactions between RL450 and various nucleic acid substrates. RL450 exhibited efficient covalent labeling with ssRNA (top), as evidenced by mass shifts corresponding to mono- and multi-labeled RNA species (up to +5). In contrast, no labeling was observed with ssDNA (middle), and a structural analog CA450 failed to label ssRNA (bottom). Mass peaks corresponding to unmodified and labeled oligonucleotides are annotated, and relative signal intensities (%) are shown.

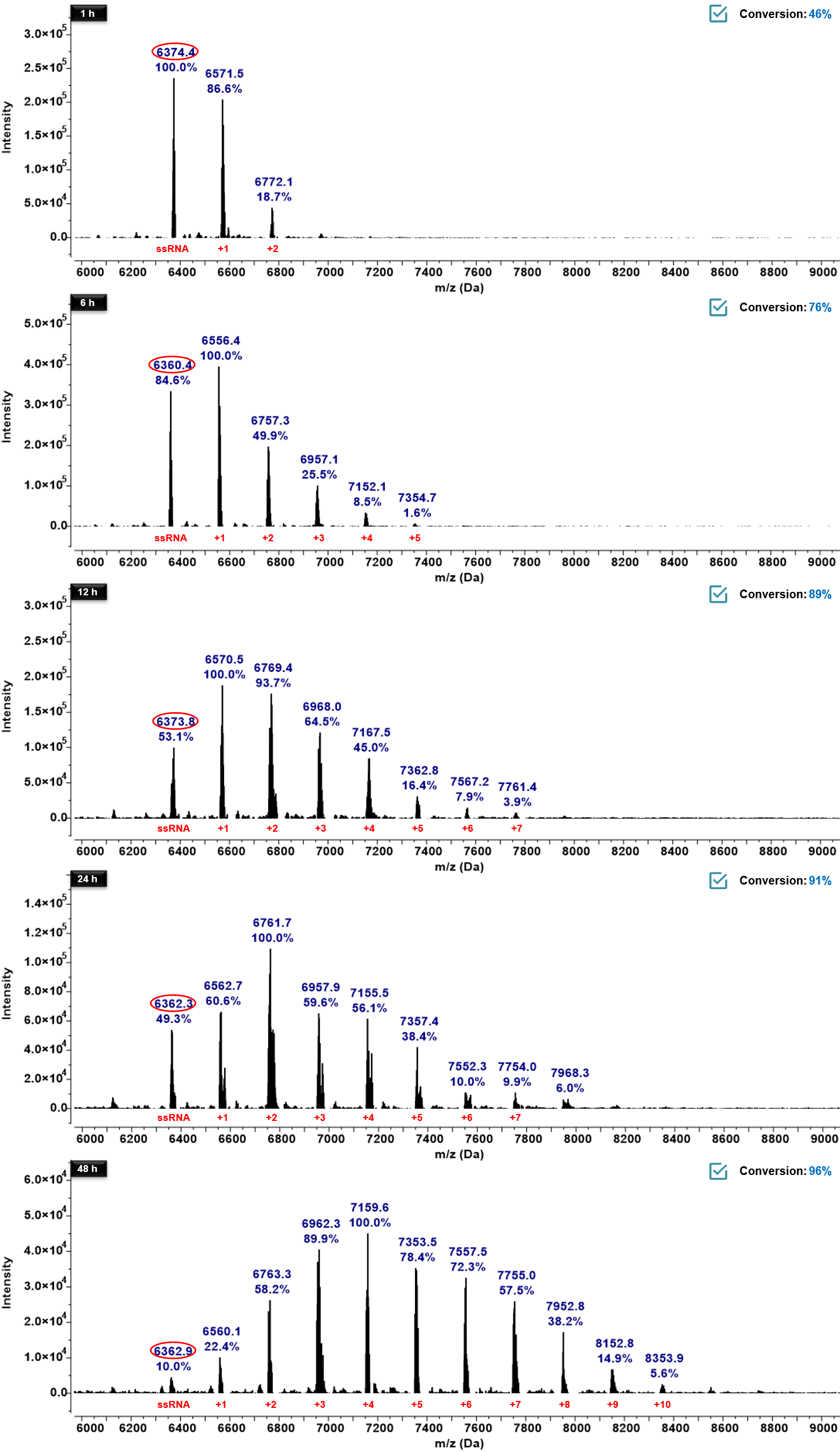

Figure S26. Time-dependent reaction conversions of RL450 with ssRNA monitored by MALDI–TOF. The reaction mixtures of RL450 and ssRNA were incubated at 37 °C and analyzed at 1 h, 6 h, 12 h, 24 h, and 48 h. Mass peaks corresponding to unreacted ssRNA and RL450-labeled products were identified, and their relative signal intensities (%) are indicated. The data demonstrates a time-dependent increase in RNA labeling efficiency by RL450.

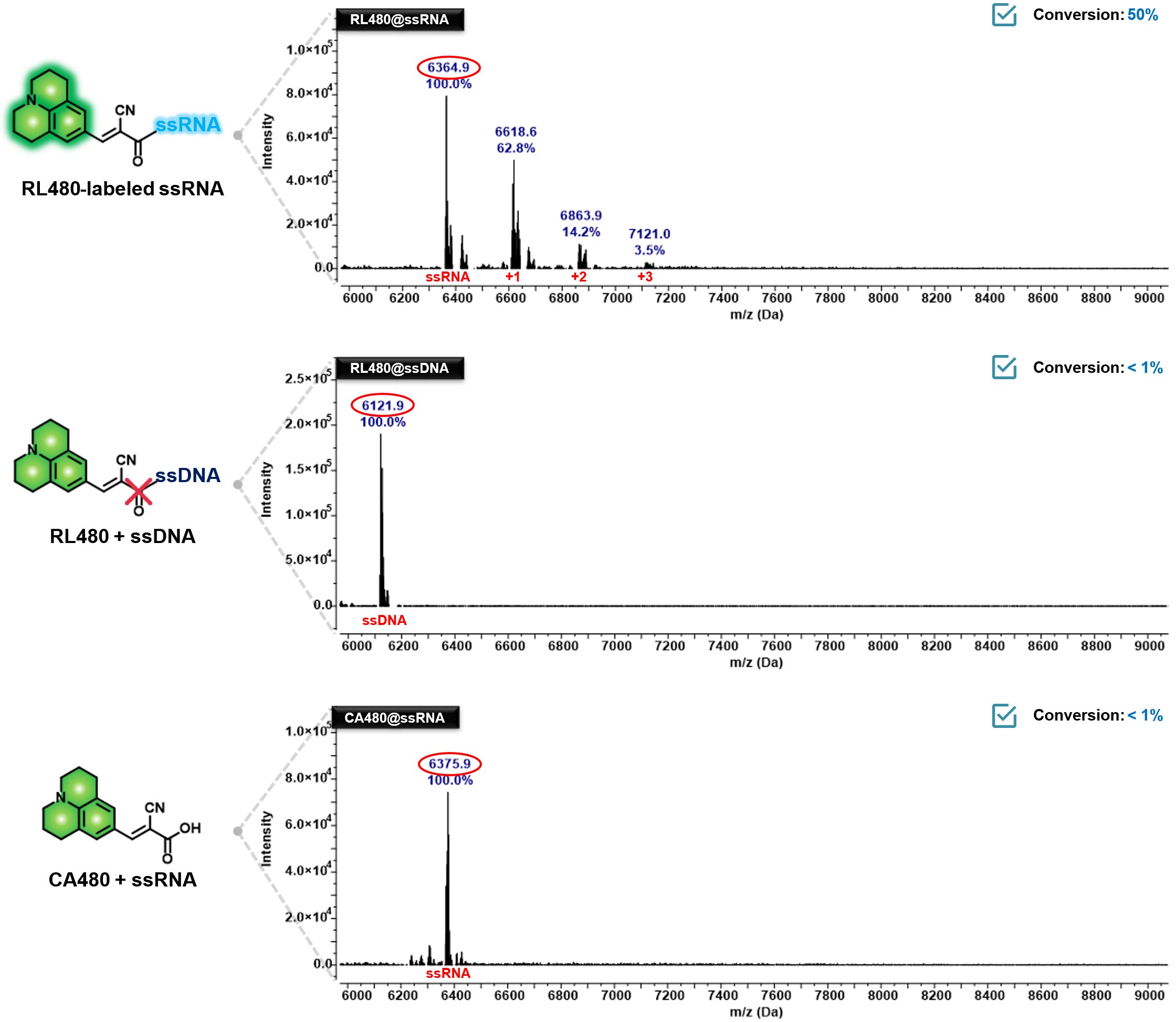

Figure S27. Selective reactivity of RL480 toward ssRNA over ssDNA and structural analog CA480. MALDI–TOF analysis of reactions between RL480 and various nucleic acid substrates. RL480 exhibited efficient covalent labeling with ssRNA (top), as evidenced by mass shifts corresponding to mono- and multi-labeled RNA species (up to +3). In contrast, no labeling was observed with ssDNA (middle), and a structural analog CA480 failed to label ssRNA (bottom). Mass peaks corresponding to unmodified and labeled oligonucleotides are annotated, and relative signal intensities (%) are shown.

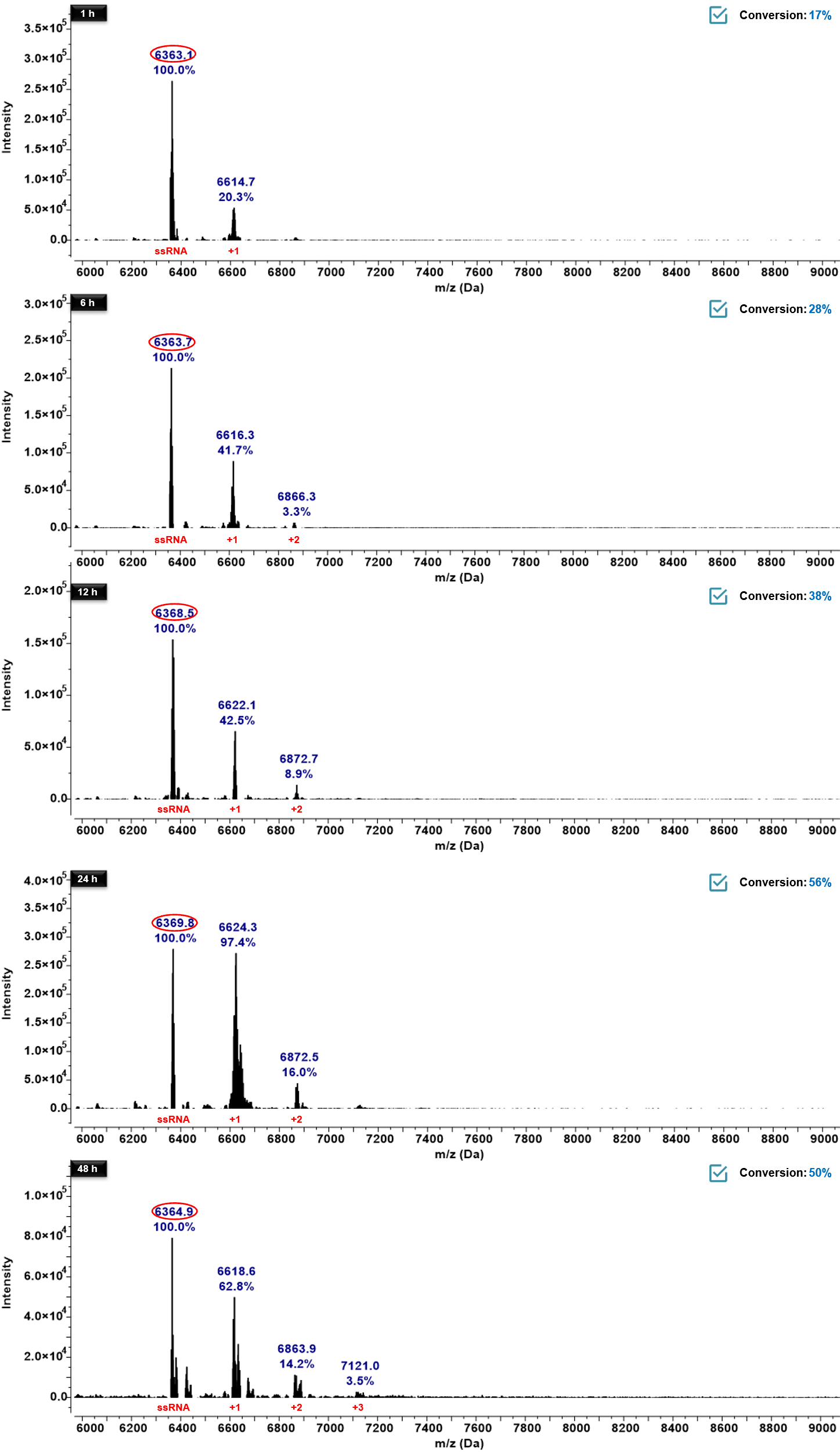

Figure S28. Time-dependent reaction conversions of RL480 with ssRNA monitored by MALDI–TOF. The reaction mixtures of RL480 and ssRNA were incubated at 37 °C and analyzed at 1 h, 6 h, 12 h, 24 h, and 48 h. Mass peaks corresponding to unreacted ssRNA and RL480-labeled products were identified, and their relative signal intensities (%) are indicated. The data demonstrates a time-dependent increase in RNA labeling efficiency by RL480.

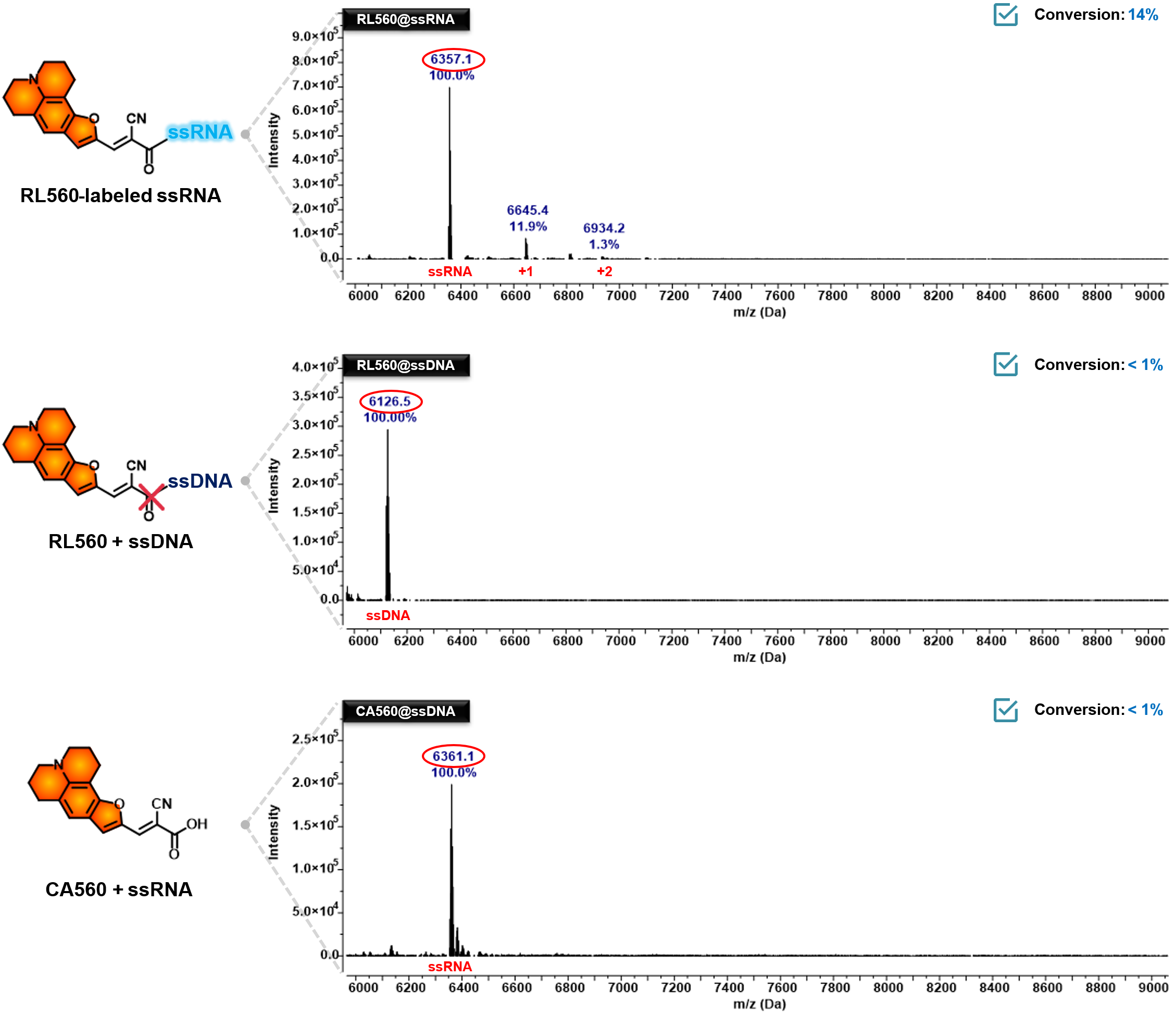

Figure S29. Selective reactivity of RL560 toward ssRNA over ssDNA and structural analog CA560. MALDI–TOF analysis of reactions between RL560 and various nucleic acid substrates. RL560 exhibited efficient covalent labeling with ssRNA (top), as evidenced by mass shifts corresponding to mono- and multi-labeled RNA species (up to +2). In contrast, no labeling was observed with ssDNA (middle), and a structural analog CA560 failed to label ssRNA (bottom). Mass peaks corresponding to unmodified and labeled oligonucleotides are annotated, and relative signal intensities (%) are shown

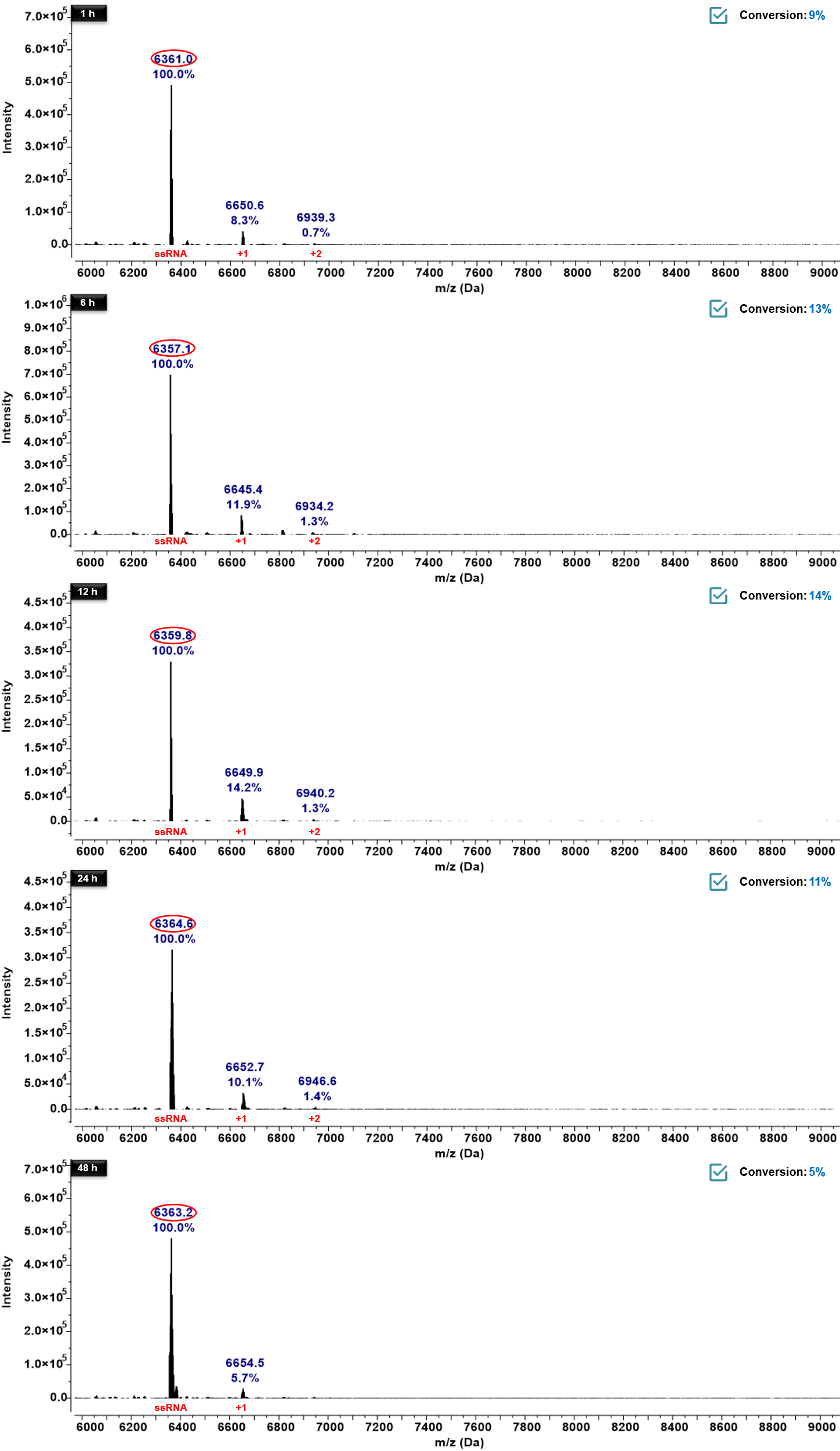

Figure S30. Time-dependent reaction conversions of RL560 with ssRNA monitored by MALDI–TOF. The reaction mixtures of RL560 and ssRNA were incubated at 37 °C and analyzed at 1 h, 6 h, 12 h, 24 h, and 48 h. Mass peaks corresponding to unreacted ssRNA and RL560-labeled products were identified, and their relative signal intensities (%) are indicated. The data demonstrates a time-dependent increase in RNA labeling efficiency by RL560.

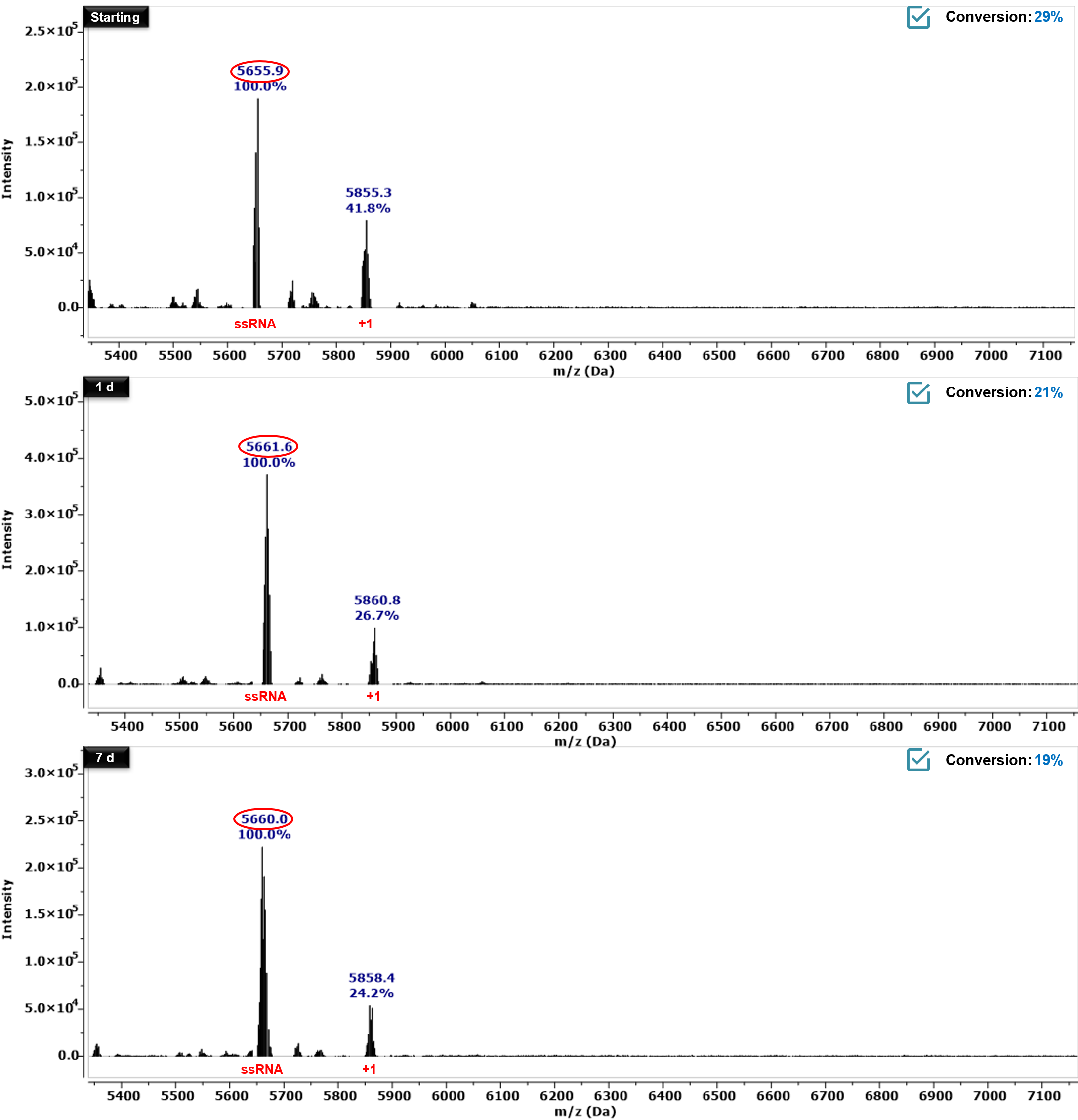

Figure S31. Stability of RL-labeled RNA. RL450-labeled ssRNA was analyzed by MALDI–TOF mass spectrometry at the initial time point (Starting), after 1 d and after 7 d of incubation in nuclease-free water at room temperature. Peaks corresponding to unmodified ssRNA and +1 adduct species are indicated. The relative abundance of each species was calculated from peak intensities, showing minimal loss of covalent adducts over the period, indicating that RL450 labeling remains stable under these conditions.

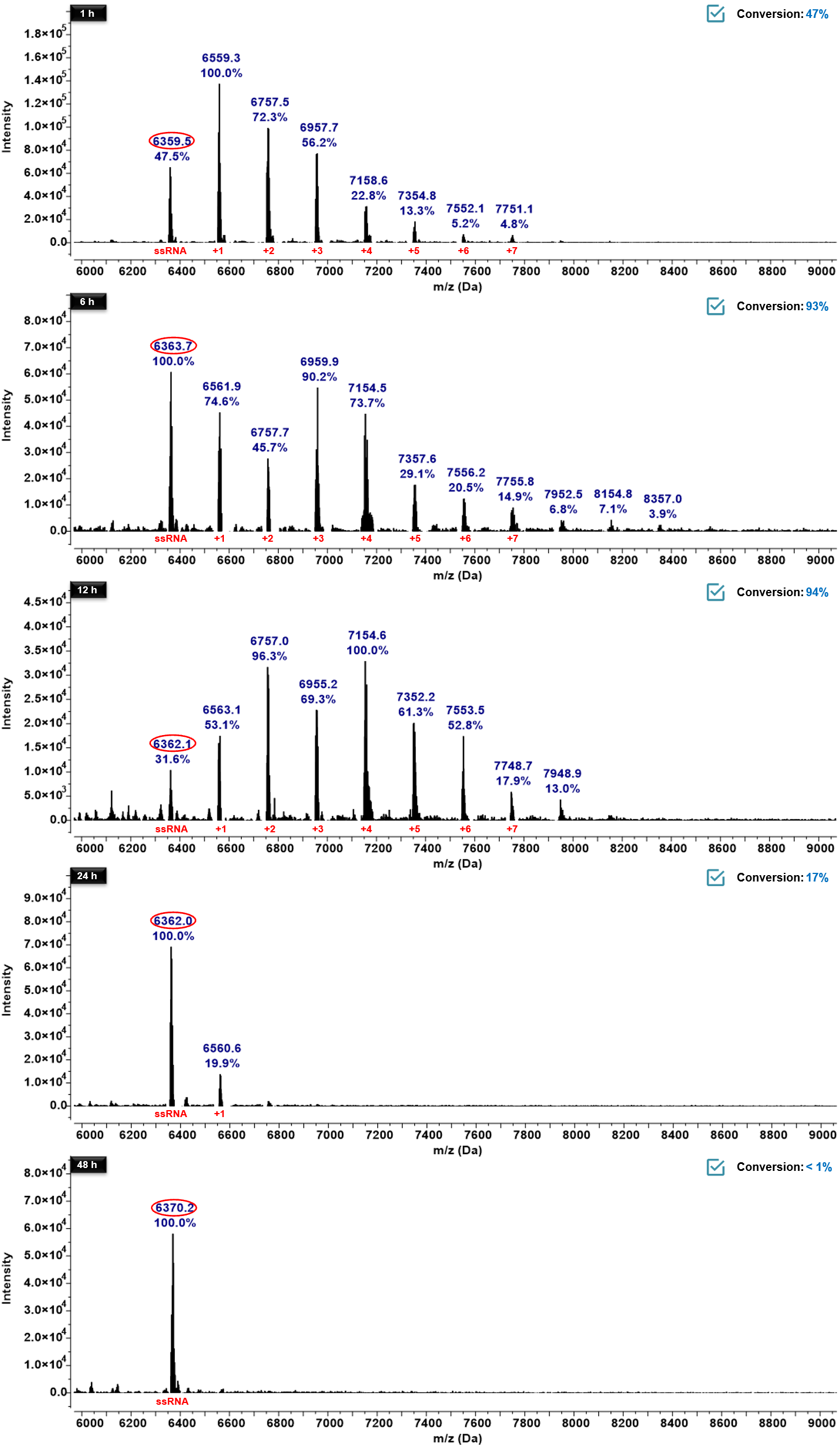

Figure S32. Reversibility of RL-labeled RNA via DMAP treatment. MALDI–TOF mass spectra of RL450-labeled ssRNA at different reaction time points (1, 6, 12, 24 and 48 h), followed by treatment with 100 mM DMAP at 37 °C. Subsequent DMAP treatment results in complete reversal of RNA acylation, regenerating fully unmodified RNA. These results demonstrate the chemical reversibility of the acylation reaction and the potential for temporal control over RNA labeling.

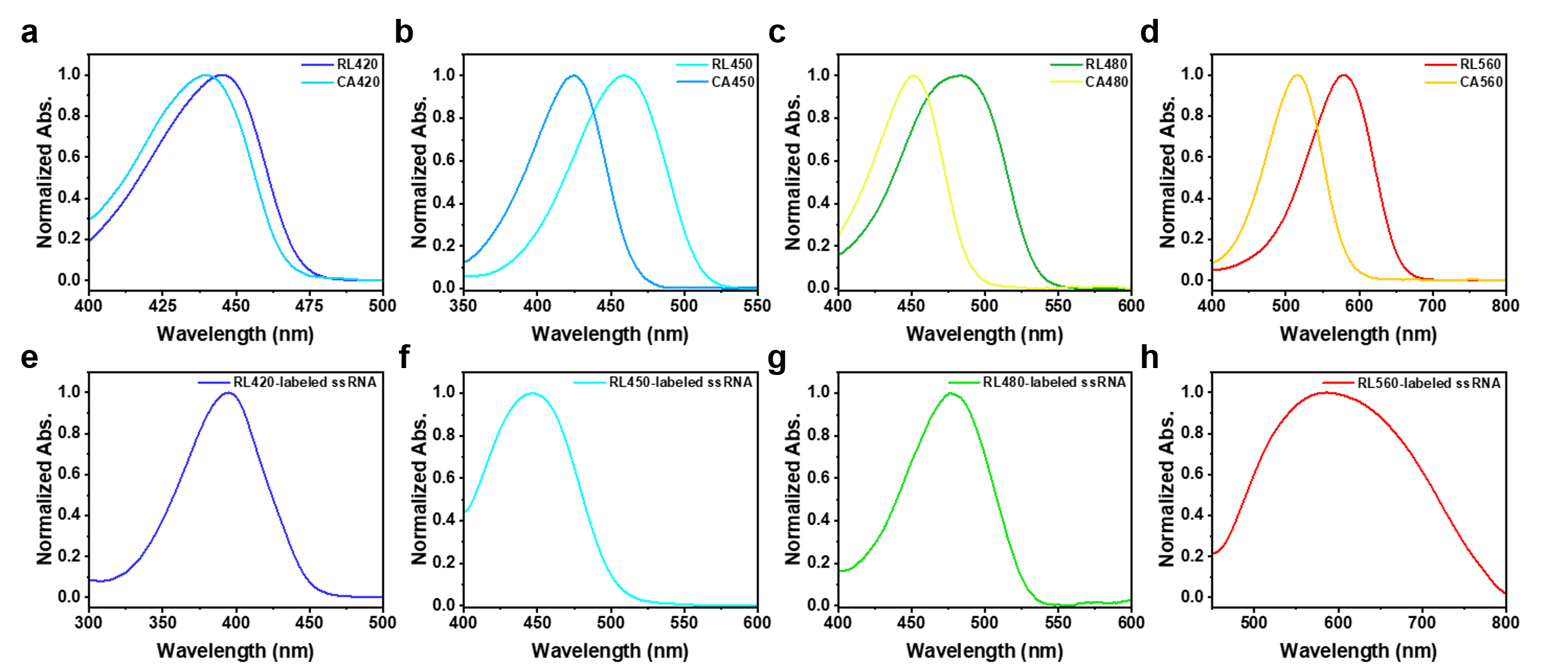

Figure S33. UV/Vis absorption spectra of RL fluorophores, their precursors and RNA-labeled fluorophores. (a–d) Normalized UV/Vis absorption spectra of carboxylic acid precursors and RL fluorophores: RL420/CA420 (a), RL450/CA450 (b), RL480/CA480 (c) and RL560/CA560 (d). Distinct bathochromic shifts in absorption are observed following CDI activation. e–h, Normalized UV/vis spectra of RNA-labeled RL fluorophores: RL420 (e), RL450 (f), RL480 (g) and RL560 (h), highlighting their distinct emission maxima and separation suitable for multicolor RNA imaging applications.

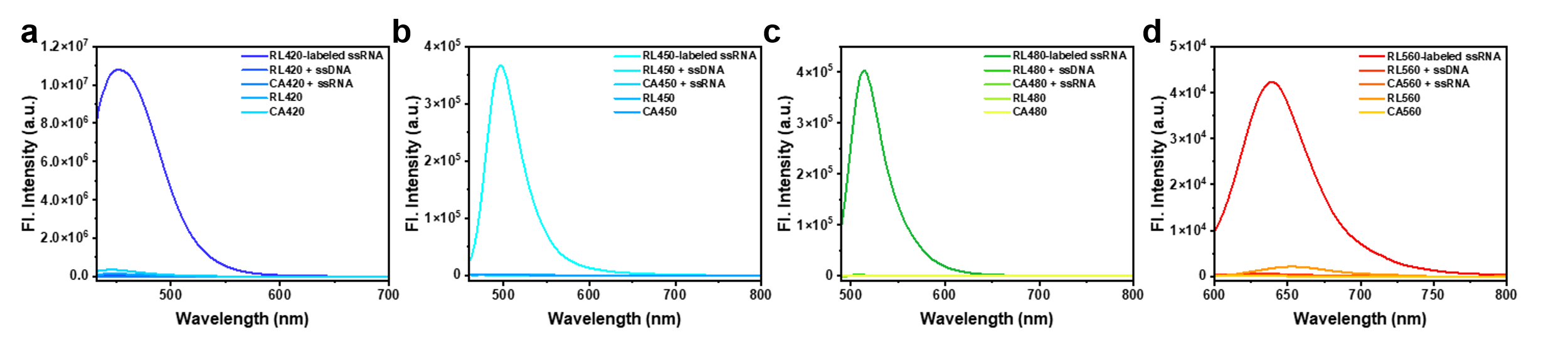

Figure S34. Comparative fluorescence emission of RL fluorophores with ssRNA, ssDNA, CA control, and free dyes. (a–d) Fluorescence emission spectra of RL420 (a), RL450 (b), RL480 (c), and RL560 (d) measured under four different conditions: RL fluorophore incubated with ssRNA, RL fluorophore with ssDNA of identical sequence, corresponding carboxylic acid precursor (CA control) with ssRNA, and free dyes in PBS without nucleic acids. Robust fluorescence turn-on was observed only upon covalent labeling of ssRNA with RL fluorophores, while minimal emission was detected for ssDNA, CA controls, or free dyes, underscoring the covalent and RNA-selective activation of the RL probes.

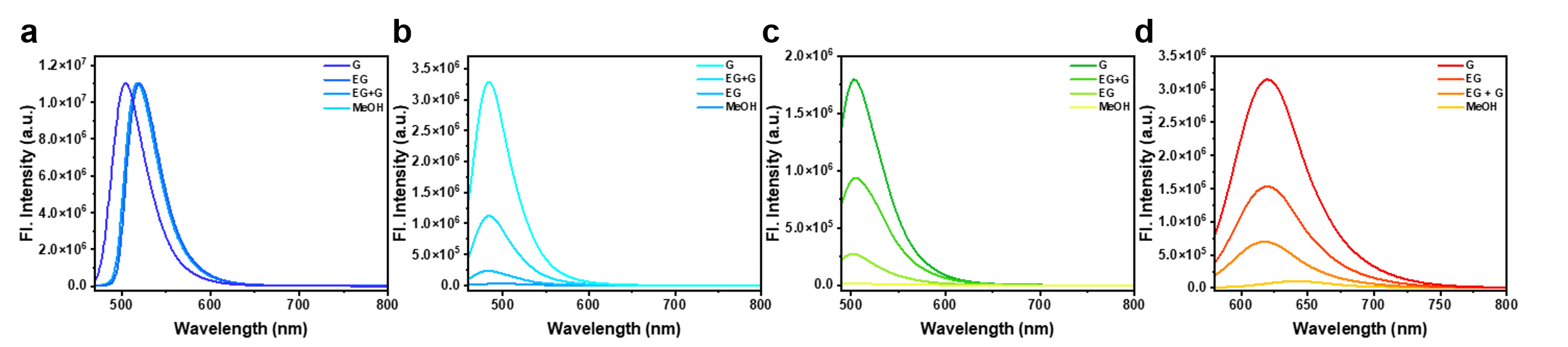

Figure S35. Solvent-dependent fluorescence emission of RL fluorophores under increasing viscosity. (a–d) Fluorescence emission spectra of RL420 (a), RL450 (b), RL480 (c), and RL560 (d) measured in solvents of increasing viscosity: methanol (MeOH), Ethylene glycol (EG), a 1:1 (v/v) mixture of ethylene glycol and glycerol (EG + G), and glycerol (G). For RL450, RL480 and RL560, a increase in fluorescence intensity was observed with increasing viscosity, consistent with a TICT-based fluorescence activation mechanism. In contrast, RL420 exhibited minimal sensitivity to viscosity but showed distinct changes depending on solvent polarity, suggesting that its fluorescence properties are more influenced by polarity than by rotational restriction.

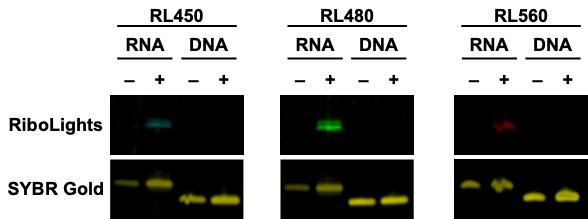

**Figure S36. PAGE gel image after a 1 h incubation with RL fluorophores.** In-gel fluorescence images of ssRNA and ssDNA stained with **RL450**, **RL480**, and **RL560** for 1 h at 37 °C. Fluorescent bands are seen only for labeled ssRNA samples.

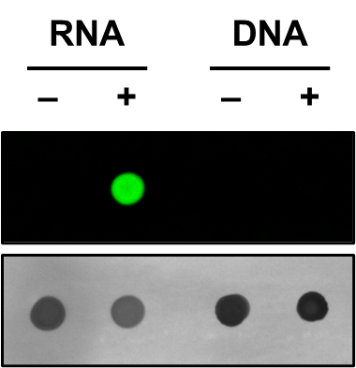

**Figure S37. Comparison of nucleic acid selectivity of RL480 and SYTO RNASelect.** Dot blot analysis of DNA and RNA isolated from HeLa cells treated with or without **RL480**. Fluorescence was detected exclusively in RNA samples from labeled cells, while DNA samples showed no detectable signal.

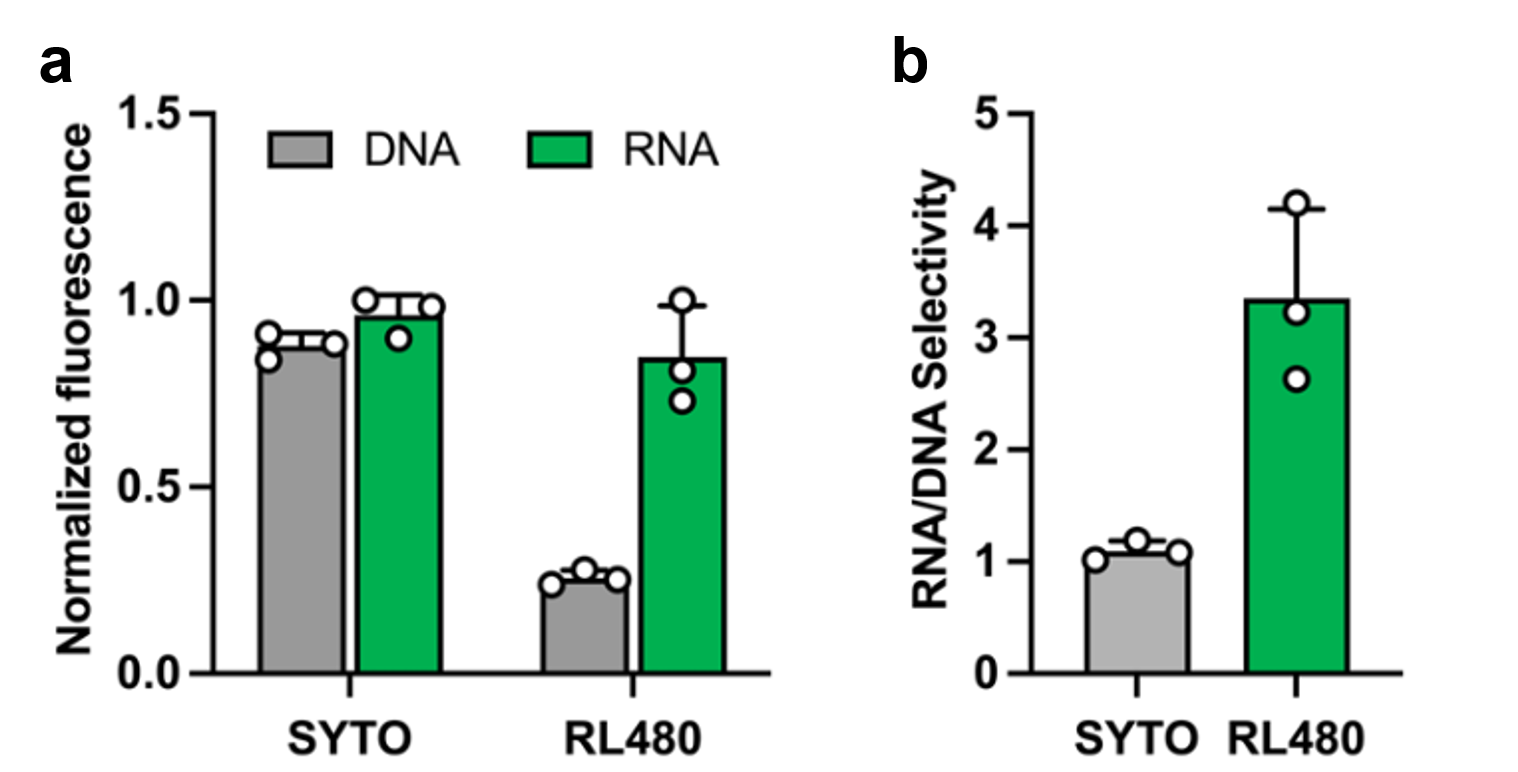

**Figure S38. Comparison of nucleic acid selectivity between RL480 and SYTO RNASelect.** (**a**) Normalized fluorescence intensities of **RL480** and SYTO RNASelect after incubated with cellular RNA or DNA. (**b**) RNA/DNA selectivity calculated from the data in a, showing that **RL480** exhibits approximately 3-fold higher RNA selectivity compared with SYTO RNASelect. Bars represent mean ± *s.d.* from triplicate experiments.

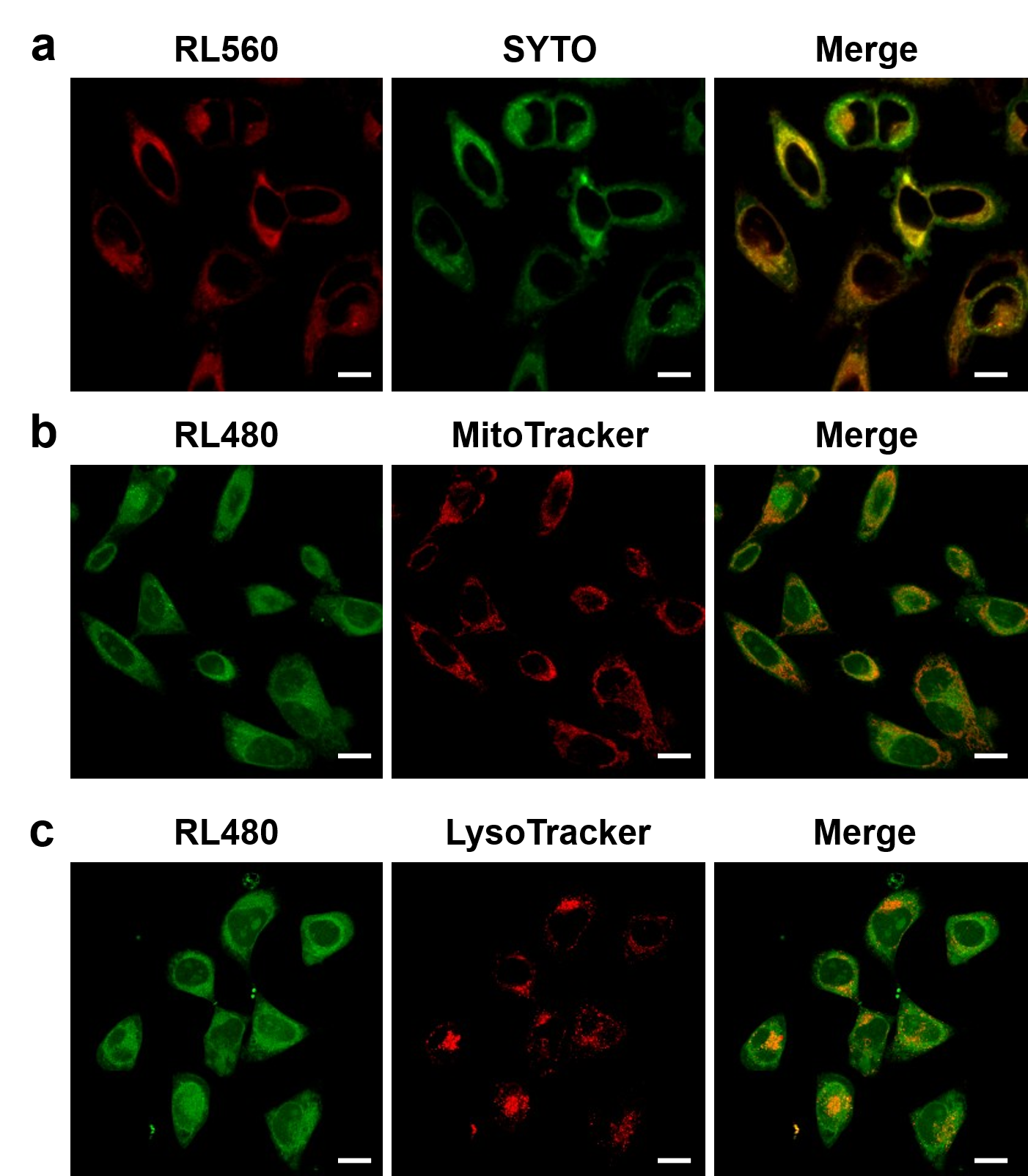

**Figure S39. Colocalization experiments in live cells.** Confocal fluorescence images of live HeLa cells co-stained with **RL560** and SYTO RNASelect (a), **RL480** and MitoTracker Deep Red FM (b), and **RL480** and LysoTracker Deep Red (**c**). **RL560** strongly colocalizes with SYTO RNASelect (Pearson’s R = 0.89) whereas only moderate signal colocalization was observed for **RL480** and MitoTracker/LysoTracker (Pearson’s R = 0.62 and 0.63, respectively). Scale bars: 20 μm.

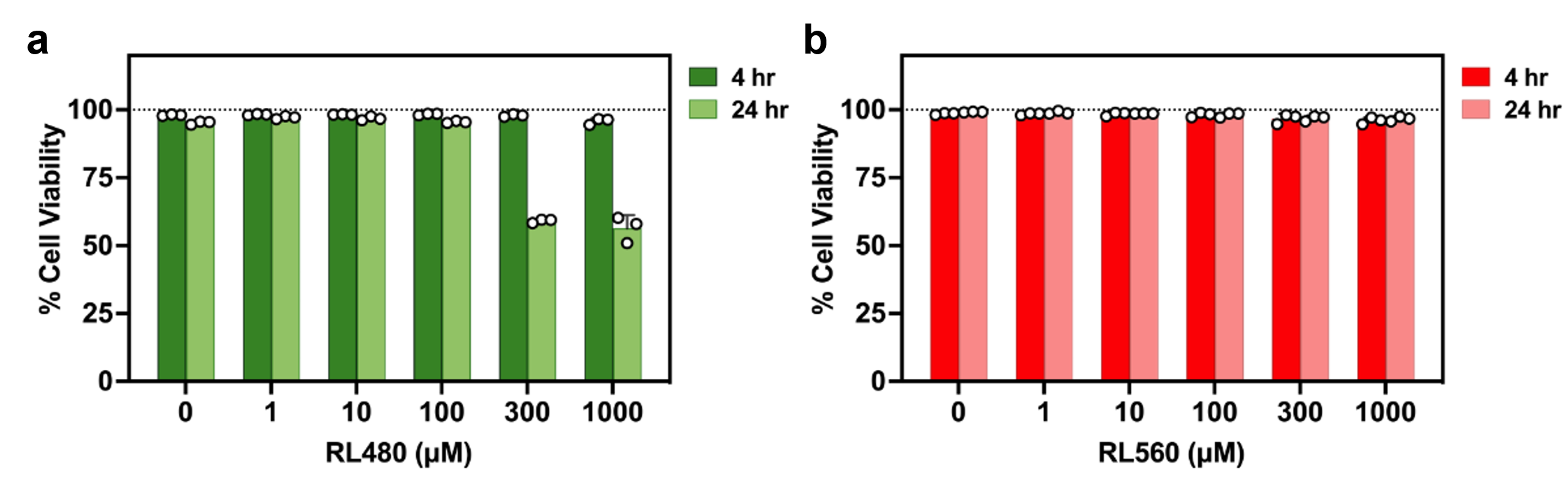

**Figure S40. Cytotoxicity of cell-permeant RL fluorophores.** HeLa cell viability assessed after incubation with 0–1000 μM **RL480** (**a**) and **RL560** (**b**) for 4 to 24 h. No significant toxicity was observed for both dyes at concentrations up to 300 μM and 1000 μM, respectively.

Supplementary Tables

**Table S1.** Comparison with conventional RNA labeling methods and current strategy (**RL** fluorophore).

| **Method** | **Labeling mechanism** | **Sequence dependence** | **Covalent/non-covalent** | **Selectivity (RNA vs. DNA)** | **Main limitations** |
| --- | --- | --- | --- | --- | --- |
| Aptamer-based | Aptamer-flujorogen/protein | Yes | Non-covalent (mainly) | Moderate | Requires RNA modification; signal loss; background |
| Molecular beacons | Hybridization | Yes | Non-covalent | High  (for target) | Target accessibility; slow kinetics; false positives |
| Hybridization probes | Hybridization | Yes | Non-covalent | High  (for target) | Labor-intensive; not ideal for live-cell |
| Enzyme-assisted | Enzyme-mediated modification | Yes | Covalent | High  (at target site) | Needs RNA engineering; substrate scope |
| Non-covalent interaction-based | Intercalation or electrostatic interaction | No | Non-covalent | Moderate | Poor selectivity on RNA over DNA |
| 2′-OH acylation (**RL fluorophore**) | Chemical acylation at RNA 2′-OH | No | Covalent | Very high | Limited selectivity toward the RNA of interest |

**Table S2.** Oligonucleotides used in this work

| RNA Oligonucleotide | |
| --- | --- |
| Name | Sequence (5' to 3') |
| 18-nt ssRNA | AUCCUGCCGACUACGCCA |
| 20-nt ssRNA | ACAAUUAUCCUAUGAGCGGU |
| DNA Oligonucleotide | |
| Name | Sequence (5' to 3') |
| 18-nt ssDNA | AUCCUGCCGACUACGCCA |
| 20-nt ssDNA | ACAATTATCCTATGAGCGGT |
| 60-nt dsDNA (1^st^) | GTCGTAGCTGGATCGTATGCTCAGCGATTTGCGTAGTCGATGAGTGACGTGGATGACTAA |
| 60-nt dsDNA (2^nd^) | ATTGTCATCCACGTCACTCATCGACTACGCAAATCGCTGAGCATACGATCCAGCTCGGAC |
